# Bromodomain and Extraterminal Inhibition Blocks Inflammation-Induced Cardiac Dysfunction and SARS-CoV-2 Infection (Pre-Clinical)

**DOI:** 10.1101/2020.08.23.258574

**Authors:** Richard J Mills, Sean J Humphrey, Patrick RJ Fortuna, Mary Lor, Simon R Foster, Gregory A Quaife-Ryan, Rebecca L. Johnston, Troy Dumenil, Cameron Bishop, Rajeev Ruraraju, Daniel J Rawle, Thuy Le, Wei Zhao, Leo Lee, Charley Mackenzie-Kludas, Neda R Mehdiabadi, Christopher Halliday, Dean Gilham, Li Fu, Stephen J. Nicholls, Jan Johansson, Michael Sweeney, Norman C.W. Wong, Ewelina Kulikowski, Kamil A. Sokolowski, Brian W. C. Tse, Lynn Devilée, Holly K Voges, Liam T Reynolds, Sophie Krumeich, Ellen Mathieson, Dad Abu-Bonsrah, Kathy Karavendzas, Brendan Griffen, Drew Titmarsh, David A Elliott, James McMahon, Andreas Suhrbier, Kanta Subbarao, Enzo R Porrello, Mark J Smyth, Christian R Engwerda, Kelli PA MacDonald, Tobias Bald, David E James, James E Hudson

## Abstract

Cardiac injury and dysfunction occur in COVID-19 patients and increase the risk of mortality. Causes are ill defined, but could be direct cardiac infection and/or inflammation-induced dysfunction. To identify mechanisms and cardio-protective drugs, we use a state-of-the-art pipeline combining human cardiac organoids with phosphoproteomics and single nuclei RNA sequencing. We identify an inflammatory ‘cytokine-storm’, a cocktail of interferon gamma, interleukin 1β and poly(I:C), induced diastolic dysfunction. Bromodomain-containing protein 4 is activated along with a viral response that is consistent in both human cardiac organoids and hearts of SARS-CoV-2 infected K18-hACE2 mice. Bromodomain and extraterminal family inhibitors (BETi) recover dysfunction in hCO and completely prevent cardiac dysfunction and death in a mouse cytokine-storm model. Additionally, BETi decreases transcription of genes in the viral response, decreases ACE2 expression and reduces SARS-CoV-2 infection of cardiomyocytes. Together, BETi, including the FDA breakthrough designated drug apabetalone, are promising candidates to prevent COVID-19 mediated cardiac damage.

## INTRODUCTION

SARS-CoV-2 infection leads to cardiac injury and dysfunction in 20-30% of hospitalized patients (Guo et al., 2020) and higher rates of mortality in patients with pre-existing cardiovascular disease (Shi et al., 2020; Wu and McGoogan, 2020). Inflammatory factors released as part of the ‘cytokine storm’ are thought to play a critical role in cardiac dysfunction in severe COVID-19 patients (Chen et al., 2020). The cardiac sequelae reported in patients with COVID-19, include acute coronary syndromes, cardiomyopathy, acute pulmonary heart disease, arrhythmias and heart failure (Gupta et al., 2020). There have been multiple proposed aetiologies for these, yet clear mechanistic insight is lacking (Gupta et al., 2020). There is a severe inflammatory response in 5% of COVID-19 patients, associated with septic shock (Wu and McGoogan, 2020). This leads to a drop in blood pressure, and approximately 30% of hospitalized patients with COVID-19 require vasopressors to improve blood pressure (Goyal et al., 2020). Furthermore, 68-78% have sustained cardiac dysfunction, primarily right ventricle dysfunction and left ventricular diastolic dysfunction (Puntmann et al., 2020; Szekely et al., 2020).

In severe disease, inflammation associated with a cytokine storm can cause cardiac dysfunction and pathology. COVID-19 induces a cytokine storm of similar magnitude to that induced by CAR-T cell-associated cytokine storms (Del Valle et al., 2020). Additionally, severe COVID-19 is associated with sepsis and bacterial products in the serum (Arunachalam et al., 2020), which are known drivers of cardiac pathology and dysfunction. In the absence of infection, well known inflammatory mediators such as TNF are associated with heart failure and have been demonstrated to induce systolic dysfunction (Feldman et al., 2000). Thus, inflammation is a likely mediator of cardiac injury and dysfunction in COVID-19 patients, which may result in further pathological consequences including inadequate organ perfusion and immune cell infiltration, further exacerbating disease. Thus, preventing cytokine-induced cardiac dysfunction may limit severe outcomes in COVID-19 patients. However, targeted treatment strategies, particularly in severe infections such as COVID-19, are currently lacking.

Several anti-inflammatory agents have shown clinical benefit for the acute management of COVID-19. Dexamethasone improved 28-day mortality in COVID-19 patients receiving invasive mechanical ventilation or oxygen at randomization (Horby et al., 2020). Additionally, Janus kinase (JAK)/signal transducer and activator of transcription (STAT) (ruxolitinib and baricitinib) and IL-6R inhibitors (tocilizumab and sarilumab) are currently in COVID-19 clinical trials. However, systemic immunosuppression may impede viral clearance thus potentially exacerbating disease (Mangalmurti and Hunter, 2020). To circumvent this, we aimed to identify cardiac-specific inflammatory targets that trigger cardiac dysfunction in response to the cytokine storm, reasoning that these might provide a safe and effective therapeutic option.

Here, we utilize multi-cellular human pluripotent stem cell (hPSC)-derived cardiac organoids (hCO) combined with phosphoproteomics and single nuclei RNA-sequencing (RNA-seq) to identify therapeutic targets and treatments for cardiac dysfunction. We recently adapted our hCO system (Mills et al., 2019; Mills et al., 2017) to include co-culture with endothelial cells that form enhanced branched endothelial structures surrounded by pericytes (**Figure S1**, Voges et al., In Preparation). This protocol results in a complex mixture of self-organising cells including epicardial, fibroblasts/pericytes, endothelial cells and cardiomyocytes. This was combined together with an optimized culture environment that reflects a maturation stage; mimicking the postnatal metabolic environment (Mills et al., 2019; Mills et al., 2017) followed by reversion to a more adult metabolic substrate provision (see **Methods**). This platform enabled rapid screening of cytokine combinations that recapitulate the COVID-19-induced cytokine storm (Mangalmurti and Hunter, 2020) and cardiac dysfunction, with the subsequent application of -omic assays and drug screening.

## RESULTS

### Cytokine-induced cardiac dysfunction

We began by examining the effects of a range of pro-inflammatory cytokines elevated in COVID-19 patients (Huang et al., 2020) on cardiac function in our hCO (Mills et al., 2017). Inflammatory molecules tested were likely candidates in COVID-19 including: TNF, IL-1β, IFN-γ, IL-6, IL-17A, and G-CSF, as well as pathogen-associated molecular patterns including poly(I:C) to mimic dsRNA, and lipopolysaccharide (LPS) to mimic TLR4 activation and septic responses. Using our RNA-seq data (Mills et al., 2017), we identified that the receptor genes *IL1R1*, *TNFRSF1A*, *TNFRSF1B*, *IFIH1*, *MYD88*, *IL6ST*, *IFNAR1*, *IL6R*, *TMEM173*, *IL17RA*, *IL17RB*, *IL17RC*, *IL17RD*, *IL17RE*, *IFNGR1*, *TLR3*, and *TLR4* were expressed at similar or higher abundance in our hCO compared to adult human heart (**Figure S2A**). In adult mouse hearts many of these are enriched in non-myocyte populations (Quaife-Ryan et al., 2017) (**Figure S2B**). We used single nuclei RNA sequencing (snRNA-seq) to assess cell specificity in our enhanced hCO (Voges et al., In Revision). Mapping to human heart snRNA-seq (Tucker et al., 2020) revealed the presence of pro-epicardial/epicardial cells, fibroblasts, activated fibroblasts/pericytes and cardiomyocytes (**Figure S2C,D**). Some cardiomyocytes were fetal-like, however there was a distinct sub-cluster that mapped adjacent to adult ventricular cardiomyocytes from human hearts (Gilsbach et al., 2018) (**Figure S2E**). The cytokine/pro-inflammatory receptors were expressed across different cell types, but were enriched and more highly expressed in epicardial cells and fibroblasts (**Figure S2F**) (Mills et al., 2017; Voges et al., 2017). We screened inflammatory factors in all pair-wise combinations in hCOs with multiple functional measurements including contractile force, rate, activation kinetics and relaxation kinetics (Mills et al., 2019; Mills et al., 2017) (**Figure 1A**). TNF caused a reduction in force, while IL-1β, IFN-γ, poly(I:C) and LPS caused diastolic dysfunction characterized by a preserved contractile force but prolonged time from peak to 50% relaxation (**Figure S3**). A secondary full-factorial screen of TNF, IFN-γ, IL-1β, and poly(I:C), once again revealed that TNF induced systolic dysfunction (**Figure 1B,D**) with a EC_50_ of 1 ng/mL at 48 hours (**Figure S4A**). A combination of IL-1β, IFN-γ and poly(I:C) induced diastolic dysfunction (**Figure 1C,E**), however also decreased the beating rate which may influence the kinetics of contraction (**Figure S5, Supplementary Video 1,2**). Changes in rate were not responsible for increased relaxation time, as hCO paced at 1 Hz retained the severe diastolic dysfunction phenotype (**Figure 1F, Supplementary Video 3,4**). Individually, IFN-γ and IL-1β caused concentration-dependent diastolic dysfunction with an EC_50_ of 0.8 ng/mL at 48 hours and 3 ng/mL at 24 hours, respectively, whereas poly(I:C) alone did not induce dysfunction (**Figure S4B to D**). These results were confirmed in an independent hPSC line, where the combination of IFN-γ, IL-1β and poly(I:C) induced the most consistent, robust diastolic dysfunction (**Figure S6A to E**). Taken together, TNF induces systolic dysfunction consistent with previous *in vitro* (Vasudevan et al., 2013) and *in vivo* (Kubota et al., 1997) studies, and the combination of IFN-γ, IL-1β and poly(I:C) induces severe diastolic dysfunction in hCO. The dominant factor identified that causes diastolic dysfunction, IFN-γ (**Figure S6C**), is generally elevated in heart failure patients, but its role in heart failure is contradictory in animal models with both detrimental and beneficial effects reported (Levick and Goldspink, 2014).

**Figure 1:**
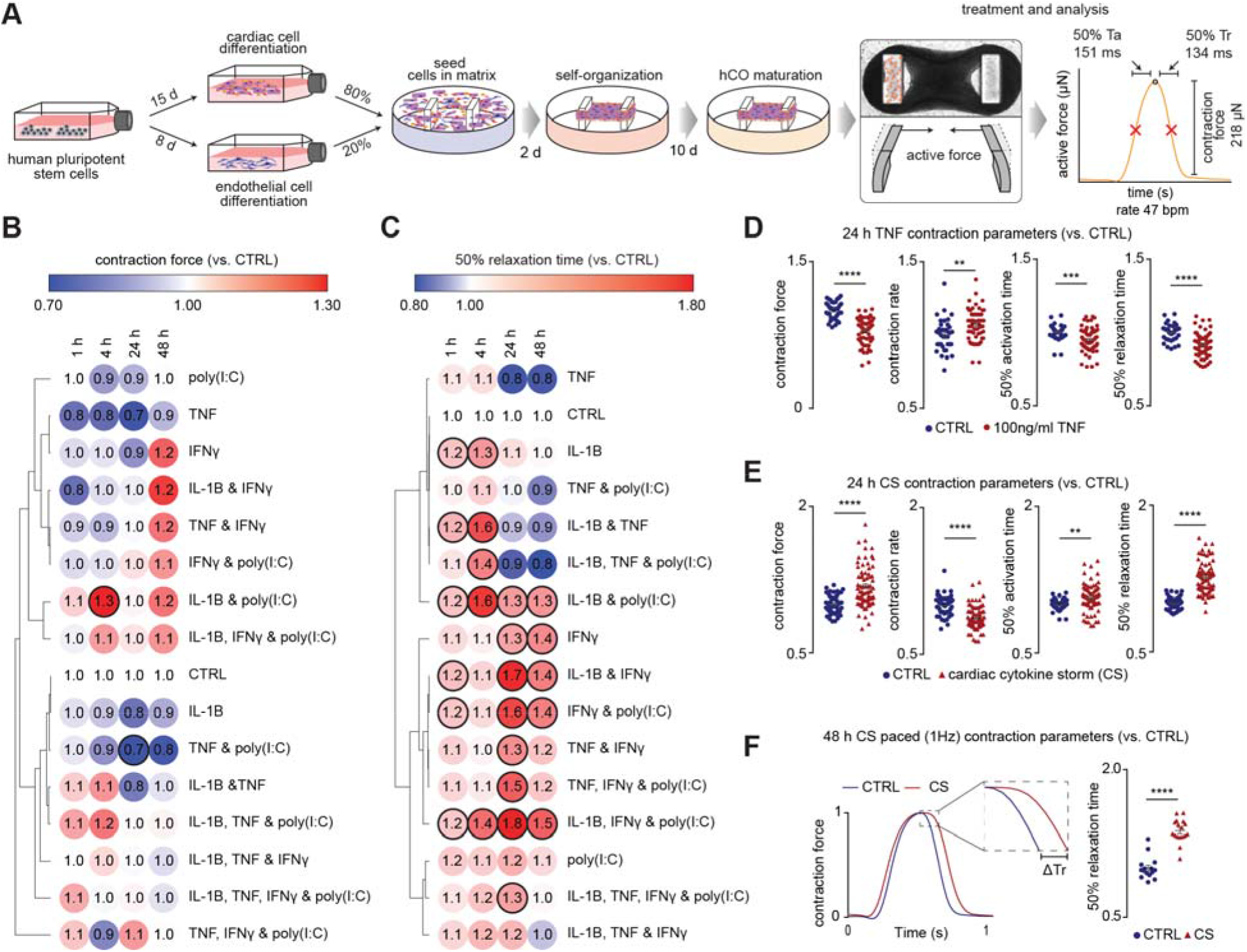
Identification of pro-inflammatory factors driving cardiac dysfunction. A) Schematic of experimental pipeline. Values for overall functional parameters are the mean of n = 1,100 hCO from 9 experiments. B) Impact of inflammatory modulators on force (systolic function). Bold outline indicates p <0.05 using a one-way ANOVA with Dunnett’s multiple comparisons test comparing each condition to CTRL at its’ time point. n= 3-5 hCOs per condition from 1 experiment. hPSC cardiac cells- AA, Endothelial cells- RM3.5. C) Impact of inflammatory modulators on time to 50% relaxation (diastolic function). Bold outline indicates p <0.05 using a one-way ANOVA with Dunnett’s multiple comparisons test comparing each condition to CTRL at the respective time points. n= 3-5 hCOs per condition from 1 experiment. hPSC cardiac cells- AA, Endothelial cells- RM3.5. D) TNF causes systolic dysfunction. n = 37 and 63 hCOs for CTRL and TNF conditions, respectively from 6 experiments. hPSC cardiac cells- HES3, Endothelial cells- RM3.5 or CC. ** p<0.01, *** p<0.001, **** p<0.0001, using Student’s t-test. E) CS causes diastolic dysfunction n = 49 and 73 hCOs for CTRL and CS conditions, respectively from 6 experiments. hPSC cardiac cells- HES3, Endothelial cells- RM3.5 or CC. ** p<0.01, **** p<0.0001, using Student’s t-test. F) Representative force curve of hCO under CTRL and CS conditions (1 Hz) 48 h after treatment. Relaxation of CTRL and CS under paced conditions (1 Hz) 48 h after treatment. n = 15 and 17 hCOs per condition, respectively, from 3 experiments. hPSC cardiac cells- HES3, Endothelial cells- RM3.5 or CC. **** p<0.0001, using Student’s t-test. CTRL – control. CS – cardiac cytokine storm conditions: IL-1β, IFN-γ and poly(I:C). Inflammatory screen in B-E repeated in an additional cell line in **Figure S6**. Ta – time from 50% activation to peak, Tr – time from peak to 50% relaxation.

### Mechanisms driving cardiac cytokine storm-induced dysfunction

The most common types of cardiac dysfunction observed in hospitalized COVID-19 patients are right ventricular dysfunction or left ventricular diastolic dysfunction (Szekely et al., 2020). Therefore, we chose to interrogate the mechanism of diastolic dysfunction induced by IFN-γ, IL-1β and poly(I:C), from here on referred to as ‘cardiac cytokine storm’ (CS). Protein phosphorylation is intimately linked with all biological functions (Needham et al., 2019), and thus reasoned that measuring the global phosphoproteome in hCO would provide an accurate fingerprint of the mechanistic targets driving dysfunction caused by CS. Leveraging the latest developments in our phosphoproteomics technology (Humphrey et al., 2015; Humphrey et al., 2018), we analysed CS-induced phosphorylation events in 20 pooled organoids, yielding ~100 μg of protein each. We identified over 7,000 phosphosites in each sample, accurately pinpointing 7,927 phosphorylation sites to a single amino acid residue on ~3,000 different phosphoproteins from single-run measurements (**Figure 2A**, **Figure S7**). Preliminary studies using TNF identified several known biological effects of this cytokine including decreased phosphorylation of protein kinase A and increased phosphorylation of beta-adrenergic receptor kinase 1 (also known as GRK2), supporting our approach (data not shown). Applying this technology to the CS treatment revealed 91 phosphosites that were consistently elevated across three biological replicates (**Figure 2B**). These sites were enriched for terms relating to proliferation, with transcription factors over-represented with 35 sites found on transcription factors or chromatin-binding proteins and 13 associated with the biological process term ‘cell proliferation’ (FDR < 0.05, Fisher’s exact test). Among these was phosphorylation of signal transducer and activator of transcription 1 (STAT1) S727 (median 13.9 fold), as well as two sites on BRD4 (Bromodomain-containing protein 4) S469 and S1083 (median 7.4 and 12.3 fold respectively) (**Figure 2B,C**). In light of the availability of specific small molecule inhibitors for each of these targets or their upstream regulators we focused on these proteins in subsequent functional assays. The cytokine receptor enrichment in non-myocytes (**Figure S2**) and broad expression of key mediators such as STAT1 and BRD4 (**Figure 3H** and **Figure 4F**) suggests a multi-cellular response mediates cardiac dysfunction.

**Figure 2:**
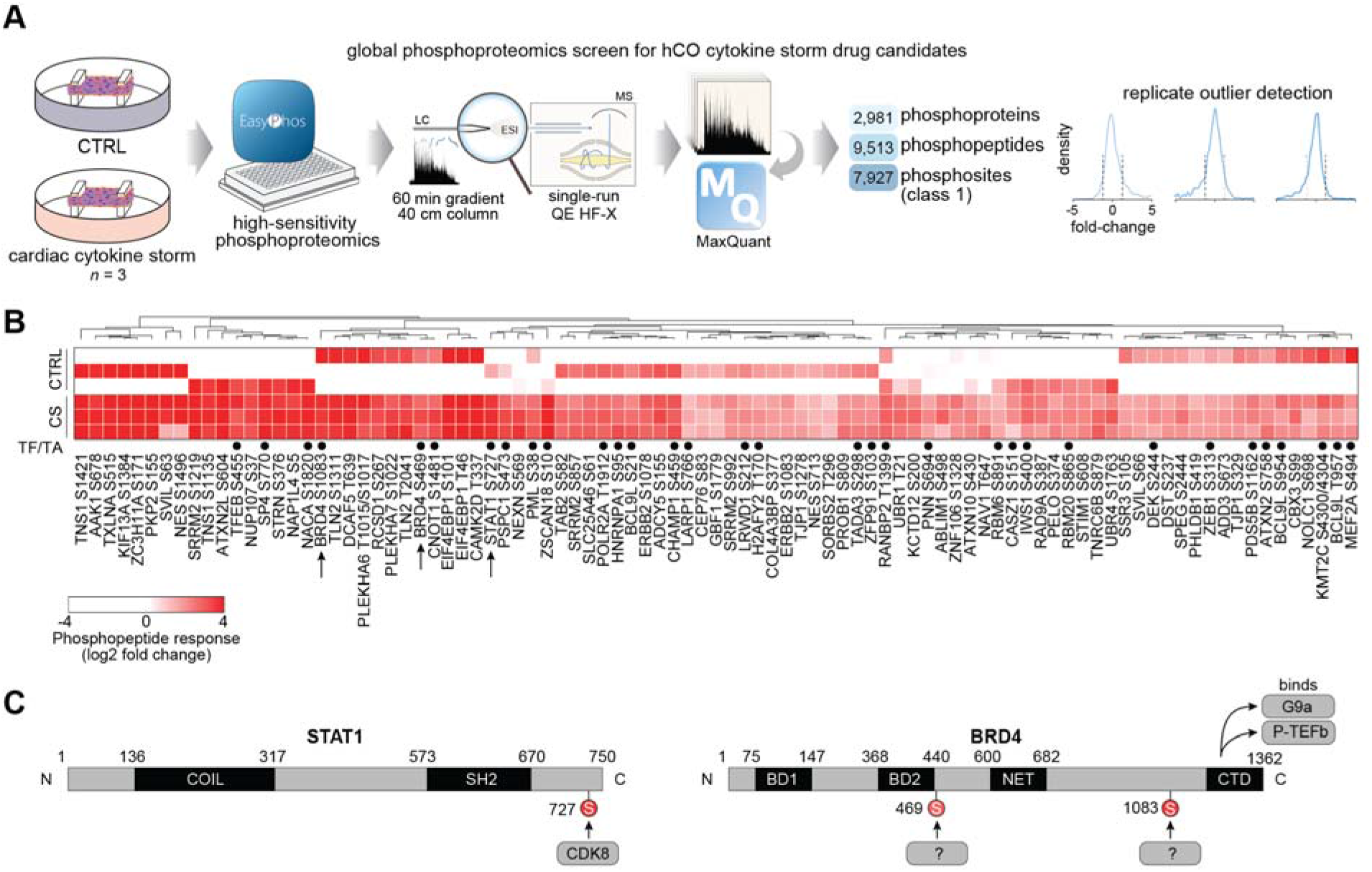
Phosphoproteomics reveals signaling driving cardiac dysfunction. A) Schematic of the experiment and analysis. B) Heat-map of enriched phosphopeptides in hCO following CS treatment (after 1 h). TF/TA circles depict transcription factors and transcriptional activators. C) Phosphorylation sites induced by CS on STAT1 and BRD4. hPSC cardiac cells- AA, Endothelial cells- RM3.5.

**Figure 3:**
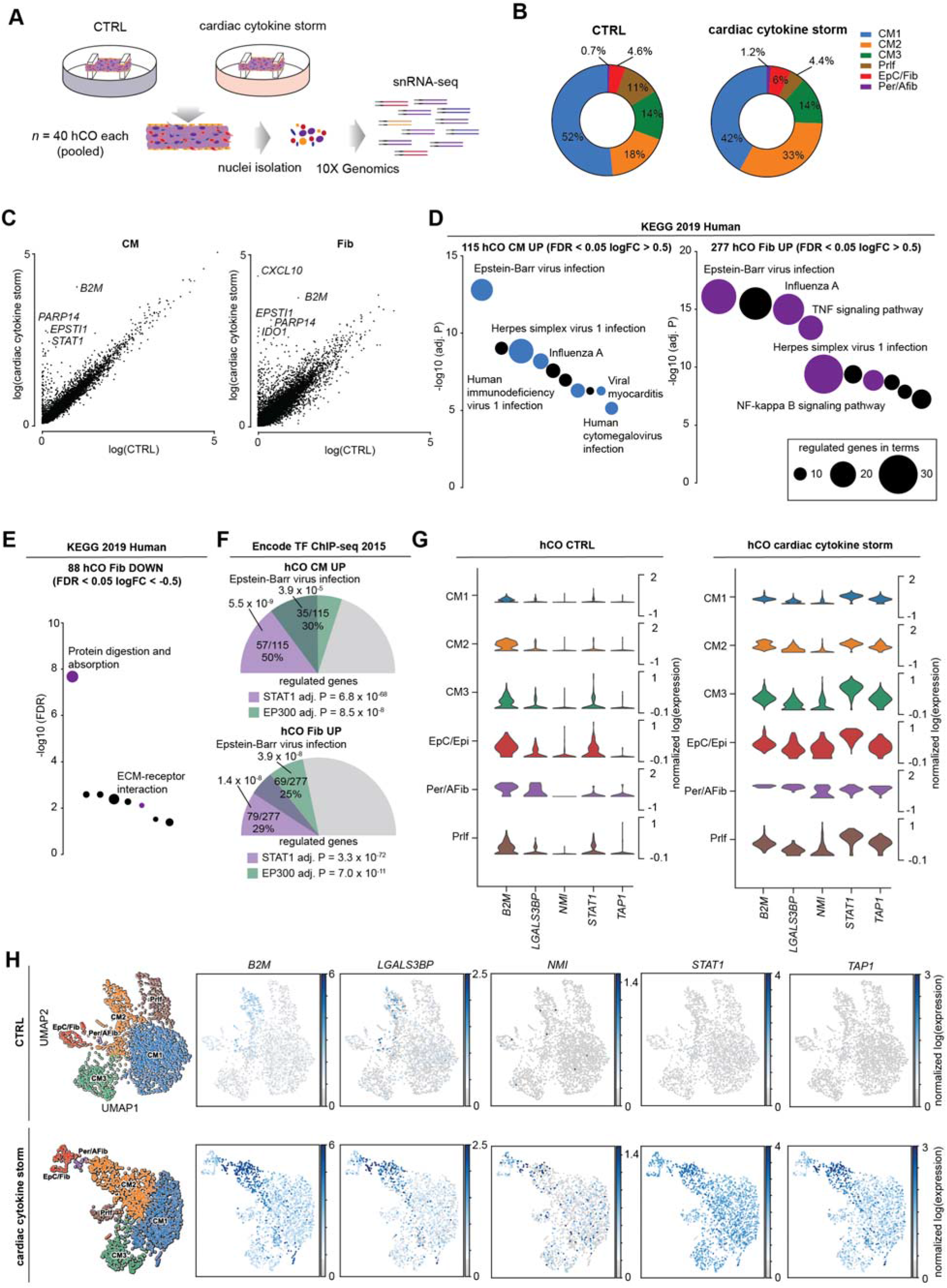
snRNA-seq reveals CS activates viral responses in hCO. A) Schematic of experiment. B) Cell compositions identified in snRNA-seq. C) Differential normalized log2 expression in cardiomyocytes and fibroblasts following CS treatment (all populations pooled for each cell type). D) Activation of viral responses in cardiomyocytes and fibroblasts revealed using KEGG pathway analysis of upregulated genes. Size represents number of genes regulated and the pathways of the coloured circles are highlighted by the text. E) Repression of ECM processes in fibroblasts revealed using KEGG pathway analysis of down-regulated genes. Size represents number of genes regulated and the pathways of the coloured circles are highlighted by the text. F) Analysis of the transcriptional response predicts STAT1 and EP300 as key mediators. Values presented are adjusted P values, number of genes regulated by the transcription factor/number of genes regulated, and % of genes regulated over the total. The size of the coloured slices represent the fraction of genes regulated (180° = 100%), and overlaps for each transcription factor are also depicted. G) Violin plots of key (see **Figure 5Q**) upregulated genes in CS treated hCO. H) UMAP of CTRL and CS treated hCO subpopulations and expression of key regulated genes. hPSC cardiac cells- HES3, Endothelial cells- RM3.5. CM – cardiomyocyte, Prlf – proliferating, EpC – epicardial cells, Fib – fibroblasts, Per – pericytes, Afib – activated fibroblasts.

**Figure 4:**
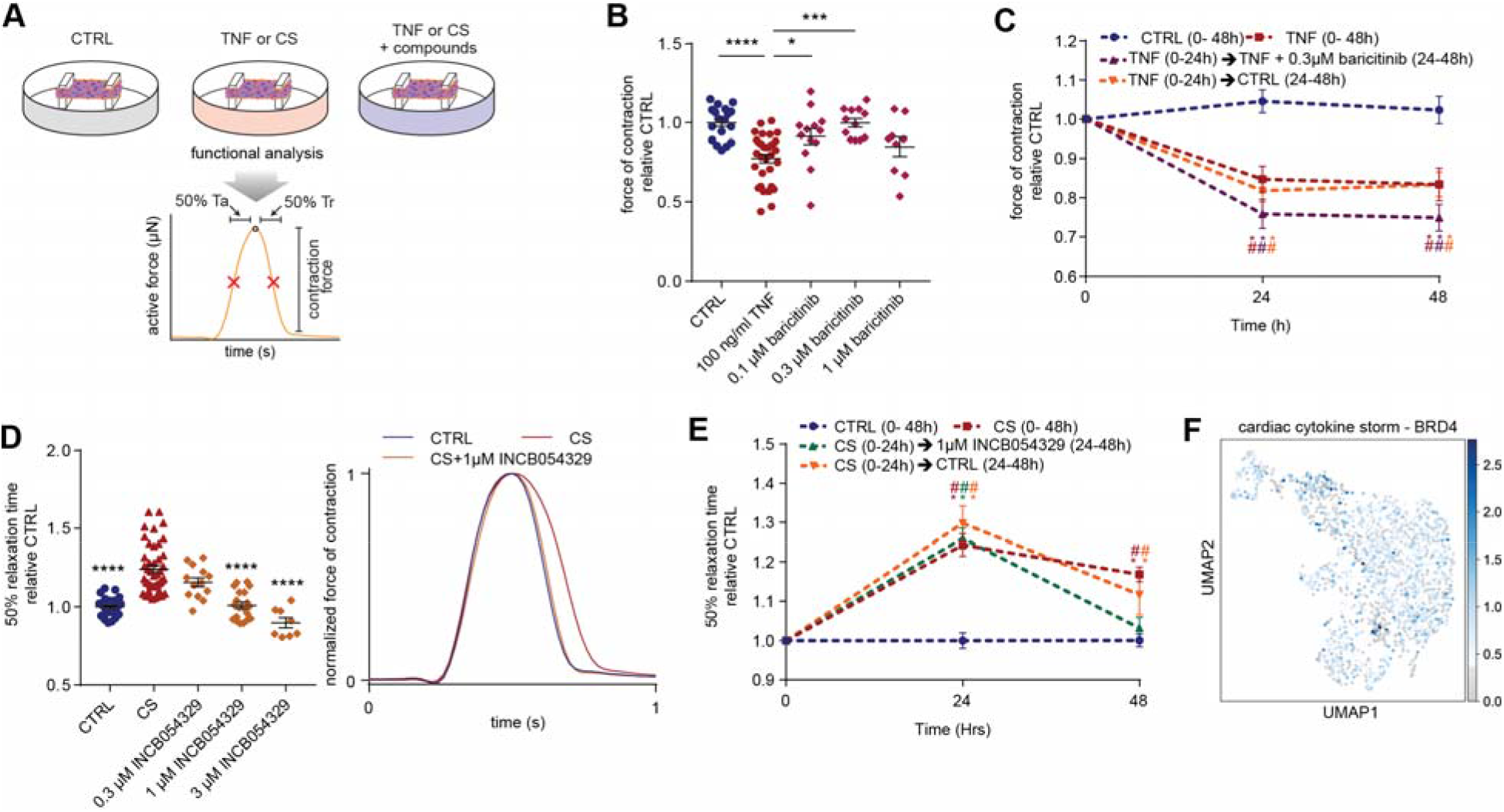
Discovery of drugs that improve cardiac function. A) Schematic of experiment. B) Protection against systolic dysfunction (force of contraction) by baricitinib. n = 9-32 hCOs per condition from 2-3 experiments. *p<0.05, *** p<0.001, **** p<0.0001 using one-way ANOVA with Dunnett’s multiple comparisons test. C) Assessment of hCO recovery from TNF and baricitinib treatment. n = 6-12 hCOs per condition from 1-2 experiments. # p<0.05 compared to CTRL at the same time-point, and * p<0.05 compared to specific condition at 0 h with colour indicating comparison, using two-way ANOVA with Dunnett’s multiple comparisons test. D) Protection against diastolic dysfunction (time to 50% relaxation time) by INCB054329. n = 8-43 hCOs per condition from 2-4 experiments. **** p<0.0001, using one-way ANOVA with Dunnett’s multiple comparisons test compared to CS. E) Assessment of hCO recovery from CS and INCB054329 treatment. n = 6-11 hCOs per condition from 1-2 experiments. # p<0.05 compared to CTRL at the same time-point, and * p<0.05 compared to specific condition at 0 h with colour indicating comparison, using two-way ANOVA with Dunnett’s multiple comparisons test. F) BRD4 is expressed in all cell-populations in hCO. hPSC cardiac cells- HES3, Endothelial cells- RM3.5. Additional drug screen data including a repeat in an additional cell line in **Supplementary Figures S9, S10, S12 and S13**.

We assessed activation of individual cell populations in hCOs using snRNA-seq of ~40 pooled hCOs per condition (**Figure 3A**) with mapping as described above (CTRL - **Figure S2C,D** and CS - **Figure S8A,B**). In the CS condition there was an increase in fibroblast and activated fibroblast number (**Figure 3B**). However, KEGG pathway analysis revealed a transcriptional response dominated by a viral response in both cardiomyocytes (CM1, CM2 and CM3 pooled) and fibroblasts (epicardial/fibroblast and perictye/activated fibroblasts) (**Figure 3C,D**). There were fewer downregulated genes, predominantly in the fibroblasts and these were dominated by extracellular matrix (ECM) genes including *COL1A1*, *COL3A1* and *COL4A5* (**Figure 3E**). The top predicted mediators of the transcriptional response were STAT1 and general epigenetic activation by EP300 (**Figure 3F**). This is consistent with our phosphoproteomic data (**Figure 2**), given EP300 has been shown to share upto 78% of DNA binding regions with BRD4 in chromatin immunprecipitation studies (Williams et al., 2020). These predicted target genes were also enriched for viral responses (**Figure 3F**). These analyses together revealed a robust viral response in the heart in multiple cell populations (**Figure 3G,H**), predicted to be mediated via STAT1 and epigenetic activation and BRD4.

### Drugs for the prevention and treatment of cardiac dysfunction

We next screened drugs that could potentially treat cardiac dysfunction caused by either TNF-induced systolic dysfunction or CS-driven diastolic dysfunction (**Figure 4A**). TNF is known to induce systolic dysfunction via GRK2 mediated repression of β-adrenergic receptor signaling (Vasudevan et al., 2013). The selective serotonin reuptake inhibitor, paroxetine hydrochloride, can inhibit GRK2 (Schumacher et al., 2015), but we found that it was toxic at effective *in vitro* concentrations (Guo et al., 2017) (**Figure S9A**). GRK2 promotes clathrin-mediated endocytosis (Evron et al., 2012), and baricitinib was recently identified as a potential AP2-associated protein kinase 1 (AAK1)-mediated endocytosis inhibitor using machine learning (Richardson et al., 2020). Baricitinib prevented TNF-induced dysfunction in hCO (**Figure 4B** and **Figure S9A,B**). However, baricitinib was only protective against TNF-induced systolic dysfunction when co-administered with TNF and was not effective after 24 h TNF treatment (**Figure 4C**), possibly due to a reduction in cell surface receptor abundance. Additionally, hCO did not recover quickly from TNF-induced systolic dysfunction after the removal of TNF (**Figure 4C**) indicating that secondary remodeling events may have occurred.

A key signature of diastolic dysfunction under CS conditions was the elevated phosphorylation of transcriptional regulators. STAT1-S727 (**Figure 2C**) is associated with assembly into chromatin and is required for STAT1 transcriptional and biological activity in response to IFN-γ (Sadzak et al., 2008). The putative STAT1-S727 kinase is CDK8 (Bancerek et al., 2013), so we next tested two CDK8 inhibitors SEL120-34A (Rzymski et al., 2017) and BI-1347 (Hofmann et al., 2020) previously shown to reduce STAT1-S727 phosphorylation. We also tested two inhibitors of the JAK/STAT pathway, baricitinib and ruxolitinib. However, none of these compounds, nor a broader spectrum CDK inhibitor flavopiridol, prevented the CS-induced diastolic dysfunction (**Figure S10**), noting that flavipiridol was toxic and reduced force and hence all kinetic parameters. Notably, SEL120-34A and BI-1347 specifically attenuated the rate and activation time defects under CS conditions (**Figure S10B,C,E,F**), which we validated in additional experiments (**Figure S11A-D**) and may still have clinical utility in this setting.

We observed elevated phosphorylation of the epigenetic regulator BRD4 and other epigenetic regulators in our CS treated hCO phosphoproteome, consistent with our snRNA-seq analysis. We have previously shown that bromodomain extraterminal inhibitors (BETi) reduce relaxation time in immature hCO (Mills et al., 2019), so we next evaluated three BETi available in an FDA compound library INCB054329 (Stubbs et al., 2019), JQ-1 (Filippakopoulos et al., 2010), and ABBV-744 (Faivre et al., 2020). Strikingly, INCB054329 prevented CS-induced diastolic dysfunction in a dose-dependent manner (**Figure 4D, Figure S12**) without affecting force or rate (**Figure S10A-G, Supplementary Video 5**). JQ-1 also showed improved diastolic function in one hPSC line at the highest concentration (**Figure S10H**), so an additional higher concentration for both JQ-1 and ABBV-744 were tested. JQ-1 protected hCO against CS-induced diastolic dysfunction, although INCB054329 was the most efficacious (**Figure S13A,B**). In contrast, ABBV-744 increased diastolic dysfunction in the hCO, potentially via its dual actions as an androgen receptor inhibitor, which is associated with prolonged QTc in patients undergoing androgen deprivation therapy (Gagliano-Jucá et al., 2018). To validate BRD4 as a target we used adeno associated virus 6 (AAV6) mediated delivery of shRNA and demonstrated that ~74% knockdown could also reduce diastolic dysfunction in CS treated hCO (**Figure S14**).

INCB054329-mediated BETi rescued dysfunctional hCO, and restored diastolic function following 24 h of CS conditions (**Figure 4E**). This is potentially because CS-induced diastolic dysfunction is reversible and is driven by the presence of the inflammatory mediators, demonstrated by partial hCO recovery 24 h after removing CS factors (**Figure 4E**). In patients, all inflammatory factors may be present simultaneously, and we found that INCB054329 attenuated diastolic dysfunction with all four dysfunction inducing factors, TNF, IFN-γ, IL-1β, and poly(I:C), present (**Figure S13C**). Taken together CS mediates diastolic dysfunction via BRD4-dependent mechanisms that can be blocked using BETi. As BRD4 is broadly expressed in our hCO (**Figure 4F**), it may also be responsible for the multi-cellular response observed.

### INCB054329 reduces the host response to SARS-CoV-2 infection in K18-hACE2 mouse hearts

We next assessed the response to SARS-CoV-2 infection *in vivo*. As mice are not susceptible to SARS-CoV-2 infection, we used a recently described K18-hACE2 model (Oladunni et al., 2020) to study the response and effects of BETi (**Figure 5A**). SARS-CoV-2 infected mice had severe lung pathology and substantial viral RNA reads in the lungs at 4-5 d.p.i. confirming successful lung infection (**Figure 5B,C**). RNA-seq of the lungs revealed increased expression of 419 genes (**Figure 5D**), strongly associated with a viral response (**Figure 5E, Table S1**). Concordant with our hCO data, Stat1 and Ep300 were top predicted transcriptional regulators (**Figure 5F, Table S2**). Interestingly, there was only negligible infection of the heart (**Figure 5C**) and no obvious pathology including necrosis, immune cell infiltration or fibrosis (**Figure 5G**). However, there was a substantial and robust upregulation of a viral response in the heart with ECM repression (**Figure 5H,I, Table S1**) including *Col1a1*, *Col3a1* and *Col4a2*. This was again enriched for Stat1 and Ep300 as top predicted transcriptional regulators (**Figure 5J, Table S2**), indicating a robust systemic response in the hearts of SARS-CoV-2 infected mice. This response could be partially blocked by INCB054329 treatment with repression of 91 genes that were enriched for the viral response (**Figure 5K-M, Table S1**). This response was more specfic to the heart, as INCB054329 did not regulate any genes in the lungs (data not shown). The repression by INCB054329 was predicted to be primarily mediated via Ep300 rather than Stat1 (**Figure 5N, Table S2**). These results were further supported by Ingenuity Pathway Analysis of upstream regulators revealing strong activation signatures for IFNG, poly rI:rC-RNA and Stat1 in both lungs and hearts of K18-hACE2 SARS-CoV-2 infected mice, which INCB054329 strongly inhibited (**Table S3**). In addition to the viral response, one of the INCB054329 regulated genes was *Tnni3k* (logFC = −0.34, FDR = 0.0044) which has been shown to be a therapeutic target for cardiac dysfunction (Vagnozzi et al., 2013).

**Figure 5:**
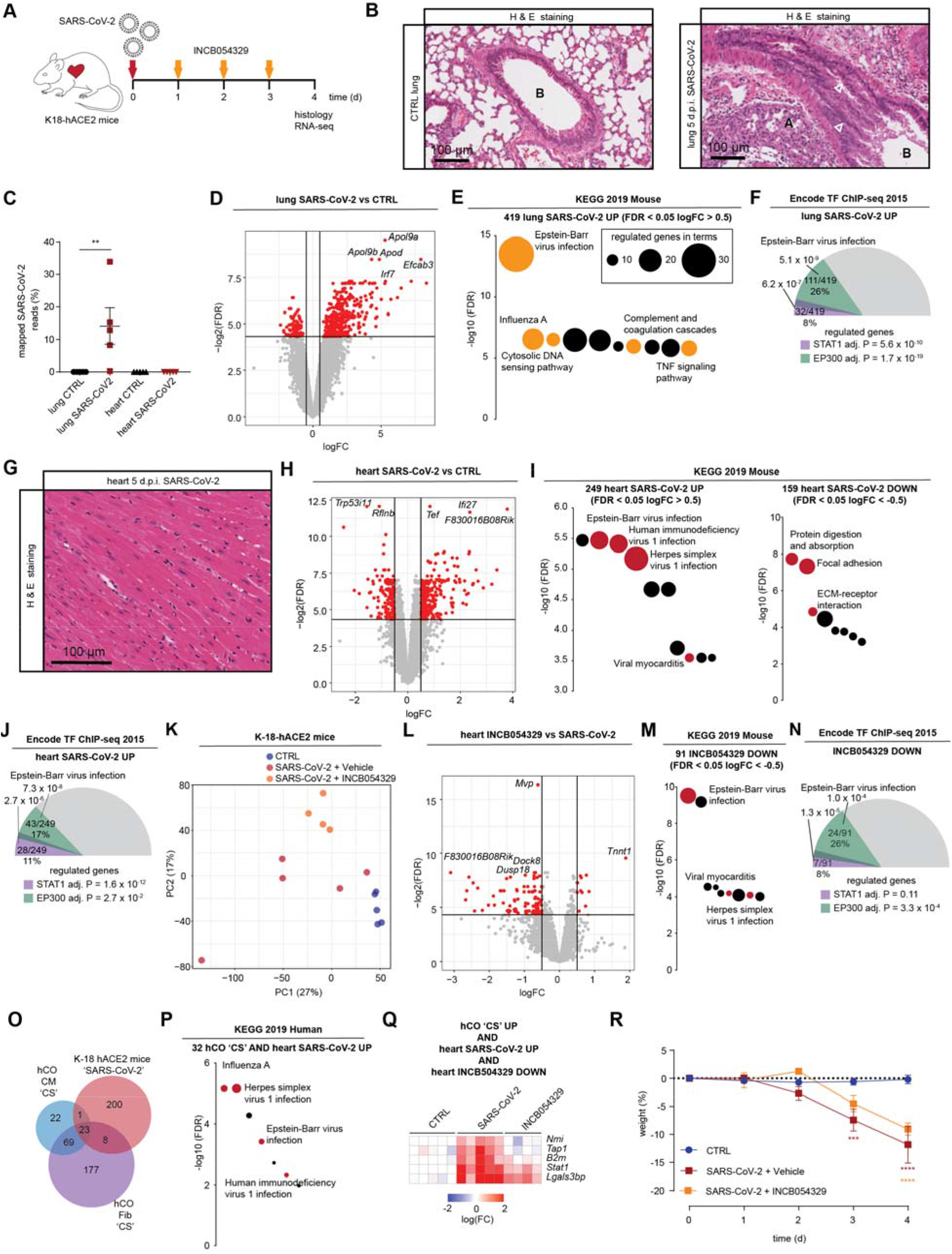
SARS-CoV-2 activates viral responses in the heart despite negligible infection that is repressed by INCB054329. A) Schematic of the experiment. B) Lungs of SARS-CoV-2 infected K18-hACE2 mice 5 d.p.i. In comparison to the control there is sloughing of bronchial epithelium, and white arrowheads show (A) collapse of alveolar spaces and (B) bronchiolar lumen. C) Qualification of mapped reads in RNA-seq reveal severe lung infection with no/negligible heart infection. n = 5 mice per group. D) Volcano plot of lung RNA-seq reveals a robust upregulation (logFC > 0.5) of 419 genes and downregulation (logFC < −0.5) of 98 genes, both FDR < 0.05. n = 5 mice per group. E) Activation of viral responses in lungs revealed using KEGG pathway analysis of upregulated genes. Size represents number of genes regulated and the pathways of the coloured circles are highlighted by the text. F) Analysis of the transcriptional response predicts Stat1 and Ep300 as key mediators of infection in the lungs. G) Hearts of SARS-CoV-2 infected K18-hACE2 mice 5 d.p.i. The hearts appeared relatively normal without significant necrosis, fibrosis (Masson’s Tri-chrome not shown) or immune infiltrates. H) Volcano plot of heart RNA-seq reveals a robust upregulation (logFC > 0.5) of 249 and downregulation (logFC < −0.5) of 159 genes, both FDR < 0.05. n = 5 mice per group. I) Activation of viral responses in hearts revealed using KEGG pathway analysis of upregulated genes. Repression of ECM in hearts revealed using KEGG pathway analysis of downregulated genes. Size represents number of genes regulated and the pathways of the coloured circles are highlighted by the text. J) Analysis of the transcriptional response predicts Stat1 and Ep300 as key mediators of the response in the heart. K) PCA plot of heart RNA-seq samples. n = 4-5. L) Volcano plot of heart RNA-seq reveals a robust upregulation (logFC > 0.5) of 11 genes and downregulation (logFC < −0.5) of 91 genes, both FDR < 0.05 in SARS-CoV-2 infected K18-hACE2 mice treated with INCB054329. n = 4-5 mice per group. M) Repression of viral responses in hearts revealed using KEGG pathway analysis of down-regulated genes. Size represents number of genes regulated and the pathways of the coloured circles are highlighted by the text. N) Analysis of the transcriptional response predicts Ep300 as the key mediator of INCB054329 effects in the heart. O) Cross-analysis of the transcriptional responses in hCO with hearts of SARS-CoV-2 infected K18-hACE2 mice. P) Co-regulated genes in panel (O) reveals a consistent activation of viral responses in both models using KEGG pathway analysis of upregulated genes. Size represents number of genes regulated and the pathways of the coloured circles are highlighted by the text. Q) Heat-map of genes induced by both CS in hCO and SARS-CoV-2 infected K18-hACE2 mouse hearts that are also repressed with INCB054329 treatment. R) Severe weight loss by 4-5 d.p.i in SARS-CoV-2 infected K-18-hACE2 mice is due to severe lung infection and also brain infection and euthanasia is required. d.p.i. – days post infection. Data presented as mean ± standard error of the mean. ** p < 0.01, using Mann-Whitney *** p < 0.001 and **** p < 0.0001 using two-way ANOVA with Sidak’s post hoc test compared to CTRL. For D, H, L – red dots are regulated as per the described cut-offs and grey dots are not. For F,J and N - Values presented are adjusted P values, number of genes regulated by the transcription factor/number of genes regulated, and % of genes regulated over the total. The size of the coloured slices represent the fraction of genes regulated (180° = 100%), and overlaps for each transcription factor are also depicted.

The CS induced response in the hCO (either fibroblasts or cardiomyocytes) and hearts of SARS-CoV-2 infected K18-hACE2 mice shared 32 regulated genes (**Figure 5O**) which were enriched for the viral response (**Figure 5P**). This indicated that CS mimics the paracrine response in the heart caused by SARS-CoV-2 infection and was further supported by Ingenuity Pathway Analysis, which predicted that the inflammatory networks were activated by IFN-γ (**Figure S15**).

Potentially important mediators and markers of the response were found by integrating the multiple datasets. The consistent transcriptional program in CS treated hCO (both fibroblasts and cardiomyocytes) and SARS-CoV2 infected K18-hACE2 mice, that were also down-regulated genes by INCB054329 treatment *in vivo* revaled 5 key targets. These comprise the key inflammatory genes *Nmi*, *Tap1*, *B2m*, *Stat1* and *Lgals3bp* (**Figure 5Q**). Of particular interest is LGALS3BP (galectin-3 binding protein) as it has been shown to be a top-predictor of COVID-19 severity in humans (Messner et al., 2020) and therefore interrogated its expression in our subsequent models.

### INCB054329 protects against inflammatory mediated dysfunction *in vivo* and by COVID-19 serum

The study into the efficacy of SARS-CoV2-related cytokine storm therapeutics on the heart is challenging, not least due to the inaccessibility of mouse echocardiography in biosafety level 3 facilities, but also due to the rapid decrease in weight associated with severe lung/brain infection (**Figure 5R**) that requires euthanasia. Therefore, we tested the ability of INCB054329 to prevent LPS-induced cytokine storm effects *in vivo* (**Figure 6A**). LPS induced pro-inflammatory cytokines TNF, IL-1β and IFN-γ, which were elevated in the plasma (**Figure 6B**) along with *Lgals3bp* in the heart (**Figure 6C**). Treatment with INCB054329 blocked the LPS-induced pro-inflammatory cytokine production (**Figure 6B**) and *Lgals3bp* induction in the heart (**Figure 6C**). We observed a marked improvement in mortality, whereby all INCB054329-treated mice survived after 24 h of the LPS-challenge, compared with only 25% in the control group (**Figure 6D**). To determine whether BETi could treat an established LPS-induced cytokine storm, we delayed injection of INCB054329 1.5 h after LPS injection and assessed cardiac function at 6 h (**Figure 6E**). INCB054329 fully prevented the decrease in cardiac function observed after LPS injection (**Figure 6F**). Together these findings demonstrate that BETi using INCB054329 has robust effects in preventing inflammatory-induced cardiac dysfunction *in vivo*.

**Figure 6:**
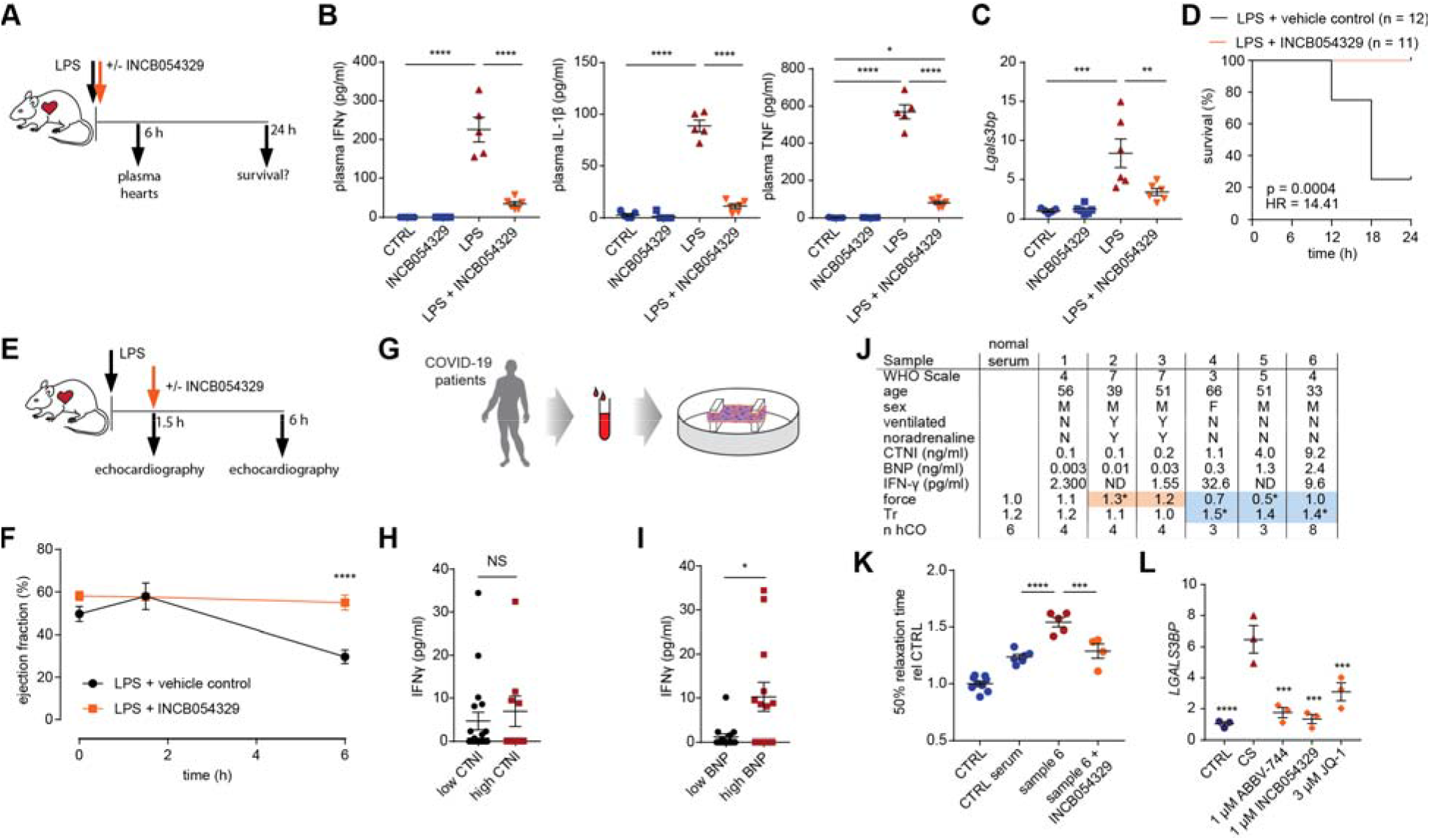
INCB054329 prevents cardiac dysfunction in a mouse LPS-induced cytokine storm model and in response to COVID-19 patient serum. A) Schematic for survival and expression experiments. B) INCB054329 blocks cytokine induction, measured 6 h after LPS using cytokine bead array assays. n = 5-6 mice. *p<0.05, **** p<0.0001, using one-way ANOVA with Tukey’s multiple comparisons test. C) INCB054329 blocks induction of *Lgals3bp* 6 h after LPS using qPCR. n = 5-6 mice. *p<0.05, **p<0.01, *** p<0.001, using one-way ANOVA with Tukey’s multiple comparisons test. D) Kaplan-Meier curve of survival after LPS injection. n = 12 control and 11 INCB054329 treatment (67 mg/kg). P-value calculated using Gehan-Breslow-Wilcoxon test. E) Schematic for the cardiac function experiment. F) INCB054329 prevents the reduction in ejection fraction measured 6 h after LPS using echocardiography. n = 3-4 mice at 0 and 1.5 h and n= 8 at 6 h. **** p<0.0001, using two-way ANOVA with Sidak’s multiple comparisons test. G) Schematic for the human COVID-19 serum experiments. H) IFN-γ was not higher in patients with elevated CTNI (> 0.5 ng/ml). n = 27. I) IFN-γ was higher in patients with elevated BNP (> 0.3 ng/ml). n = 27. J) Serum from COVID-19 patients with elevated BNP induce diastolic dysfunction. * p< 0.05 using one-way ANOVA with Dunnett’s multiple comparisons test compared to CTRL serum. Orange highlights hCO with elevated force of contraction. Blue highlights dysfunctional hCO. K) Diastolic dysfunction induced by COVID-19 patient 6 serum is prevented by 1 μM INCB054329. n = 4-9 hCO from 1 experiment. *** p< 0.001 and **** p < 0.0001 using one-way ANOVA with Dunnett’s multiple comparisons test compared to CTRL serum. L) *LGALS3BP* is induced by CS and repressed by BETi in hCO. n = 3 each 2 x hCO pooled. * p< 0.05, *** p < 0.001, **** p<0.0001 using one-way ANOVA with Dunnett’s multiple comparisons test compared to CS. Data presented as mean ± standard error of the mean. hPSC cardiac cells- HES3, Endothelial cells- RM3.5.

We next assessed whether human serum from COVID-19 patients could induce dysfunction in hCO (**Figure 6G, Supplementary Spreadsheet 1,2**). We found that IFN-γ, one of the key mediators of the cytokine response, was higher in patients with elevated BNP (> 0.3 ng/ml) but not CTNI (> 0.5 ng/ml) (**Figure 6H,I**). In accordance, IFN-γ has been shown to correlate with COVID-19 disease progression and is elevated in patient serum in one of the most comprehensive profiling studies to date (Ren et al., 2021). We were able to demonstrate that hCO function was dictated by factors present in the human serum, as patients receiving noradrenaline as inotropic support elevated contractile force in our hCO (**Figure 6J**). Human COVID-19 patient serum with elevated BNP caused diastolic dysfunction in hCO (**Figure 6J**). INCB054329 prevented diastolic dysfunction caused by patient serum (**Figure 6K**). Induction of LGALS3BP was also prevented by treatment with multiple BETi (**Figure 6L**).

Collectively, these data indicate that BET inhibition with INCB054329 prevented cardiac dysfunction in multiple inflammatory models, as well as repressing the key COVID-19 severity marker LGALS3BP.

### INCB054329 decreases hACE2 expression and reduces SARS-CoV-2 in hPSC-cardiac cells

In our SARS-CoV-2 K-18 mouse infection studies INCB054329 treatment reduced endogenous *Ace2* in hearts *in vivo* (**Figure S17A**). Recently other investigators have also demonstrated that BETi reduced ACE2 *in vivo* and SARS-CoV-2 infection (Qiao et al., 2021). As SARS-CoV-2 potentially infects human hearts and has been shown to infect human pluripotent stem cell-derived cardiac cells (hPSC-CM) (Sharma et al., 2020), we sought to determine whether BETi blocked infection (**Figure S16A**). We confirmed previous findings using 2D cultured hPSC-cardiac cell infection studies, where increasing the MOI increased the degree of cell death (**Figure S16B**). Infection with a low MOI (0.01) was sufficient for viral replication and cell death over 7 days (**Figure S16C**). A 3 day pre-incubation of INCB054329 was sufficient to reduce ACE2 expression ~4 fold (**Figure S16D**). Consequently, pre-treatment with INCB054329 reduced SARS-CoV-2 N protein expression (**Figure S16E**) and intracellular viral RNA (**Figure S16F**). In addition to INCB054329, the widely used BETi JQ-1 reduced SARS-CoV-2 RNA (**Figure S16G**). Thus, BETi also has potential to block SARS-CoV-2 infection of cardiac cells in addition to preventing dysfunction.

### BETi for translation to the clinic

We assessed the ability of all commercially available BETi compounds to prevent CS-induced diastolic dysfunction in hCO. We found that all compounds prevented dysfunction except for ABBV-744 (**Figure 7A**). As BETi with dual BD1 and BD2 activities display side-effects (Gilan et al., 2020), it is critical that we determine selectivity of the response. BD2-selective drugs such as ABBV-744 and apabetalone have limited side effects. Apabetalone has been used for up to 3 years in >1,700 humans at risk of cardiac disease with efficacy in preventing heart failure and favourable safety profile (Nicholls et al., 2021; Ray et al., 2020). ABBV-744 elevated diastolic dysfunction (**Figure 7A**), and we suspect its lack of efficacy was due to its on-target inhibition of the androgen receptor (AR). BD2-specific efficacy was confirmed using a novel BD2-specific molecule, RXV-2157 (**Figure 7B**) and we also confirmed efficacy with the BD-2 selective apabetalone (**Figure 7A,B**). Additionally, analysis of plasma from the ASSURE phase IIb clinical trial indicated that BD2-selective apabetalone reduced LGALS3BP in patients with cardiovascular disease (**Figure 7C**), which is a marker of COVID-19 severity (Messner et al., 2020).

**Figure 7:**
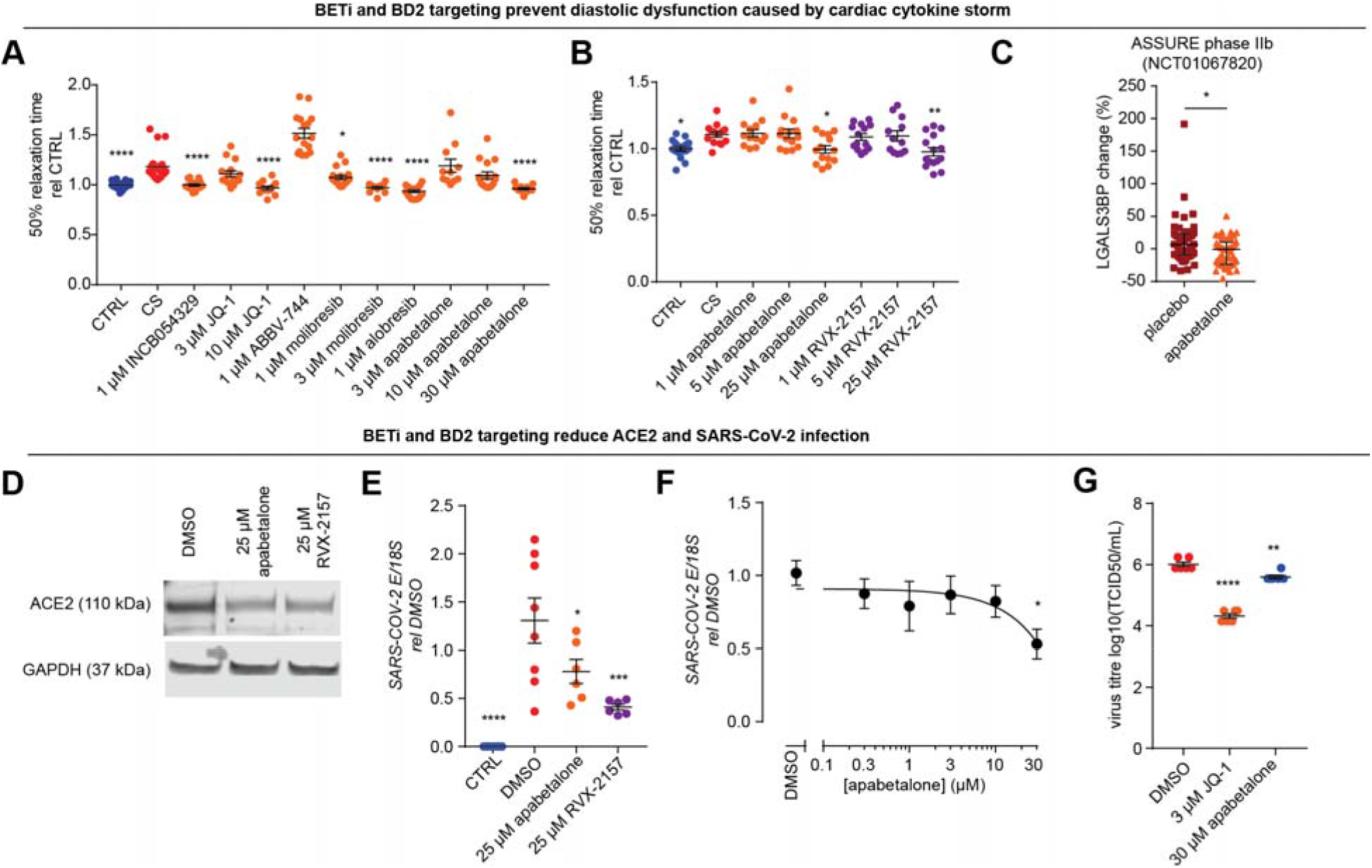
BETi using BD2-selective molecules prevent CS induced diastolic dysfunction in hCO and SARS-CoV-2 infection. A) All BETi (except ABBV-744) previously used in clinical trials prevent CS-induced diastolic dysfunction. n = 12-19 hCOs per condition from 3 experiments. hPSC cardiac cells- HES3 (no endothelial cells). B) BETi specific for the BD2 domain (RVX-2157) or selective for the BD2 domain (apabetalone) prevent CS-induced diastolic dysfunction. n = 12-18 hCOs per condition from 3 experiments. hPSC cardiac cells- HES3 (no endothelial cells). C) Apabetalone decreases serum LGALS3BP in the phase IIb ASSURE clinical trial. Data are presented as a change from baseline. n = 47 placebo and 47 apabetalone. D) BETi specific for the BD2 domain (RVX-2157) or selective for the BD2 domain (apabetalone) decrease ACE2 expression after 3 days. hPSC cardiac cells – HES3. E) Pre-treatment with BETi specific for the BD2 domain (RVX-2157) or selective for the BD2 domain (apabetalone) for 3 days block SARS-CoV-2 infection. E-gene expression in 2D cultured hPSC-CM 3 days after infection. n = 6-8 from 2 experiments (pooled from hPSC cardiac cells- HES3 and AA). F) Dose-response curve for apabetalone 3 day pre-treatment to block SARS-CoV-2 infection. E-gene expression in 2D cultured hPSC-CM 3 days after infection. n = 6 from 1 experiment. hPSC cardiac cells- HES3. G) Apabetalone or JQ-1 3 day pre-treatment reduces SARS-CoV-2 titre in hPSC-CM 3 days after infection. n = 6 from 1 experiment. hPSC cardiac cells- HES3. TCID50 - Fifty-percent tissue culture infective dose. Data presented as mean ± standard error of the mean for all plots except for C which is median ± interquartile range. * p < 0.05, ** p < 0.01, *** p < 0.001, **** p<0.0001, using one-way ANOVA with Dunnett’s multiple comparisons test (A,B – CS, E,F,G - DMSO) or Mann-Whitney (C).

We confirmed the BD2-specificity of blocking SARS-CoV-2 infection. The BD2-specific RXV-2157 and BD2-selective apabetalone molecules downregulated hACE2 (**Figure 7D**), which led to decreased surface expression (**Figure S16B**) and SARS-CoV-2 spike protein binding (**Figure S16C**). These compounds also reduced SARS-CoV-2 RNA (**Figure S17D**, **Figure 7E,F**) and viral titre, including a 2.6 fold decrease in viral titre with apabetalone (**Figure 7G**).

Together, this demonstrates that BD2 -selective BETi drugs are lead candidates for rapid clinical translation to prevent COVID-19 injury in the heart.

## DISCUSSION

Cardiac dysfunction could be caused by systemic inflammation (Del Valle et al., 2020) or local cytokine production (Lindner et al., 2020). In this study we show that inflammatory mediators directly impacted cardiac function in hCO, a model free from the secondary effects and neurohormonal compensation present *in vivo*. We found that a combination of classical viral response cytokines IFN-γ and IL-1β, combined with dsRNA mimics cytokine storm, resulting in severe diastolic dysfunction manifested by 20-50% increases in relaxation time without decline in systolic function. This is consistent with clinical data from heart failure with preserved ejection fraction (HFpEF) patients, where cardiomyocytes have increased time to 50% relaxation by ~13-18%, with similar overall values of 100-150 ms in both humans and hCO (Runte et al., 2017). It is also consistent with diastolic dysfunction reported in COVID-19 patients (Szekely et al., 2020). Therefore, our hCO model recapitulates key clinical features of diastolic dysfunction.

Our data highlight fibroblasts as key mediators in the inflammation-induced cardiac dysfunction. The receptors for inflammatory cytokines are more abundant in cardiac fibroblasts than other cardiac cell types both in hCO and *in vivo* (**Figure S2**), and CS elicited a more pronounced transcriptional response in the fibroblasts within hCO (**Figure 3C,D**). The interplay and intra-organ paracrine signalling between cardiac cell types is likely involved in this inflammation-induced cardiac dysfunction.

By performing high-sensitivity phosphoproteomics, RNA-seq and drug screening with our optimized hCO we identified a therapeutically tractable inflammation-BRD4 axis in inflammation-induced cardiac dysfunction. Our transcriptional profiling revealed a striking similarity between the inflammatory response in CS treated hCO and the *in vivo* response of SARS-CoV-2 infected K18-hACE2 mice, despite negligible cardiac infection in the heart. This is important, as systemic inflammatory signalling may be an important aspect of multi-organ dysfunction documented in COVID-19 patients and other inflammatory diseases.

Our data establish BET inhibition as a viable therapeutic strategy to attenuate cytokine storm-induced cardiac dysfunction. Previously, BETi compounds have shown efficacy in small animal cytokine storm models (Nicodeme et al., 2010). There is also compelling evidence implicating bromdodomains as key mediators in pathological pro-fibrotic signaling in heart failure (Duan et al., 2017; Stratton et al., 2019), in experimental models of pressure overload and myocardial infarction–induced HF (Anand et al., 2013) and in genetic cardiomyopathies (Antolic et al., 2020; Auguste et al., 2020). However, this study is instrumental in establishing BET inhibition as a therapeutic intervention to prevent cardiac dysfunction caused by inflammation.

Clinical data from COVID-19 patients also point to additional cardiac pathologies. Microthrombi were reported in the hearts of 14 out of 40 patients (35%) that died from COVID-19, which was associated with areas of myocardial necrosis (Pellegrini et al., 2021). Consistent with these observations, we observed that “complement and coagulation cascades” were enriched in the lungs of K18-hACE2 mice with SARS-CoV-2 infection (**Figure 5E** and **Table S1**), and were likely related to the viral-induced inflammatory response. Intriguingly, we identified *serpine1* (also known as plasminogen activator inhibitor 1), was upregulated both in the lungs (3.9 fold) and in the heart (2.8 fold, both FDR < 0.05) of SARS-CoV-2 infected K18-hACE2 mice, which was abrogated in the heart upon treatment with the BETi INCB054329 (FDR < 0.05). *Serpine 1* is a key inhibitor of tissue-type plasminogen activator and urokinase-type plasminogen activator, and is required for fibrinolysis down-regulation and degradation of blood clots. In addition, arrhythmic events have been widely reported in COVID-19 patients (Nishiga et al., 2020). Indeed, we observed that arrhythmic events increased in hCO with CS, for which INCB054329 also conferred protection (**Figure S18A-C**). These data suggest that BET inhibition may be effective in attenuating multiple deleterious aspects of systemic inflammation on the heart that warrant further investigation.

We demonstrated that BETi are attractive therapeutic candidates, however the side effect profiles of some BETi may preclude their use in the clinic. Genetic ablation studies have shown that BRD4 plays an integral homeostatic role in cardiomyocytes, suggesting that the loss of BET proteins may have detrimental effects on mitochondrial energy production (Kim et al., 2020; Padmanabhan et al., 2020). Emerging evidence dissecting the roles of BD1 and BD2 bromodomains in inflammatory disease models has indicated that BD2-selective inhibition preferentially blocks the induction of gene expression while minimally affecting established transcription programs (Gilan et al., 2020). More recently, BD2-selective drugs such as ABBV-744 and apabetalone have been developed to overcome these side-effect profiles. Whilst ABBV-744 was not effective in our hCO model (potentially due to its targeting of AR), we demonstrate that BD-2 selective compounds, RXV-2157 and apabetalone demonstrate efficacy. This underscores the need for careful BETi selection, despite broad ability to modulate critical target genes (**Figure 6L**) and utility for a variety of clinical conditions (Cochran et al., 2019). Importantly, BD2-selective BETi apabetalone reduced CS-induced diastolic dysfunction and downregulated ACE2 and reduced viral infection (**Figure S16 and Figure 7**). Taken together, the efficacy and known safety profile of apabetalone make it a prime candidate to protect against cardiac injury for inflammatory diseases such as COVID-19.

The overlap in risk factors for HFpEF and COVID-19 mortality suggests that our findings may also have broader implications. HFpEF risk factors including diabetes and obesity are also associated with chronic inflammation. Recent studies have shown that elevated inflammatory markers are associated with worsening heart function in HFpEF (Sanders-van Wijk et al., 2020), thus indicating that inflammation may drive dysfunction across multiple cardiac diseases and that BET inhibitors are putative therapeutic candidates.

## Supporting information

Mills et al Supplementary Data Biorxiv

## ACKNOWLEDGEMENTS

We thank Clive Berghofer and Lyn Brazil (and others) for their generous philanthropic donations to help set up SARS-CoV-2 research at QIMR Berghofer MRI. We used the Australian National Fabrication Facility Queensland Node for the fabrication of the Heart-Dyno molds. We thank Dr I Anraku for his assistance in managing the PC3 (BSL3) facility at QIMR Berghofer MRI. We thank Dr Alyssa Pye and Mr Fredrick Moore (Queensland Health, Brisbane) for providing the SARS-CoV-2 virus. We thank Grace Chojnowski and Michael Rist for FACS at QIMR Berghofer. Microscopy was aided by Tam Nguyen and Nigel Waterhouse at QIMR Berghofer. Prof Edouard Stanley for provision of the RM3.5 iPSC line (Murdoch Children’s Research Institute, Melbourne, Australia). We thank Nadine Shultz and Paul Collins for the sequencing and also Scott Wood, Pamela Mukhopadhyay, John Pearson, Nic Waddell and Ross Koufariotis for bioinformatics assistance. We thank Compounds Australia (www.compoundsaustralia.com) for providing access to compounds, however all experiments herein used compounds sourced from MedChem Express or Selleckchem. We thank the Translational Research Institute for providing core facilities that enabled this research, particularly Preclinical Imaging and Biological Resources Facility.

We acknowledge grant and fellowship support from the National Health and Medical Research Council of Australia (J.E.H., M.J.S., C.R.E., T.B.), Heart Foundation of Australia (J.E.H.), QIMR Berghofer Medical Research Institute (J.E.H.), The Stafford Fox Foundation (E.R.P.), the Royal Children’s Hospital Foundation (E.R.P.), Australian Research Council Strategic Initiative in Stem Cell Science (Stem Cells Australia) (E.R.P. and J.E.H.) and the Medical Research Future Fund (MRFF9200008) (J.E.H., T.B., M.J.S., K.P.A.MD., C.R.E., E.R.P.). M.J.S. is supported by Health and Medical Research Council of Australia Program (APP1132519) and Investigator (APP1173958) grants. A.S. is also supported by Investigator grant (APP1173880). Queensland Health supported this research project. The Murdoch Children’s Research Institute is supported by the Victorian Government’s Operational Infrastructure Support Program. This project received support from Dynomics Inc. J.E.H. is supported by a Snow Medical Fellowship.

## AUTHOR CONTRIBUTIONS

R.J.M., S.J.H., G.A.Q-R., S.K., M.L., L.T.R., R.R., K.A.S., B.W.C.T., D.J.R., T.L., S.R.F., W.Z., L.L., C.M-K., D.A-B., K.K., T.B., J.E.H., performed experiments, L.D., H.K.V.,L.T.R., E.M., developed cardiac organoid cultures, R.J.M., S.J.H., P.R.J.F., G.A.Q-R., M.L., N.R.M., D.G., L.F., E.K., R.L.J., T.D., C.B., B.G., D.T., R.R., K.S., E.R.P., T.B., J.E.H., analysed data, J.M., C.H., D.G., L.F., S.J.N., J.J., M.S., N.C.W.W., E.K., provided clinical samples, R.J.M., S.J.H., M.J.S., C.R.E., K.P.A.MD., D.G., E.K., T.B., D.A.E, K.S., A.S., D.E.J, J.E.H, designed the project, R.J.M., S.J.H., M.J.S., C.R.E., K.P.A.MD., D.G., L.F., E.K., R.R., K.S., E.R.P., A.S., T.B., D.E.J, J.E.H, interpreted data, R.J.M., S.J.H., D.E.J, J.E.H. wrote the manuscript, all authors edited the manuscript.

## MODELS

### Mice

Mouse work was undertaken in accordance with the Australian Code for Care and Use of Animals for Scientific Purposes, as outlined by the National Health and Medical Research Council of Australia. Animal work was approved by the QIMR Berghofer Medical Research Institute and University of Queensland Animal Ethics Committees.

For LPS experiments wild-type (WT) C57BL/6 were purchased from Walter and Eliza Hall Institute for Medical Research, the Australian Research Centre in Western Australia or bred in house at QIMR Berghofer Medical Research Institute. Mice used in this study were older than 6 weeks and were sex-matched. The number of mice in each group of treatment for each experiment is indicated in the figure legends. No mice were excluded based on pre-established criteria and randomization was applied immediately prior to treatment in therapy experiments.

For SARS-CoV-2 infection studies, heterozygous K18-hACE2 C57BL/6J mice (strain B6.Cg-Tg(K18-ACE2)2Prlmn/J) were purchased from The Jackson Laboratory, USA, and bred in house at QIMR Berghofer Medical Research Institute and genotyped using standard PCR as per Jackson Labs genotyping protocol.

### Cell lines

Ethical approval for the use of human embryonic stem cells (hESCs) was obtained from QIMR Berghofer’s Ethics Committee and was carried out in accordance with the National Health and Medical Research Council of Australia (NHMRC) regulations. hESCs utilized were female HES3 (WiCell), and male RM3.5 iPSC line (generated by Edouard Stanley (Murdoch Children’s Research Institute, Melbourne, Australia)). The following cell lines were obtained from the CIRM hPSC Repository funded by the California Institute of Regenerative Medicine: CW30382A (male, designated AA) and CW30318C (female, designated CC) which were both obtained from FujiFilm. hPSC lines were maintained in mTeSR-1 (Stem Cell Technologies)/Matrigel (Millipore) and passaged using TrypLE (ThermoFisher Scientific) or ReLeSR (Stem Cell Technologies). Quality control was performed with Karyotyping and mycoplasma testing.

### Human COVID-19 plasma and serum

Plasma samples were obtained from individuals with PCR confirmed COVID-19 infection in the community or hospital as a part of the COVID-19 Biobank (Alfred Human Research and Ethics Committee - Project 182/20). Individuals consented to provide additional blood that was processed within 24 h of collection for plasma and peripheral blood mononuclear cells. Whole blood was centrifuged at 1000 x g for 10 min at 22°C-24°C then plasma removed within 5 mm of the buffy coat. Plasma aliquots were transferred to 2 ml cryovials for long term storage at – 80°C. Plasma was thawed and immediately use for ELISA for CTNI, IFN-γ and BNP. Calcium was added to 10 mM to normalize calcium levels of citrated plasma and clot. The supernatant was removed (serum) and used for the hCO experiments (50% serum/50% WM). No viral RNA was detected in hCO treated with human COVID-19 serum.

### Human ASSURE trial plasma

The design and rationale of the ASSURE trial (ClinicalTrials.gov identifier NCT01067820) is described in. Patients with established cardiovascular disease received 200 mg apabetalone daily for 26 weeks on top of standard of care, which included statins. Baseline and end of study EDTA plasma samples were analysed using SOMAScan™.

### SARS-CoV-2 stock production and titration at QIMR Berghofer

SARS-CoV-2 infection studies at QIMR Berghofer were conducted in a dedicated PC3 (BSL3) suite, with safety approval from the QIMR Safety Committee (P3600). The SARS-CoV-2 virus was isolated from a patient and was a kind gift from Queensland Health Forensic & Scientific Services, Queensland Department of Health; the isolate, hCoV-19/Australia/QLD02/2020; has been sequenced as is available at GISAID (https://www.gisaid.org/). Virus stock was generated by infecting Vero E6 cells (C1008, ECACC, Wiltshire, England; Sigma Aldridge, St. Louis, MO, USA) and after 3 days culture supernatant was clarified by centrifugation at 3000 x g for 15 min at 4°C, and was aliquoted and stored at ^-^80°C. Virus titre was determined using standard TCID_50_ assay by infecting Vero E6 cells with 10-fold serial dilutions of virus stock and measuring cytopathic effect with titre calculation by the method of Spearman and Karber. Virus was determined to be mycoplasma free (La Linn et al., 1995) and fetal calf serum used for culture determined to be endotoxin free (Johnson et al., 2005).

### SARS-CoV-2 stock production at The Doherty Institute

SARS-CoV-2 isolate CoV/Australia/VIC01/2020, provided by the Victorian Infectious Diseases Reference Laboratory (VIDRL) was amplified in Vero cells and stock vials were stored at −80°C. The amplified virus was sequenced to confirm that there were no mutations resulting from passage in Vero cells. All work with infectious virus was performed inside a biosafety cabinet, in a biosafety containment level 3 facility, and personnel wore powered air-purifying respirators (3M TR-315A VERSAFLO Cat# RPPKTR315A, from Safetyquip) or P2 masks. Vero cells were obtained from VIDRL and maintained in Minimum Essential Medium (MEM, Media Preparation Unit, Peter Doherty Institute) with 5% FBS, Penicillin-Streptomycin, GlutaMax (and 7.5ml HEPES (all ThermoFisher Scientific).

## METHODS

### Cardiac differentiation

Cardiac differentiation was performed as previously described (Hudson et al., 2012; Mills et al., 2017; Voges et al., 2017). hPSCs were seeded on Matrigel-coated flasks at 2 × 10^4^ cells/cm^2^ and cultured in mTeSR-1 for 4 days. To induce cardiac mesoderm, hPSCs were cultured in RPMI B27-medium (RPMI 1640 GlutaMAX+ 2% B27 supplement without insulin, 200 μM L-ascorbic acid 2-phosphate sesquimagnesium salt hydrate (Sigma) and 1% Penicillin/Streptomycin (ThermoFisher Scientific), supplemented with 5 ng/ml BMP-4 (RnD Systems), 9 ng/ml Activin A (RnD Systems), 5 ng/ml FGF-2 (RnD Systems) and 1 μM CHIR99021 (Stem Cell Technologies). Mesoderm induction required daily medium exchanges for 3 days. This was followed by cardiac specification using RPMI B27-containing 5 μM IWP-4 (Stem Cell Technologies) for another 3 days, and then further 7 days using 5 μM IWP-4 RPMI B27+ (RPMI1640 Glutamax + 2% B27 supplement with insulin, 200 μM L-ascorbic acid 2-phosphate sesquimagnesium salt hydrate and 1% Penicillin/Streptomycin) with media change every 2-3 days. For the final 2 days of differentiation, hPSCs were cultured in RPMI B27+. Harvest of differentiated cardiac cells involved enzymatic digestion, firstly in 0.2% collagenase type I (Sigma) containing 20% fetal bovine serum (FBS) in PBS (with Ca^2+^ and Mg^2+^) at 37°C for 1 h, and secondly in 0.25% trypsin-EDTA at 37°C for 10 minutes. Cells were filtered through a 100 μm mesh cell strainer (BD Biosciences), centrifuged at 300 x g for 3 min, and resuspended in α-MEM Glutamax, 10% FBS, 200 μM L-ascorbic acid 2-phosphate sesquimagnesium salt hydrate and 1% Penicillin/Streptomycin. Previous flow cytometry analysis indicated that differentiated cardiac cells were ~70% α-actinin^+^/CTNT^+^ cardiomyocytes, ~30% CD90 stromal cells (Voges et al., 2017).

### Endothelial differentiation

Endothelial cell differentiation was performed following a protocol modified from (Orlova et al., 2014). hPSCs were seeded onto Matrigel-coated T-25 or T-75 tissue culture flasks at the density 5 × 10^3^ cells/cm^2^ and cultured in mTeSR-1 for 3 days. Mesoderm was induced with RPMI B27- (RPMI 1640 GlutaMAX+ 2% B27 supplement without insulin, 200 μM L-ascorbic acid 2-phosphate sesquimagnesium salt hydrate (Sigma) and 1% Penicillin/Streptomycin (ThermoFisher Scientific), and the small molecules 25 ng/ml Activin A (R&D systems), 30 ng/ml Bone morphogenic protein-4 (BMP4) (R&D systems), 1.5 μM CHIR99021 (Stemgent), and 50 ng/ml Vascular Endothelial Growth Factor type A (VEGF-A) (Peprotech) for 3 days (no media changes). Endothelial cell fate was further specified with RPMI B27- medium supplemented with 50 ng/ml VEGF-A and 10 μM SB431542 with media changes every 2 to 3 days until day 8.

### FACS sorting endothelial cells

Endothelial cells were harvested on day 8 of differentiation using TrypLE (ThermoFisher Scientific). Single cells were separated using a 100 μm strainer and labelled with CD31 antibody (1:200, M082329-2, DAKO) at 4°C for 45 min followed by 30 min staining with a goat anti-mouse secondary antibody conjugated to AlexaFluor 488 or 555 (1:400, A-11001 and A-21422, ThermoFisher Scientific). Cells were analysed using Becton Dickinson FACSAria II, gated on forward and side scatter. Single cells were identified and sorted based on CD31+ expression. CD31+ endothelial cells were expanded in EGM-2 (Lonza) in Matrigel flasks and cryopreserved.

### hCO fabrication

hCO culture inserts were fabricated using SU-8 photolithography and PDMS molding (Mills et al., 2017). Differentiated cells were mixed at ratio of 20% endothelial cells and 80% cardiomyocytes/fibroblasts to form hCO. Acid-solubilized bovine collagen 1 (Devro) was salt balanced using 10x DMEM (ThermoFisher Scientific) and pH neutralized using 0.1M NaOH before combining with Matrigel and then the cell suspension on ice. Each hCO contained 5 × 10^4^ cells, a final concentration of 2.6 mg/ml collagen I and 9% Matrigel. 3.5 μL of suspension was pipetted into the hCO culture insert and incubated at 37°C with 5% CO_2_ for 45 min in order to gel. After gelling, α-MEM GlutaMAX (ThermoFisher Scientific), 10% fetal bovine serum (FBS), 200 μM L-ascorbic acid 2-phosphate sesquimagnesium salt hydrate (Sigma) and 1% Penicillin/Streptomycin (ThermoFisher Scientific) was added. hCO were subsequently cultured in maturation media (MM) (Mills et al., 2017) with medium changes every 2 to 3 days for 5 days (7 days old hCO). To better approximate adult metabolic provisions a ‘weaning medium’ (WM) was utilized. hCO were cultured in WM containing 4% B27 – insulin, 5.5 mM glucose, 1 nM insulin, 200 μM L-ascorbic acid 2-phosphate sesquimagnesium salt hydrate, 1% P/S, 1% GlutaMAX (100x), 33 μg/mL aprotinin and 10 μM palmitate (conjugated to bovine serum albumin in B27) in DMEM without glucose, glutamine and phenol red (ThermoFisher Scientific) with media changes every 2-3 days.

### Force analysis of hCO

The elasticity of the Heart Dyno poles enables the contractile properties to be determined via tracking pole deflection, which directly correlates with force (Mills et al., 2017). Videos of 10 seconds were made of each hCO with the Nikon ANDOR WD Revolution Spinning Disk microscope (magnification 4x). While imaging, hCO were incubated at 37°C, 5% CO_2_ to prevent changes in contractile behaviour. For pacing, hCOs were electrically stimulated at 1 Hz with 5 ms square pulses with 20 mA current using a Panlab/Harvard Apparatus Digital Stimulator. Videos were then analysed with a custom written Matlab program (Mills et al., 2017). This facilitated the analysis of the contractile properties of the organoids and the production of time-force graphs (Mills et al., 2017). Moreover, data was obtained regarding additional important functional parameters including the contraction rate and the activation and relaxation time of the organoids.

### Immunostaining of hCO (endothelial)

hCO were fixed with 1% paraformaldehyde (Sigma) for 1 h. Cells were stained with primary antibodies CD31 (1:200, M082329-2, DAKO), NG2 (1:200, 14-6504-82, ThermoFisher Scientific) and cardiac troponin T (1:400, ab45932, Abcam) in 5% FBS and 0.25% TritonX-100 Blocking Buffer at 4°C overnight on a rocker. Cells were washed twice for 1 h with Blocking Buffer and labelled with secondary antibodies goat anti-mouse IgG_1_ AlexaFluor 488 (1:400, A-21121), goat anti-mouse IgG_2a_ AlexaFluor 555 (1:400, A-21137) and goat anti-rabbit IgG AlexaFluor 633 (1:400, A-21070) and Hoechst33324 (all ThermoFisher Scientific) at 4°C overnight on a rocker. Cells were again washed with Blocking Buffer twice for 1 h and mounted on microscope slides using ProLong Glass (ThermoFisher Scientific).

### Phosphoproteomics

Phosphoproteomics experiments were performed with biological triplicates. Phosphopeptides were enriched from 20 pooled hCO, yielding approximately 100 μg of total protein per condition. The high-sensitivity EasyPhos workflow was employed as previously described (Humphrey et al., 2018). Briefly, pooled organoids were lysed in SDC buffer (4% Sodium deoxycholate, 100 mM Tris pH 8.5) and immediately heated for 5 min at 95°C. Lysates were cooled on ice, and sonicated with a tip-probe sonicator (50% output power, 30 seconds). An aliquot of lysate was taken and diluted 1:5 in 8 M Urea from which protein concentration was determined by BCA assay (Thermo Fisher Scientific). Aliquots corresponding to 100 μg of protein were subsequently diluted in SDC buffer into a 96-well deep-well plate, reduced and alkylated at 45°C for 5 min by the addition of 10 mM Tris (2-carboxyethyl)phosphine (TCEP)/40 mM 2-Chloroacetamide (CAA) pH 8, and digested by the addition of 1:100 Lys-C and Trypsin overnight at 37°C with agitation (1,500 rpm). After digestion phosphopeptides were enriched in parallel according to the EasyPhos workflow as described (Humphrey et al., 2018). Eluted phosphopeptides were dried in a SpeedVac concentrator (Eppendorf) and resuspended in MS loading buffer (0.3% TFA/2% acetonitrile) prior to LC-MS/MS measurement.

### LC-MS/MS Measurement

Phosphopeptides were loaded onto a 40 cm column fabricated in-house with 75 μM inner diameter fused silica packed with 1.9 μM C18 ReproSil particles (Dr. Maisch GmBH). A column oven (Sonation) was used to maintain column temperature at 60°C, and a U3000 HPLC system (Dionex, Thermo Fisher Scientific) was connected to a Q Exactive HF X benchtop Orbitrap mass spectrometer (Thermo Fisher Scientific) with a NanoSpray Flex ion source (Thermo Fisher Scientific). For all samples, peptides were separated using a binary buffer system of 0.1% (v/v) formic acid (buffer A) and 80% (v/v) acetonitrile/0.1% (v/v) formic acid (buffer B). Peptides were eluted at a flow rate of 400 nl/min and separated with a gradient of 3 – 19% buffer B over 40 minutes, followed by 19 – 41% buffer B over 20 minutes, and peptides were analysed with a full scan (350 – 1,400 m/z; R=60,000 at 200 m/z) at a target of 3e6 ions, followed by up to ten data-dependent MS2 scans using HCD (target 1e5; max. IT 50 ms; isolation window 1.6 m/z; NCE 27%; min. AGC target 2e4), detected in the Orbitrap mass analyser (R=15,000 at 200 m/z). Dynamic exclusion (30 s) and Apex trigger (2 to 4 s) were enabled.

### MS data processing

RAW MS data was processed in the MaxQuant software environment (Cox and Mann, 2008) (version 1.6.0.9), searching against the Human UniProt Reference database (December 2019 release), using default settings with the addition of ‘Phospho(STY)’ as a variable modification and ‘Match between runs’ switched on for all analyses. Data analysis was performed using the Perseus software package (Tyanova and Cox, 2018).

### Single nuclei RNA-sequencing of hCO

Pooled hCO (~40) were homogenized in 4 mL lysis buffer (300 mM sucrose, 10 mM Tris-HCl (pH = 8), 5 mM CaCl_2_, 5 mM magnesium acetate, 2 mM EDTA, 0.5 mM EGTA, 1 mM DTT)(all Sigma-Aldrich) with 30 strokes of a dounce tissue grinder (Wheaton). Large pieces of hCO were allowed to settle and homogenate was passed through pre-wetted 40 μm cell strainers (Becton Dickinson). Remaining hCO material in the douncer was resuspended in 4 ml and the douncing and filtering steps were repeated twice. All steps of the homogenization were performed on ice. The filtered homogenate was centrifuged at 1500 x g for 5 min at 4°C. Nuclei pellets were then re-suspended in PBS. A fraction of resuspended nuclei were then stained with Hoechst33324 nuclear stain (1:500 dilution) and counted on a haemocytometer under fluorescent microscope.

The nuclei were then re-centrifuged (1500 x g for 5 min at 4°C) and resuspended at a density to load ~5,000 nuclei per sample. Cells were loaded into the Chromium Controller (10X Genomics) for gel bead emulsion (GEM) formation. Library preparation was conducted according to the manufacturer’s recommended protocol using the Chromium Next GEM Single Cell 3’ GEM, Library & Gel Bead Kit v3.1. Libraries were sequenced on the NextSeq 500/550 v2 (Illumina) with 150 bp reads and were sequenced to ~250,000 reads per cell.

Raw fastq reads for each sample were processed using CellRanger v3.1.0. Default options were used with CellRanger and a custom made pre mRNA reference using GRCh38 v3.0.0 was used to map the reads and for count quantification with the CellRanger counts tool. Following this, the counts were then aggregated together to create a single matrix that contained all the samples. Reads from the single nuclei sequencing data were aligned to a human pre-mRNA GRCh38 reference genome. All pre-processing and filtering steps of the datasets were subsequently carried out via the Python package Scanpy (https://scanpy.readthedocs.io/en/stable/) (Wolf et al., 2018). Briefly, there was an initial filtering step for genes that are expressed in 3 or more cells and cells with at least 200 detected genes, subsequently removing cells displaying high expression of mitochondrial genes using a cut-off of 4%. We then filtered out cells that had a count depth with a threshold of under 2,000 and higher than 40,000 to remove debris and potential doublets. Gene expression was subsequently normalized for each cell by total expression, scaled by 10,000 and log scaled. Highly variable genes were then identified for clustering. Leiden clustering with an initial resolution of 0.2 was performed to identify clusters within the data. A published human cardiac snRNA-seq dataset was then used to identify overlap of marker gene sets for main human heart cell types (e.g. cardiomyocytes, fibroblasts, epicardium, pericytes, etc) with the clusters of our dataset, and were labelled accordingly (Tucker et al., 2020). Further refined clustering was carried out on specific clusters that showed overlap of more than one of the various heart cell types. Visualization of the datasets was primarily carried out using nonlinear dimensionality reduction UMAP plots (Becht et al., 2019). In the snRNA-seq we note the lower than expected percentage of non-myocytes and the loss of endothelial cells (Mills et al., 2017; Voges et al., 2017), indicating the protocol requires further optimization for hCO samples.

### hCO comparison to bulk nuclei RNA sequencing data for PCA

Nuclear RNA-seq dataset generated from sorted cardiomyocyte nuclei at two stages (fetal and adult) was obtained from BioProject ID: PRJNA353755 (Gilsbach et al., 2018). RNA-seq dataset was mapped to the human genome (hg38) using STAR aligner (version 2.7.3a). Annotations and genome files (hg38) were obtained from *Ensembl (release 102).* Uniquely mapped reads were counted across genes with a program in Bioconductor R (Huber et al., 2015) package, featureCounts (version 2.0.1) (Liao et al., 2014). Subsequent analyses of the count data were performed in the R statistical programming language with the Bioconductor packages edgeR (Robinson et al., 2010) and the annotation package org. Hs.eg.db. In this dataset, only genes with > 0.5 counts per million (CPM) in at least 4 samples were retained for statistical analysis. Additionally, ribosomal and mitochondrial genes as well as pseudogenes, and genes with no annotation (Entrez Gene identification) were removed before normalization and statistical analysis.

Principal component analysis (PCA) performed using the intersection of the 25% most highly variable genes of the snRNA-seq dataset and genes expressed in the bulk samples. Each of the two datasets were log transformed and scaled separately before running the dimensionality reduction method.

### Pro-inflammatory stimulation of hCO

Cytokines (human) and factors were added individually and in combinations in WM: 100 ng/ml TNF, 10 ng/ml IL-1β, 100 ng/ml IFN-γ, 100 ng/ml IL-6, 100 ng/ml IL-17A, 100 ng/ml G-CSF (Amgen), 10 μg/ml poly(I:C) (HMW, Invivogen) or 1 μg/ml LPS (from *Escherichia coli* strain 0127:B8, Sigma) (all Peprotech unless noted). Additional concentrations were performed for the dose-response curves for TNF, IL-1β, IFN-γ and poly(I:C) as indicated. The function of hCO was determined before addition of these factors as a baseline (time 0 h) and any changes normalized to the original baseline and to the control. The medium was not exchanged unless noted.

### Drug screening

Compounds were sources from MedChem Express (unless noted) and dissolved at 10 mM in DMSO and vehicle controls used. A larger batch of INCB054329 was sourced from Selleckchem. The following compounds were used at 2 or 3 doses previously shown to have *in vitro* efficacy as per the reference papers for: JQ-1 (Selleckchem), INCB054329, ABBV-744, ruxolitinib, baricitinib, flavopiridol, SEL120-34A, BI-1347, and paroxetine hydrochloride. Additional compounds tested were molibresib, alobresib, and apabetalone. For some experiments apabetalone and RVX-2157 were sent blinded by Resverogix. Compounds were given at the time of pro-inflammatory factor addition, except for experiments with baricitnib and INCB054329 where recovery of function was also assessed by addition 24 h following addition of inflammatory factors.

### Linear regression of cytokine storm responses

To determine factor effects, second order OLS linear regression was performed across all relevant samples using binary predictors (cytokine presence/absence) with force, relaxation/activation times as the outcome variables. p-values were determined using two-tailed t-tests. Normality (Shipiro-Wilks), heteroskedasticity (Breusch-Pagan), linearity (Harvey-Collier), multicollinearity (condition no.) and skewness/kurtosis (Jarque-Bera) were all checked with the respective tests. The coefficient of determination, R^2 (adjusted), was used to determine goodness of fit.

### SARS-CoV-2 K18-hACE2 mouse infection model

Female K18-hACE2 mice were lightly anesthetized using isoflurane and 50 μl of SARS-CoV-2 at 5 × 10^4^ TCID50 per mouse was administered via intranasal inoculation (i.n.). On 1, 2 and 3 dpi, INCB054329 or placebo control was administered via oral gavage. Mice were randomized and received either vehicle 30% (m/v) Kolliphor 15 HS (Sigma) in PBS or 2 mg per 30 g mouse body weight of INCB054329 at 20 mg/ml in the Kolliphor solution (66.7 mg/kg). At 4 or 5 d.p.i., mice were euthanized by cervical dislocation and heart and lung tissue was fixed in 10% formalin for histology. At 4 d.p.i. lungs or hearts were homogenized in TRIzol for RNA extraction and stored at −80°C.

### Bulk RNA-seq from SARS-CoV-2 K18-hACE2 mouse infection model

Illumina Stranded Total RNA Prep with Ribo-Zero Plus kits were used to prepare total RNA for sequencing. Libraries were sequenced on the NextSeq 500/550 v2 (Illumina) with 150 bp reads and were sequenced to ~60,000,000 reads per sample. Sequence reads were trimmed for adapter sequences using Cutadapt version1.9 (Martin, 2011) and aligned using STAR version 2.5.2a (Dobin et al., 2013) to the Mus Musculus GRCm38 assembly with the gene, transcript, and exon features of Ensembl (release 102) gene model, and the SARS-CoV-2 Ensembl genome assembly ASM985889v3. Quality control metrics were computed using RNA-SeQC version 1.1.8 (DeLuca et al., 2012) and expression was estimated using RSEM version 1.2.30 (Li and Dewey, 2011). Protein-coding genes with > 5 CPM in ≥ 5 samples were kept for further analysis. Trimmed mean of M-values (TMM) normalization and differential expression analysis were performed using the R package edgeR (Robinson et al., 2010). The glmQLFit() function was used to fit a quasi-likelihood negative binomial generalised log-linear model to the read counts for each gene. Using the glmQLFTest() function, we conducted gene-wise empirical Bayes quasi-likelihood F-tests for a given contrast. Differentially expressed genes (DEGs) were determined using absolute log2 fold change (logFC) > 0.5 and a false discovery rate (FDR) < 0.05. The function prcomp() was used for principal component analysis (PCA). To estimate SARS-CoV-2 replication levels, sequence reads were aligned to SARS-CoV-2 only, and samtools (Li et al., 2009) version 1.9 was used to estimate the mapping rate of the reads to the viral genes.

### Comparison of different RNA-seq data

KEGG and Encode TF analyses was performed using Enrichr (Kuleshov et al., 2016). BioMart Ensembl (release 102) was used to obtain the mouse to human orthologues used in comparisons. Networks were generated through the use of Ingenuity Pathway Analysis (IPA) on DEGs (QIAGEN Inc., https://www.qiagenbioinformatics.com/products/ingenuity-pathway-analysis). Up-Stream Regulators enriched in differentially expressed genes (no logFC cutoff, FDR < 0.05) in direct and indirect interactions were investigated by performing Core Analysis using (QIAGEN).

### Quantitative RT-PCR

RNA was extracted using QIAGEN RNAeasy Micro Kits (Qiagen) or Trizol. cDNA synthesis using Superscript III (ThermoFisher Scientific) was carried out as per manufacturer’s instructions. Final primer concentration of 200-250 nM was used and gene expression was assessed over 40 cycles on an Applied Biosciences Quant Studio 5. *18S* (for viral infection studies) or *HPRT1* or *hprt* (for gene expression) were used as internal controls.

### Mouse LPS cytokine storm model

LPS (from *Escherichia coli* strain 0127:B8, Sigma) suspended in PBS was injected intraperitoneally into mice at 0.6 mg per 30 g mouse body weight for inflammatory cytokine, gene expression and survival studies and 1 mg was used for cardiac function studies. After the injection of LPS mice were randomized and received either vehicle 30% (m/v) Kolliphor 15 HS (Sigma) in PBS or 2 mg per 30 g mouse body weight of INCB054329 at 20 mg/ml in the Kolliphor solution (66.7 mg/kg). Treatment groups were blinded. For the experiments, mice were closely monitored and checked hourly for signs of sepsis.

### Mouse LPS plasma cytokine assays

Serum cytokine levels (Ifn-γ, Il-1β and Tnf) were determined with a CBA Flex Set Multiplex Cytokine Bead Array (BD Biosciences).

### Cardiac function in vivo

Cardiac function was assessed using a Vevo 2100 ultrasound system fitted with a MS550D transducer, which has a 40 MHz center frequency (Fujifilm Visualsonics). Depilated mice were anaesthetized by isoflurane inhalation (1.5% at 1 L oxygen / min) delivered via a nose cone, kept warm on a heated stage, with respiration and heart rate monitored on ECG pads. B Mode images of the left ventricle were obtained from three short-axis views (proximal, mid and distal positions) and one parasternal long-axis view. Cardiac function parameters were calculated using the Simpson’s tool in Vevolab analysis software v3.2.6 (Fujifilm Visualsonics). Briefly, the endocardial areas were traced from all three short-axis views, and the length of the ventricle was determined, in systole and diastole. These measurements were used to calculate ejection fraction.

### ELISA

Human CNTI ELISA (RayBiotech), IFN-γ (RnD Systems) and BNP ELISA (Abcam) was used as per manufacturer’s instructions.

### Immunoblotting

Protein from 2D hPSC-cardiac cell cultures was extracted using RIPA lysis buffer supplemented with protease inhibitor cocktail (Roche). Protein lysate concentration was estimated using a BCA assay (ThermoFisher Scientific). 15 μg protein was resolved on 4-12% Bis-Tris polyacrylamide gel (Invitrogen) at 200V for 20 min and then transferred at 20 V for 1 h onto polyvinylidene difluoride (PVDF) membrane as per manufacturer’s recommendations. After 1 h blocking using a 1:1 mix of LI-COR Odyssey Blocking Buffer (LI-COR Biotechnology) and PBS, membranes were incubated overnight on a platform shaker with primary antibodies for ACE2 (1:200, R&D Systems, AF933) and GAPDH (1:1000, Cell Signaling Technologies, 97166S). Membranes were washed 5 times 3 minutes in PBS with 0.5% Tween, prior to incubation with IRDye® secondary antibodies (1:10000 for IRDye® 800CW Goat anti-Mouse IgG Secondary Antibody, 926-32210, and 680RD Donkey anti-Goat IgG Secondary Antibody, LI-COR Biotechnology, 925-68074) for 1 h at room temperature. Membranes were washed thoroughly (5 x 3 min in PBS + 0.5% Tween) and were then imaged on a LI-COR Odyssey® CLx. Densitometry was performed using ImageStudio Lite (version 4).

### Flow Cytometry for ACE2 and SARS-CoV-2 spike binding assays

hPSC-cardiac cells were differentiated as above, then plated at 100,000 per cm^2^ on gelatin coated plates and cultured for 5 days prior to infection experiments. For 2D cultured hPSC-CM, cells were pre-treated with MM with or without the indicated compounds or DMSO as a vehicle control for 3 days prior to assays. Cells were was 2 x with PBS and detached using 0.25% Trypsin/EDTA (ThermoFisher Scientific) for ~10-15 min at 37°C. This was then neutralized with equivolume 3% bovine serum albumin (Sigma) in PBS (Binding Buffer). Cells were then centriguged at 300 x g for 3 min and the supernatant removed. Cells were then incubated under different conditions.

For ACE2 assays the following was used for control, 1:200 Goat IgG Alexa Fluor 647-conjugated antibody, and assay 1:200 anti-human ACE2 AlexFluor 647 conjugated antibody and 1:200 anti-human CD90 (all RnD Systems) and were incubated for 60 min at 4°C in Binding Buffer. The cells were then washed in Binding Buffer, centrifuged at 300 x g for 3 min and supernatant removed. Both conditions were then incubated with 1:400 goat anti-mouse IgG secondary antibody conjugated to Alexa Fluor 555 (ThermoFisher Scientific) in Binding Buffer for 45 min at 4°C. The cells were then washed in Binding Buffer, centrifuged at 300 x g for 3 min and supernatant removed.

For SARS-CoV-2 binding assays the following was used for control, Binding Buffer only, and assay 1 μg rSARS‐CoV‐2 Spike RB (per ~100,000 cells) and 1:200 anti-human CD90 (all RnD Systems) and were incubated for 60 min at 4°C in binding buffer. The cells were then washed in Binding Buffer, centrifuged at 300 x g for 3 min and supernatant removed. Both conditions were then incubated with 1:400 F(ab’)2-goat anti-human IgG Fc secondary antibody conjugated to Alexa Fluor 488 and 1:400 goat anti-mouse IgG secondary antibody conjugated to Alexa Fluor 555 (both ThermoFisher Scientific) in Binding Buffer for 45 min at 4°C. The cells were then washed in binding buffer, centrifuged at 300 x g for 3 min and supernatant removed.

For flow cytometry cells were resuspended in 300 μl Binding Buffer, put through a 100 μm cell strainer to remove any clumps. Cells were assessed on a BD LSRFortessa Flow Cytometer, gated on FSC-A/SSC-A and then remove doublets using FSC-W/H, and CD90 assessed using YG586/15, ACE2 using R670/14 and SARS-CoV-2 spike protein B530/30. Negative controls were used to draw gates.

### hPSC-CM SARS-CoV-2 infection at QIMR Berghofer

#### hPSC-cardiac cell infection

hPSC-cardiac cells were differentiated as above, then plated at 100,000 per cm^2^ on gelatin coated plates and cultured for 5 days prior to infection experiments. For 2D cultured hPSC-CM, cells were pre-treated with 1 ml of MM with or without the indicated compounds or DMSO as a vehicle control for 3 days prior to infection. Media was removed before infecting cells with SARS-CoV-2 at MOI 0.01 for 1 h at 37°C. Cells were washed 3 x with MM and replaced with 1 ml of MM with compounds or DMSO. For intracellular RNA cells were washed 3x with PBS before harvesting RNA in Trizol. For supernatant experiments 500 μl of supernatants were harvested each day and replaced with 500 μl of media. Supernatants were frozen at −80°C until they were titred. Titration was performed in Vero cell monolayers in 96 well plates: inoculated serial 10-fold dilutions in quadruplicate and incubated for 4 days. Cytopathic effect recorded and virus titre in log10 TCID_50_/ml recorded.

#### Imaging

For immunostaining cells or hCO were fixed in 4% paraformaldehyde and stained. Cells were stained with primary antibodies Nucleocaspid protein SARS-CoV-2 (1:200, 40143-MM05, Sino Biological) and cardiac troponin T (1:400, ab45932, Abcam) in Blocking Buffer at 4°C for 2 h for 2D or overnight for hCO on a rocker. Cells were washed twice with Blocking Buffer and labelled with secondary antibodies goat anti-mouse IgG AlexaFluor 488 (1:400, A-11001) and goat anti-rabbit IgG AlexaFluor 555 (1:400, A-21428) and Hoechst3332 (all ThermoFisher Scientific) at 4°C for 2 h for 2D or overnight for hCO on a rocker. Cells were again washed with Blocking Buffer twice and then put into PBS and were imaged using a Leica Thunder microscope.

#### Quantitative RT-PCR

Performed as described above.

### hPSC-CM SARS-CoV-2 infection experiments at The Peter Doherty Institute

#### hPSC-CM infection

The human embryonic stem cell line HES3 NKX2-5^eGFP/w^ was used for viral infection studies in 2D monolayer cultures (Elliott et al., 2011). Cardiac cells were differentiated as previously described (Anderson et al., 2018), frozen at day 10 following differentiation and stored at −80°C. Cardiac cells were subsequently thawed in RPMI+B27 media (RPMI 1640 supplemented with 2% B-27 Supplement minus vitamin A, 1% GlutaMAX and 1% Penicillin/Streptomycin- All from ThermoFisher Scientific) with Rock inhibitor (Selleck Chemicals) for 24 h. Cardiac cells were then maintained in RPMI + B27 media for an additional 2 days, enriched for cardiomyocytes with lactate purification media - DMEM, no glucose, no glutamine, no phenol red supplemented with 1% GlutaMAX and 1% Penicillin/Streptomycin (Thermo Fisher Scientific) and 5 mM Sodium L-Lactate (Sigma Aldrich) for 2 days, and subsequently cultured in MM from day 15 to day 23 post differentiation prior to viral infection.

The media was removed from cultures of cardiac myocytes and Vero cells in 24 well plates. Vero cell plates were washed with 1ml of serum free media per well. Media was removed, wells were inoculated with 10^4^ TCID_50_ of virus (MOI = 0.01) in 100 μl serum free media and incubated for 1 h at room temperature. The inoculum for Vero cells incorporated TPCK-treated Trypsin (Worthington Biochemical Corporation) at 1 μg/ml. Following virus adsorption, the inoculum was removed and replaced with 500 μl media. After ~10 minutes, this was removed and stored at −80°C as the day 0 sample. 500 μl per well of media was replenished and plates were incubated at 37°C in 5% CO2. Each day from day 1 through 6 post-infection, 500 μl of culture supernatant was harvested and replenished with 500 μl of media. Supernatants were stored at −80°C till they were titrated.

#### Virus titration

The amount of infectious virus present in the samples was assayed in Vero cell monolayers in 96 well plates. Samples inoculated into 4 wells were diluted serially in 10-fold dilutions in serum free media containing 1 μg/ml TCPK-treated Trypsin and incubated for at 37°C in 5% CO2. Cytopathic effect was scored on day 4 post-infection and virus titre expressed in log_10_ TCID_50_/ml. The lower limit of detection was 1.7 log_10_ TCID_50_/ml.

### SOMAscan™ Proteomic Analysis

SOMAScan™ proteomic technology uses Somamers as an affinity reagent (Somalogic Inc.). Plasma samples from 47 patients from each group that received apabetalone or placebo in the ASSURE trial were analysed and LGALS3BP quantified (Nicholls et al., 2016).

### Data reproducibility and statistical analysis

hCO force experiments were performed on quality controlled hCO (proper formation around the poles, non-arrhythmic, no broken arms, no necking (Mills et al., 2017) across multiple experiments with multiple cell line combinations to ensure reproducibility. Automated force analysis removes the requirement for blinding of hCO experiments. Personnel performing the animal experiments and analyses were blinded to the conditions or treatments. Statistics were performed using GraphPad Prism v8 unless noted.

## REFERENCES

Anand, P., Brown, J.D., Lin, C.Y., Qi, J., Zhang, R., Artero, P.C., Alaiti, M.A., Bullard, J., Alazem, K., Margulies, K.B., et al. (2013). BET bromodomains mediate transcriptional pause release in heart failure. Cell 154, 569–582.

Antolic, A., Wakimoto, H., Jiao, Z., Gorham, J.M., DePalma, S.R., Lemieux, M.E., Conner, D.A., Lee, D.Y., Qi, J., Seidman, J.G., et al. (2020). BET bromodomain proteins regulate transcriptional reprogramming in genetic dilated cardiomyopathy. JCI Insight 5.

Arunachalam, P.S., Wimmers, F., Mok, C.K.P., Perera, R.A.P.M., Scott, M., Hagan, T., Sigal, N., Feng, Y., Bristow, L., Tak-Yin Tsang, O., et al. (2020). Systems biological assessment of immunity to mild versus severe COVID-19 infection in humans. Science 369, 1210–1220.

Auguste, G., Rouhi, L., Matkovich, S.J., Coarfa, C., Robertson, M.J., Czernuszewicz, G., Gurha, P., and Marian, A.J. (2020). BET bromodomain inhibition attenuates cardiac phenotype in myocyte-specific lamin A/C-deficient mice. The Journal of clinical investigation 130, 4740–4758.

Bancerek, J., Poss, Z.C., Steinparzer, I., Sedlyarov, V., Pfaffenwimmer, T., Mikulic, I., Dölken, L., Strobl, B., Müller, M., Taatjes, D.J., et al. (2013). CDK8 kinase phosphorylates transcription factor STAT1 to selectively regulate the interferon response. Immunity 38, 250–262.

Chen, C., Li, H., Hang, W., and Wang, D.W. (2020). Cardiac injuries in coronavirus disease 2019 (COVID-19). Journal of Molecular and Cellular Cardiology 145, 25–29.

Cochran, A.G., Conery, A.R., and Sims, R.J. (2019). Bromodomains: a new target class for drug development. Nature Reviews Drug Discovery 18, 609–628.

Del Valle, D.M., Kim-Schulze, S., Huang, H.-H., Beckmann, N.D., Nirenberg, S., Wang, B., Lavin, Y., Swartz, T.H., Madduri, D., Stock, A., et al. (2020). An inflammatory cytokine signature predicts COVID-19 severity and survival. Nature Medicine.

Duan, Q., McMahon, S., Anand, P., Shah, H., Thomas, S., Salunga, H.T., Huang, Y., Zhang, R., Sahadevan, A., Lemieux, M.E., et al. (2017). BET bromodomain inhibition suppresses innate inflammatory and profibrotic transcriptional networks in heart failure. Science translational medicine 9, eaah5084.

Evron, T., Daigle, T.L., and Caron, M.G. (2012). GRK2: multiple roles beyond G protein-coupled receptor desensitization. Trends Pharmacol Sci 33, 154–164.

Faivre, E.J., McDaniel, K.F., Albert, D.H., Mantena, S.R., Plotnik, J.P., Wilcox, D., Zhang, L., Bui, M.H., Sheppard, G.S., Wang, L., et al. (2020). Selective inhibition of the BD2 bromodomain of BET proteins in prostate cancer. Nature 578, 306–310.

Feldman, A.M., Combes, A., Wagner, D., Kadakomi, T., Kubota, T., Li, Y.Y., and McTiernan, C. (2000). The role of tumor necrosis factor in the pathophysiology of heart failure. Journal of the American College of Cardiology 35, 537–544.

Filippakopoulos, P., Qi, J., Picaud, S., Shen, Y., Smith, W.B., Fedorov, O., Morse, E.M., Keates, T., Hickman, T.T., Felletar, I., et al. (2010). Selective inhibition of BET bromodomains. Nature 468, 1067–1073.

Gagliano-Jucá, T., Travison, T.G., Kantoff, P.W., Nguyen, P.L., Taplin, M.-E., Kibel, A.S., Huang, G., Bearup, R., Schram, H., Manley, R., et al. (2018). Androgen Deprivation Therapy Is Associated With Prolongation of QTc Interval in Men With Prostate Cancer. J Endocr Soc 2, 485–496.

Gilan, O., Rioja, I., Knezevic, K., Bell, M.J., Yeung, M.M., Harker, N.R., Lam, E.Y.N., Chung, C.-w., Bamborough, P., Petretich, M., et al. (2020). Selective targeting of BD1 and BD2 of the BET proteins in cancer and immuno-inflammation. Science, eaaz8455.

Gilsbach, R., Schwaderer, M., Preissl, S., Grüning, B.A., Kranzhöfer, D., Schneider, P., Nührenberg, T.G., Mulero-Navarro, S., Weichenhan, D., Braun, C., et al. (2018). Distinct epigenetic programs regulate cardiac myocyte development and disease in the human heart in vivo. Nature communications 9, 391.

Goyal, P., Choi, J.J., Pinheiro, L.C., Schenck, E.J., Chen, R., Jabri, A., Satlin, M.J., Campion, T.R., Nahid, M., Ringel, J.B., et al. (2020). Clinical Characteristics of Covid-19 in New York City. New England Journal of Medicine 382, 2372–2374.

Guo, S., Carter, R.L., Grisanti, L.A., Koch, W.J., and Tilley, D.G. (2017). Impact of paroxetine on proximal β-adrenergic receptor signaling. Cell Signal 38, 127–133.

Guo, T., Fan, Y., Chen, M., Wu, X., Zhang, L., He, T., Wang, H., Wan, J., Wang, X., and Lu, Z. (2020). Cardiovascular Implications of Fatal Outcomes of Patients With Coronavirus Disease 2019 (COVID-19). JAMA Cardiology 5, 811–818.

Gupta, A., Madhavan, M.V., Sehgal, K., Nair, N., Mahajan, S., Sehrawat, T.S., Bikdeli, B., Ahluwalia, N., Ausiello, J.C., Wan, E.Y., et al. (2020). Extrapulmonary manifestations of COVID-19. Nature Medicine 26, 1017–1032.

Hofmann, M.H., Mani, R., Engelhardt, H., Impagnatiello, M.A., Carotta, S., Kerenyi, M., Lorenzo-Herrero, S., Böttcher, J., Scharn, D., Arnhof, H., et al. (2020). Selective and Potent CDK8/19 Inhibitors Enhance NK-Cell Activity and Promote Tumor Surveillance. Molecular Cancer Therapeutics 19, 1018–1030.

Horby, P., Lim, W.S., Emberson, J., Mafham, M., Bell, J., Linsell, L., Staplin, N., Brightling, C., Ustianowski, A., Elmahi, E., et al. (2020). Effect of Dexamethasone in Hospitalized Patients with COVID-19: Preliminary Report. medRxiv, 2020.2006.2022.20137273.

Huang, C., Wang, Y., Li, X., Ren, L., Zhao, J., Hu, Y., Zhang, L., Fan, G., Xu, J., Gu, X., et al. (2020). Clinical features of patients infected with 2019 novel coronavirus in Wuhan, China. The Lancet 395, 497–506.

Humphrey, S.J., Azimifar, S.B., and Mann, M. (2015). High-throughput phosphoproteomics reveals in vivo insulin signaling dynamics. Nature biotechnology 33, 990–995.

Humphrey, S.J., Karayel, O., James, D.E., and Mann, M. (2018). High-throughput and high-sensitivity phosphoproteomics with the EasyPhos platform. Nature Protocols 13, 1897–1916.

Kim, S.Y., Zhang, X., Schiattarella, G.G., Altamirano, F., Ramos, T.A.R., French, K.M., Jiang, N., Szweda, P.A., Evers, B.M., May, H.I., et al. (2020). Epigenetic Reader BRD4 (Bromodomain-Containing Protein 4) Governs Nucleus-Encoded Mitochondrial Transcriptome to Regulate Cardiac Function. Circulation 142, 2356–2370.

Kubota, T., McTiernan, C.F., Frye, C.S., Slawson, S.E., Lemster, B.H., Koretsky, A.P., Demetris, A.J., and Feldman, A.M. (1997). Dilated Cardiomyopathy in Transgenic Mice With Cardiac-Specific Overexpression of Tumor Necrosis Factor-&#x3b1. Circulation research 81, 627–635.

Levick, S.P., and Goldspink, P.H. (2014). Could interferon-gamma be a therapeutic target for treating heart failure? Heart failure reviews 19, 227–236.

Lindner, D., Fitzek, A., Bräuninger, H., Aleshcheva, G., Edler, C., Meissner, K., Scherschel, K., Kirchhof, P., Escher, F., Schultheiss, H.-P., et al. (2020). Association of Cardiac Infection With SARS-CoV-2 in Confirmed COVID-19 Autopsy Cases. JAMA Cardiology.

Mangalmurti, N., and Hunter, C.A. (2020). Cytokine Storms: Understanding COVID-19. Immunity 53, 19–25.

Messner, C.B., Demichev, V., Wendisch, D., Michalick, L., White, M., Freiwald, A., Textoris-Taube, K., Vernardis, S.I., Egger, A.S., Kreidl, M., et al. (2020). Ultra-High-Throughput Clinical Proteomics Reveals Classifiers of COVID-19 Infection. Cell systems 11, 11–24.e14.

Mills, R.J., Parker, B.L., Quaife-Ryan, G.A., Voges, H.K., Needham, E.J., Bornot, A., Ding, M., Andersson, H., Polla, M., Elliott, D.A., et al. (2019). Drug Screening in Human PSC-Cardiac Organoids Identifies Pro-proliferative Compounds Acting via the Mevalonate Pathway. Cell stem cell 24, 895–907.e896.

Mills, R.J., Titmarsh, D.M., Koenig, X., Parker, B.L., Ryall, J.G., Quaife-Ryan, G.A., Voges, H.K., Hodson, M.P., Ferguson, C., Drowley, L., et al. (2017). Functional screening in human cardiac organoids reveals a metabolic mechanism for cardiomyocyte cell cycle arrest. Proceedings of the National Academy of Sciences 114, E8372–E8381.

Needham, E.J., Parker, B.L., Burykin, T., James, D.E., and Humphrey, S.J. (2019). Illuminating the dark phosphoproteome. Science Signaling 12, eaau8645.

Nicholls, S.J., Schwartz, G.G., Buhr, K.A., Ginsberg, H.N., Johansson, J.O., Kalantar-Zadeh, K., Kulikowski, E., Toth, P.P., Wong, N., Sweeney, M., et al. (2021). Apabetalone and hospitalization for heart failure in patients following an acute coronary syndrome: a prespecified analysis of the BETonMACE study. Cardiovascular diabetology 20, 13.

Nicodeme, E., Jeffrey, K.L., Schaefer, U., Beinke, S., Dewell, S., Chung, C.W., Chandwani, R., Marazzi, I., Wilson, P., Coste, H., et al. (2010). Suppression of inflammation by a synthetic histone mimic. Nature 468, 1119–1123.

Nishiga, M., Wang, D.W., Han, Y., Lewis, D.B., and Wu, J.C. (2020). COVID-19 and cardiovascular disease: from basic mechanisms to clinical perspectives. Nature Reviews Cardiology.

Oladunni, F.S., Park, J.G., Pino, P.A., Gonzalez, O., Akhter, A., Allué-Guardia, A., Olmo-Fontánez, A., Gautam, S., Garcia-Vilanova, A., Ye, C., et al. (2020). Lethality of SARS-CoV-2 infection in K18 human angiotensin-converting enzyme 2 transgenic mice. Nature communications 11, 6122.

Padmanabhan, A., Alexanian, M., Linares-Saldana, R., González-Terán, B., Andreoletti, G., Huang, Y., Connolly, A.J., Kim, W., Hsu, A., Duan, Q., et al. (2020). BRD4 (Bromodomain-Containing Protein 4) Interacts with GATA4 (GATA Binding Protein 4) to Govern Mitochondrial Homeostasis in Adult Cardiomyocytes. Circulation 142, 2338–2355.

Pellegrini, D., Kawakami, R., Guagliumi, G., Sakamoto, A., Kawai, K., Gianatti, A., Nasr, A., Kutys, R., Guo, L., Cornelissen, A., et al. (2021). Microthrombi As A Major Cause of Cardiac Injury in COVID-19: A Pathologic Study. Circulation.

Puntmann, V.O., Carerj, M.L., Wieters, I., Fahim, M., Arendt, C., Hoffmann, J., Shchendrygina, A., Escher, F., Vasa-Nicotera, M., Zeiher, A.M., et al. (2020). Outcomes of Cardiovascular Magnetic Resonance Imaging in Patients Recently Recovered From Coronavirus Disease 2019 (COVID-19). JAMA Cardiology.

Qiao, Y., Wang, X.-M., Mannan, R., Pitchiaya, S., Zhang, Y., Wotring, J.W., Xiao, L., Robinson, D.R., Wu, Y.-M., Tien, J.C.-Y., et al. (2021). Targeting transcriptional regulation of SARS-CoV-2 entry factors ACE2 and TMPRSS2. Proceedings of the National Academy of Sciences 118, e2021450118.

Quaife-Ryan, G.A., Sim, C.B., Ziemann, M., Kaspi, A., Rafehi, H., Ramialison, M., El-Osta, A., Hudson, J.E., and Porrello, E.R. (2017). Multicellular Transcriptional Analysis of Mammalian Heart Regeneration. Circulation 136, 1123–1139.

Ray, K.K., Nicholls, S.J., Buhr, K.A., Ginsberg, H.N., Johansson, J.O., Kalantar-Zadeh, K., Kulikowski, E., Toth, P.P., Wong, N., Sweeney, M., et al. (2020). Effect of Apabetalone Added to Standard Therapy on Major Adverse Cardiovascular Events in Patients With Recent Acute Coronary Syndrome and Type 2 Diabetes: A Randomized Clinical Trial. Jama 323, 1565–1573.

Ren, X., Wen, W., Fan, X., Hou, W., Su, B., Cai, P., Li, J., Liu, Y., Tang, F., Zhang, F., et al. (2021). COVID-19 immune features revealed by a large-scale single cell transcriptome atlas. Cell.

Richardson, P., Griffin, I., Tucker, C., Smith, D., Oechsle, O., Phelan, A., Rawling, M., Savory, E., and Stebbing, J. (2020). Baricitinib as potential treatment for 2019-nCoV acute respiratory disease. Lancet 395, e30–e31.

Runte, K.E., Bell, S.P., Selby, D.E., Häußler, T.N., Ashikaga, T., LeWinter, M.M., Palmer, B.M., and Meyer, M. (2017). Relaxation and the Role of Calcium in Isolated Contracting Myocardium From Patients With Hypertensive Heart Disease and Heart Failure With Preserved Ejection Fraction. Circ Heart Fail 10.

Rzymski, T., Mikula, M., Żyłkiewicz, E., Dreas, A., Wiklik, K., Gołas, A., Wójcik, K., Masiejczyk, M., Wróbel, A., Dolata, I., et al. (2017). SEL120-34A is a novel CDK8 inhibitor active in AML cells with high levels of serine phosphorylation of STAT1 and STAT5 transactivation domains. Oncotarget 8, 33779–33795.

Sadzak, I., Schiff, M., Gattermeier, I., Glinitzer, R., Sauer, I., Saalmüller, A., Yang, E., Schaljo, B., and Kovarik, P. (2008). Recruitment of Stat1 to chromatin is required for interferon-induced serine phosphorylation of Stat1 transactivation domain. Proceedings of the National Academy of Sciences of the United States of America 105, 8944–8949.

Sanders-van Wijk, S., Tromp, J., Beussink-Nelson, L., Hage, C., Svedlund, S., Saraste, A., Swat, S.A., Sanchez, C., Njoroge, J., Tan, R.S., et al. (2020). Proteomic Evaluation of the Comorbidity-Inflammation Paradigm in Heart Failure With Preserved Ejection Fraction: Results From the PROMIS-HFpEF Study. Circulation 142, 2029–2044.

Schumacher, S.M., Gao, E., Zhu, W., Chen, X., Chuprun, J.K., Feldman, A.M., Tesmer, J.J.G., and Koch, W.J. (2015). Paroxetine-mediated GRK2 inhibition reverses cardiac dysfunction and remodeling after myocardial infarction. Science translational medicine 7, 277ra231–277ra231.

Sharma, A., Garcia, G., Jr., Wang, Y., Plummer, J.T., Morizono, K., Arumugaswami, V., and Svendsen, C.N. (2020). Human iPSC-Derived Cardiomyocytes Are Susceptible to SARS-CoV-2 Infection. Cell Rep Med 1, 100052–100052.

Shi, S., Qin, M., Shen, B., Cai, Y., Liu, T., Yang, F., Gong, W., Liu, X., Liang, J., Zhao, Q., et al. (2020). Association of Cardiac Injury With Mortality in Hospitalized Patients With COVID-19 in Wuhan, China. JAMA Cardiology 5, 802–810.

Stratton, M.S., Bagchi, R.A., Felisbino, M.B., Hirsch, R.A., Smith, H.E., Riching, A.S., Enyart, B.Y., Koch, K.A., Cavasin, M.A., Alexanian, M., et al. (2019). Dynamic Chromatin Targeting of BRD4 Stimulates Cardiac Fibroblast Activation. Circulation research 125, 662–677.

Stubbs, M.C., Burn, T.C., Sparks, R., Maduskuie, T., Diamond, S., Rupar, M., Wen, X., Volgina, A., Zolotarjova, N., Waeltz, P., et al. (2019). The Novel Bromodomain and Extraterminal Domain Inhibitor INCB054329 Induces Vulnerabilities in Myeloma Cells That Inform Rational Combination Strategies. Clinical Cancer Research 25, 300–311.

Szekely, Y., Lichter, Y., Taieb, P., Banai, A., Hochstadt, A., Merdler, I., Oz, A.G., Rothschild, E., Baruch, G., Peri, Y., et al. (2020). The Spectrum of Cardiac Manifestations in Coronavirus Disease 2019 (COVID-19) - a Systematic Echocardiographic Study. Circulation 0.

Tucker, N.R., Chaffin, M., Fleming, S.J., Hall, A.W., Parsons, V.A., Jr, K.C.B., Akkad, A.-D., Herndon, C.N., Arduini, A., Papangeli, I., et al. (2020). Transcriptional and Cellular Diversity of the Human Heart. Circulation 0.

Vagnozzi, R.J., Gatto, G.J., Jr., Kallander, L.S., Hoffman, N.E., Mallilankaraman, K., Ballard, V.L.T., Lawhorn, B.G., Stoy, P., Philp, J., Graves, A.P., et al. (2013). Inhibition of the cardiomyocyte-specific kinase TNNI3K limits oxidative stress, injury, and adverse remodeling in the ischemic heart. Science translational medicine 5, 207ra141–207ra141.

Vasudevan, N.T., Mohan, M.L., Gupta, M.K., Martelli, E.E., Hussain, A.K., Qin, Y., Chandrasekharan, U.M., Young, D., Feldman, A.M., Sen, S., et al. (2013). Gβγ-independent recruitment of G-protein coupled receptor kinase 2 drives tumor necrosis factor α-induced cardiac β-adrenergic receptor dysfunction. Circulation 128, 377–387.

Voges, H.K., Mills, R.J., Elliott, D.A., Parton, R.G., Porrello, E.R., and Hudson, J.E. (2017). Development of a human cardiac organoid injury model reveals innate regenerative potential. Development (Cambridge, England) 144, 1118–1127.

Williams, L.M., McCann, F.E., Cabrita, M.A., Layton, T., Cribbs, A., Knezevic, B., Fang, H., Knight, J., Zhang, M., Fischer, R., et al. (2020). Identifying collagen VI as a target of fibrotic diseases regulated by CREBBP/EP300. Proceedings of the National Academy of Sciences of the United States of America 117, 20753–20763.

Wu, Z., and McGoogan, J.M. (2020). Characteristics of and Important Lessons From the Coronavirus Disease 2019 (COVID-19) Outbreak in China: Summary of a Report of 72⍰314 Cases From the Chinese Center for Disease Control and Prevention. JAMA 323, 1239–1242.

## REFERENCES

Anderson, D.J., Kaplan, D.I., Bell, K.M., Koutsis, K., Haynes, J.M., Mills, R.J., Phelan, D.G., Qian, E.L., Leitoguinho, A.R., Arasaratnam, D., et al. (2018). NKX2-5 regulates human cardiomyogenesis via a HEY2 dependent transcriptional network. Nature communications 9, 1373.

Becht, E., McInnes, L., Healy, J., Dutertre, C.-A., Kwok, I.W.H., Ng, L.G., Ginhoux, F., and Newell, E.W. (2019). Dimensionality reduction for visualizing single-cell data using UMAP. Nature biotechnology 37, 38–44.

Cox, J., and Mann, M. (2008). MaxQuant enables high peptide identification rates, individualized p.p.b.-range mass accuracies and proteome-wide protein quantification. Nature biotechnology 26, 1367–1372.

DeLuca, D.S., Levin, J.Z., Sivachenko, A., Fennell, T., Nazaire, M.D., Williams, C., Reich, M., Winckler, W., and Getz, G. (2012). RNA-SeQC: RNA-seq metrics for quality control and process optimization. Bioinformatics (Oxford, England) 28, 1530–1532.

Deutsch, E.W., Csordas, A., Sun, Z., Jarnuczak, A., Perez-Riverol, Y., Ternent, T., Campbell, D.S., Bernal-Llinares, M., Okuda, S., Kawano, S., et al. (2017). The ProteomeXchange consortium in 2017: supporting the cultural change in proteomics public data deposition. Nucleic Acids Res 45, D1100–d1106.

Dobin, A., Davis, C.A., Schlesinger, F., Drenkow, J., Zaleski, C., Jha, S., Batut, P., Chaisson, M., and Gingeras, T.R. (2013). STAR: ultrafast universal RNA-seq aligner. Bioinformatics (Oxford, England) 29, 15–21.

Elliott, D.A., Braam, S.R., Koutsis, K., Ng, E.S., Jenny, R., Lagerqvist, E.L., Biben, C., Hatzistavrou, T., Hirst, C.E., Yu, Q.C., et al. (2011). NKX2-5(eGFP/w) hESCs for isolation of human cardiac progenitors and cardiomyocytes. Nature methods 8, 1037–1040.

Huber, W., Carey, V.J., Gentleman, R., Anders, S., Carlson, M., Carvalho, B.S., Bravo, H.C., Davis, S., Gatto, L., Girke, T., et al. (2015). Orchestrating high-throughput genomic analysis with Bioconductor. Nature methods 12, 115–121.

Hudson, J., Titmarsh, D., Hidalgo, A., Wolvetang, E., and Cooper-White, J. (2012). Primitive cardiac cells from human embryonic stem cells. Stem cells and development 21, 1513–1523.

Johnson, B.J., Le, T.T., Dobbin, C.A., Banovic, T., Howard, C.B., Flores Fde, M., Vanags, D., Naylor, D.J., Hill, G.R., and Suhrbier, A. (2005). Heat shock protein 10 inhibits lipopolysaccharide-induced inflammatory mediator production. The Journal of biological chemistry 280, 4037–4047.

Kuleshov, M.V., Jones, M.R., Rouillard, A.D., Fernandez, N.F., Duan, Q., Wang, Z., Koplev, S., Jenkins, S.L., Jagodnik, K.M., Lachmann, A., et al. (2016). Enrichr: a comprehensive gene set enrichment analysis web server 2016 update. Nucleic acids research 44, W90–W97.

La Linn, M., Bellett, A.J., Parsons, P.G., and Suhrbier, A. (1995). Complete removal of mycoplasma from viral preparations using solvent extraction. J Virol Methods 52, 51–54.

Li, B., and Dewey, C.N. (2011). RSEM: accurate transcript quantification from RNA-Seq data with or without a reference genome. BMC Bioinformatics 12, 323.

Li, H., Handsaker, B., Wysoker, A., Fennell, T., Ruan, J., Homer, N., Marth, G., Abecasis, G., and Durbin, R. (2009). The Sequence Alignment/Map format and SAMtools. Bioinformatics (Oxford, England) 25, 2078–2079.

Liao, Y., Smyth, G.K., and Shi, W. (2014). featureCounts: an efficient general purpose program for assigning sequence reads to genomic features. Bioinformatics (Oxford, England) 30, 923–930.

Martin, M. (2011). Cutadapt removes adapter sequences from high-throughput sequencing reads. 2011 17, 3.

Nicholls, S.J., Puri, R., Wolski, K., Ballantyne, C.M., Barter, P.J., Brewer, H.B., Kastelein, J.J., Hu, B., Uno, K., Kataoka, Y., et al. (2016). Effect of the BET Protein Inhibitor, RVX-208, on Progression of Coronary Atherosclerosis: Results of the Phase 2b, Randomized, Double-Blind, Multicenter, ASSURE Trial. American journal of cardiovascular drugs: drugs, devices, and other interventions 16, 55–65.

Orlova, V.V., van den Hil, F.E., Petrus-Reurer, S., Drabsch, Y., Ten Dijke, P., and Mummery, C.L. (2014). Generation, expansion and functional analysis of endothelial cells and pericytes derived from human pluripotent stem cells. Nature protocols 9, 1514–1531.

Robinson, M.D., McCarthy, D.J., and Smyth, G.K. (2010). edgeR: a Bioconductor package for differential expression analysis of digital gene expression data. Bioinformatics (Oxford, England) 26, 139–140.

Tyanova, S., and Cox, J. (2018). Perseus: A Bioinformatics Platform for Integrative Analysis of Proteomics Data in Cancer Research. Methods in molecular biology (Clifton, NJ) 1711, 133–148.

Wolf, F.A., Angerer, P., and Theis, F.J. (2018). SCANPY: large-scale single-cell gene expression data analysis. Genome biology 19, 15.

