## Supplementary material for "Bromodomain and Extraterminal Inhibition Blocks Inflammation-Induced Cardiac Dysfunction and SARS-CoV-2 Infection (Pre-Clinical)": Mills et al Supplementary Data Biorxiv

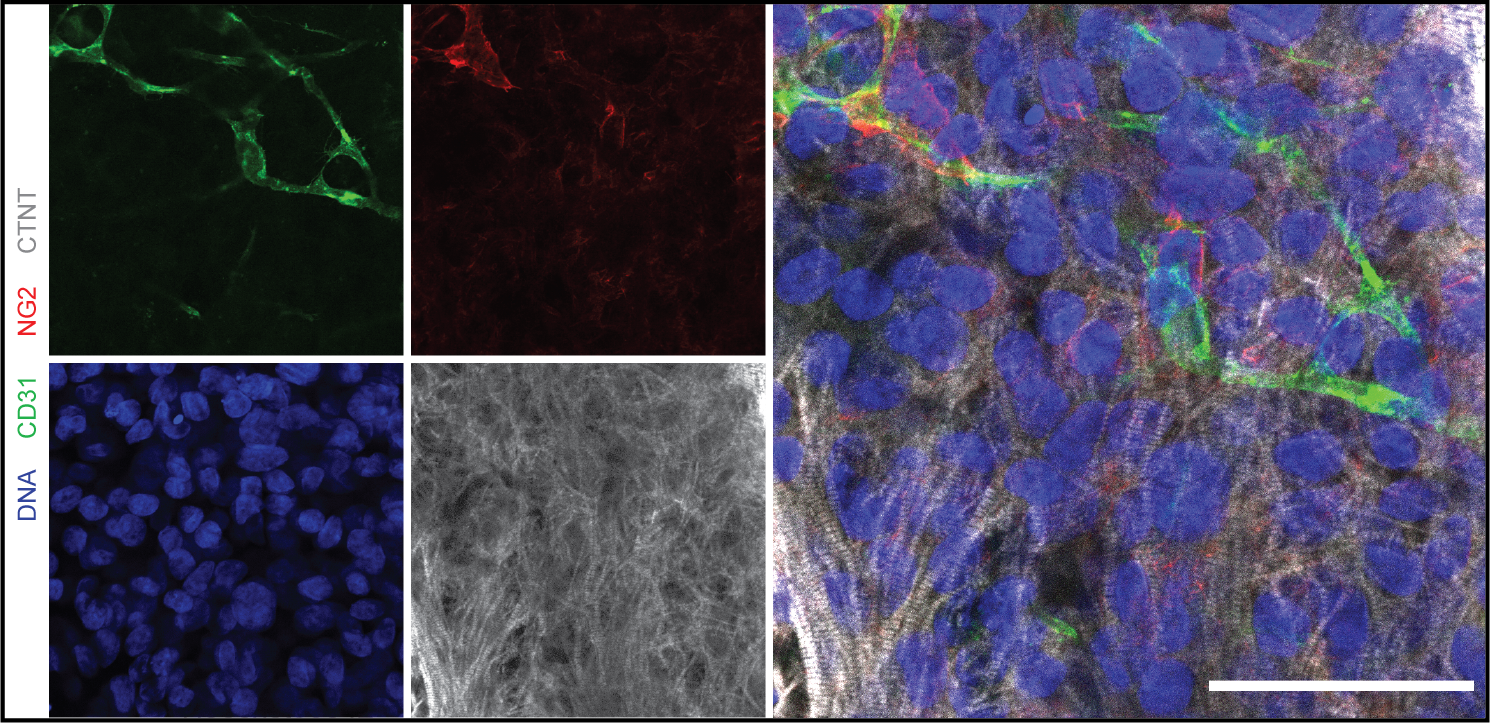

**Supplementary Figure S1: Presence of endothelial structures supported by pericytes in enhanced hCO.** Whole-mount immunofluorescent images of hCO stained with CD31 (endothelial cells), NG2 (pericytes), cardiac troponin T (cardiomyocytes) and Hoescht33342. Scale = 50 µm.

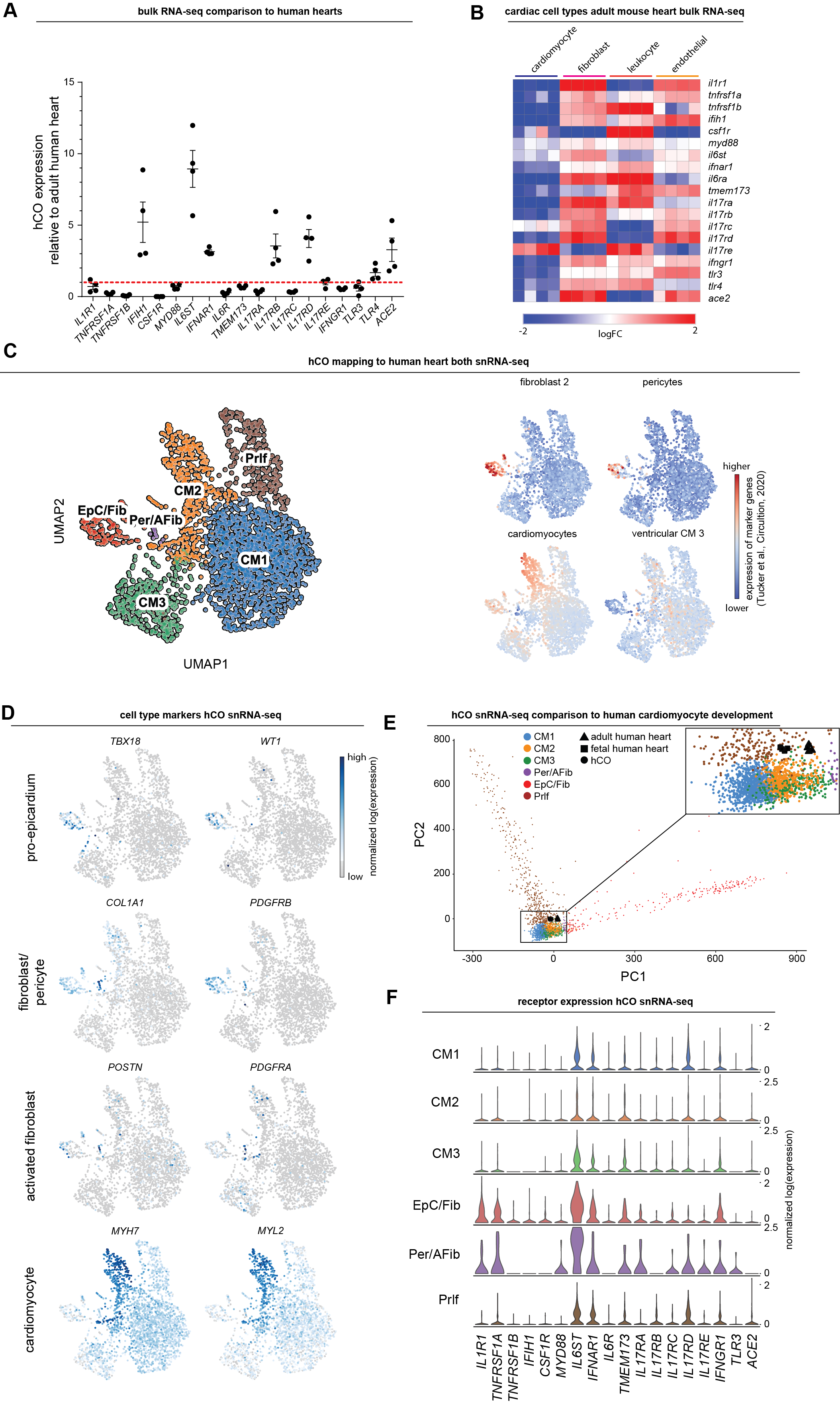

**Supplementary Figure S2:** **Expression of immunomodulatory receptors and signalling mediators.**

A) Comparison of immunomodulatory receptors and signalling mediators in hCO relative to human adult heart using existing bulk RNA-sequencing data (n = 4 experiments for hCO) (*1*). All were identified in hCO except CSF1R which is leukocyte specific.
B) Cell type specificity of immunomodulatory receptors and signalling mediators in adult mouse hearts using existing bulk RNA sequencing data (n = 4 experiments) (*2*).
C) Normalized expression of genes in hCO that mark the human heart sub-populations defined in a recent publication (*3*).
D) UMAP clustering of snRNA-seq of hCO using the new enhanced protocol (Voges et al., In Preparation). Location of key markers for different cell populations are also highlighted.
E) Principal component analysis of our snRNA-seq in comparison to purified bulk RNA-seq of purified human cardiomyocyte nuceli (*4*).
F) Expression of immunomodulatory receptors and signalling mediators in different cell populations in the enhanced hCO.

hPSC cardiac cells- AA, Endothelial cells- RM3.5.

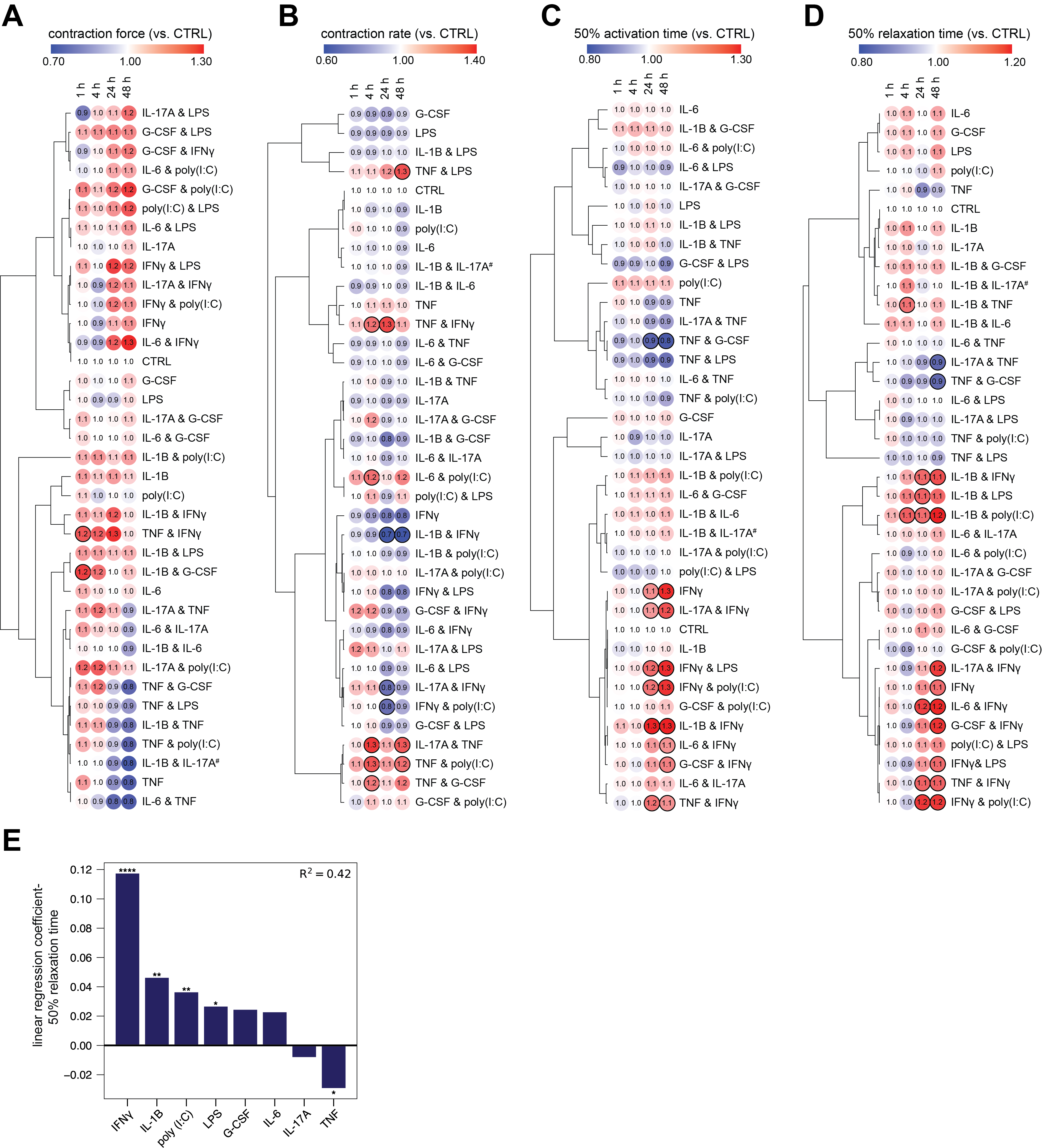

**Supplementary Figure S3:** **Pair-wise screening for the impact of pro-inflammatory factors on hCO function.**

A) Contraction force
B) Contraction rate
C) Time from 50% activation to peak
D) Time from peak to 50% relaxation

hCO normalized to baseline contraction parameters at 0 hr. ^#^ Due to a microscope camera shutter issue at the 0 h time point, IL-1β & IL-17A is normalized to the 1 h time point. Bold outline indicates p <0.05 using a one-way ANOVA with Dunnett's multiple comparisons test comparing each condition to CTRL at comparable time point. n= 2-5 hCOs per condition for cytokine treatments, n= 7-9 hCOs for CTRL from 1 experiment. hPSC cardiac cells- AA, Endothelial cells- RM3.5.

E) Coefficients of linear regression performed using binary predictors (cytokine presence/absence) with time from peak to 50% relaxation as the outcome variable at 24 h. Coefficients represent the mean change in the response given one unit change in the predictor. Sign of the coefficient represents the direction of the change between predictor and response. Overall, presence of IFN-γ, IL-1β, poly(I:C) all lead to increased relaxation times whilst presence of TNF leads to a reduced relaxation. * p <0.05, ** p<0.01, **** p<0.0001 using regression modelling.

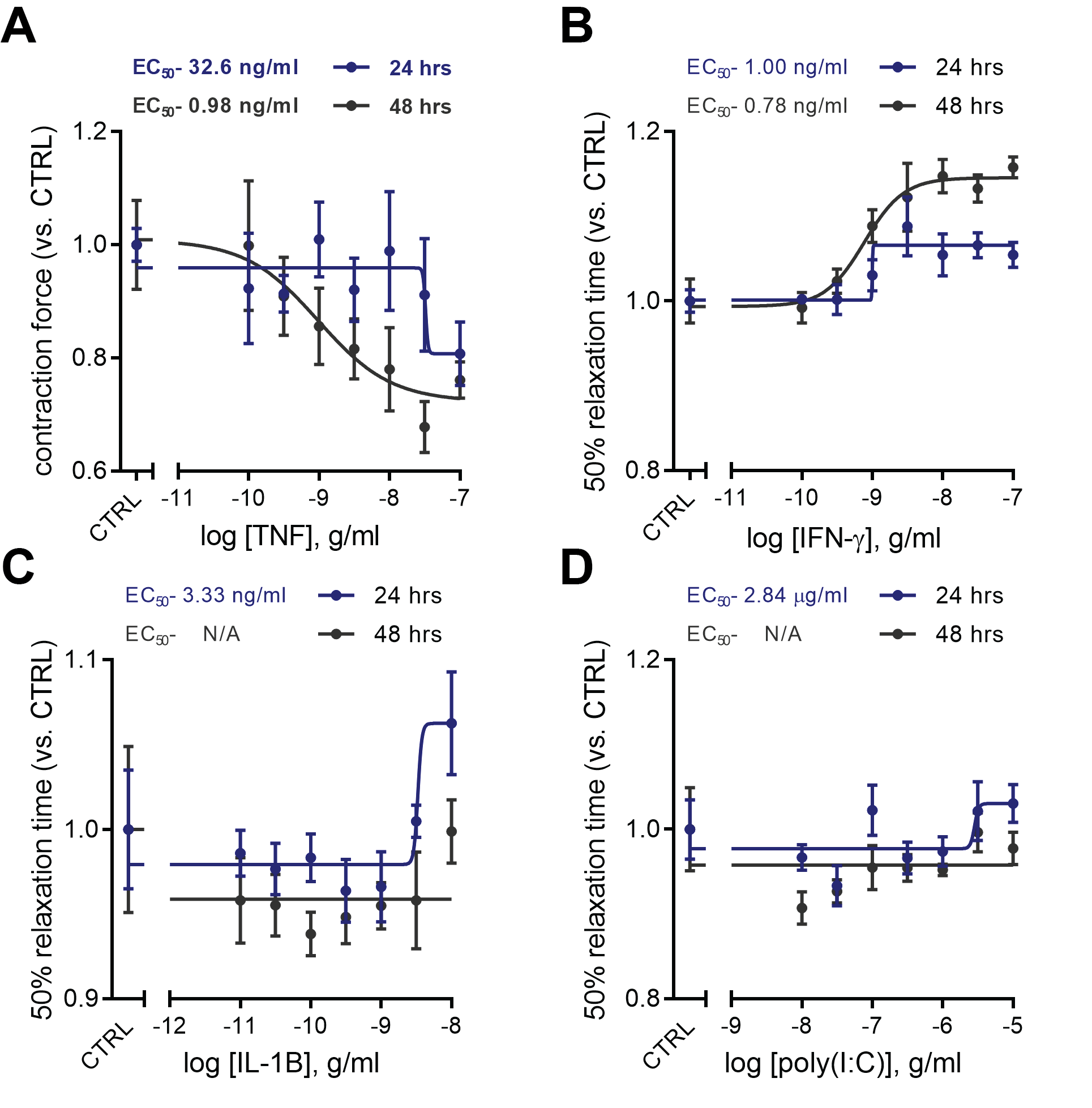

**Supplementary Figure S4: Dose-response curves of key inflammatory mediators for hCO dysfunction.**

A) Dose-response curves for TNF, force of contraction
B) Dose-response curves for IFN-γ, time to 50% relaxation
C) Dose-response curves for IL-1β, time to 50% relaxation
D) Dose-response curves for poly(I:C), time to 50% relaxation

n= 4-5 hCOs per concentration, per condition from 1 experiment. hiPSC cardiac cells- HES3, Endothelial cells- RM3.5. Dose-response curve was generated using nonlinear regression (variable slope model, sigmoidal- 4 parameter logistic) to determine cytokine EC_50_.

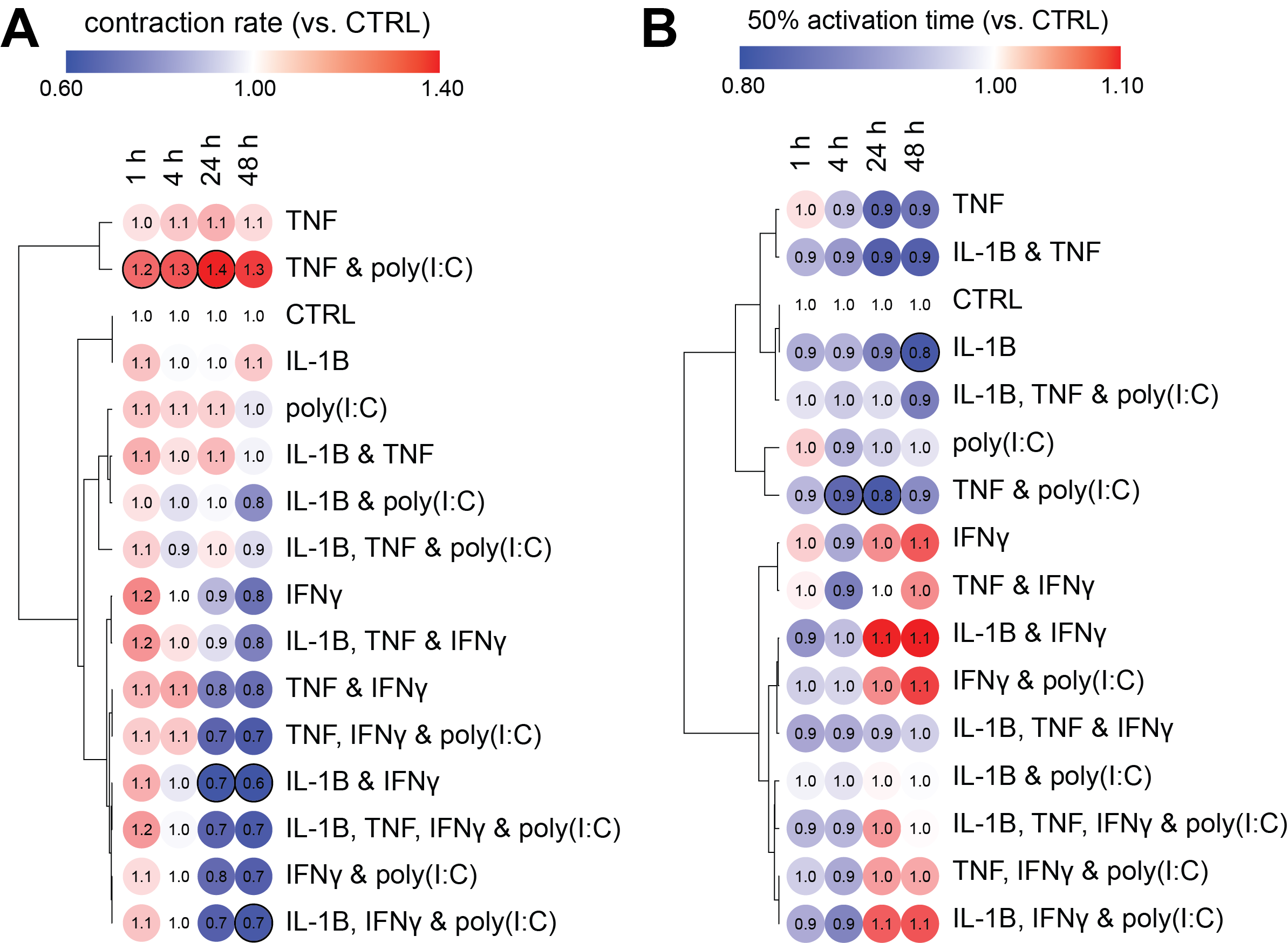

**Supplementary Figure S5: Additional parameters of pro-inflammatory factors driving hCO dysfunction.**

A) Contraction rate
B) Time from 50% activation to peak

Bold outline indicates p <0.05 using one-way ANOVA with Dunnett's multiple comparisons test comparing each condition to CTRL at its’ time point. n= 3-5 hCOs per condition from 1 experiment. hPSC cardiac cells- AA, Endothelial cells- RM3.5. Functional parameters relate to **Fig 1B,C**.

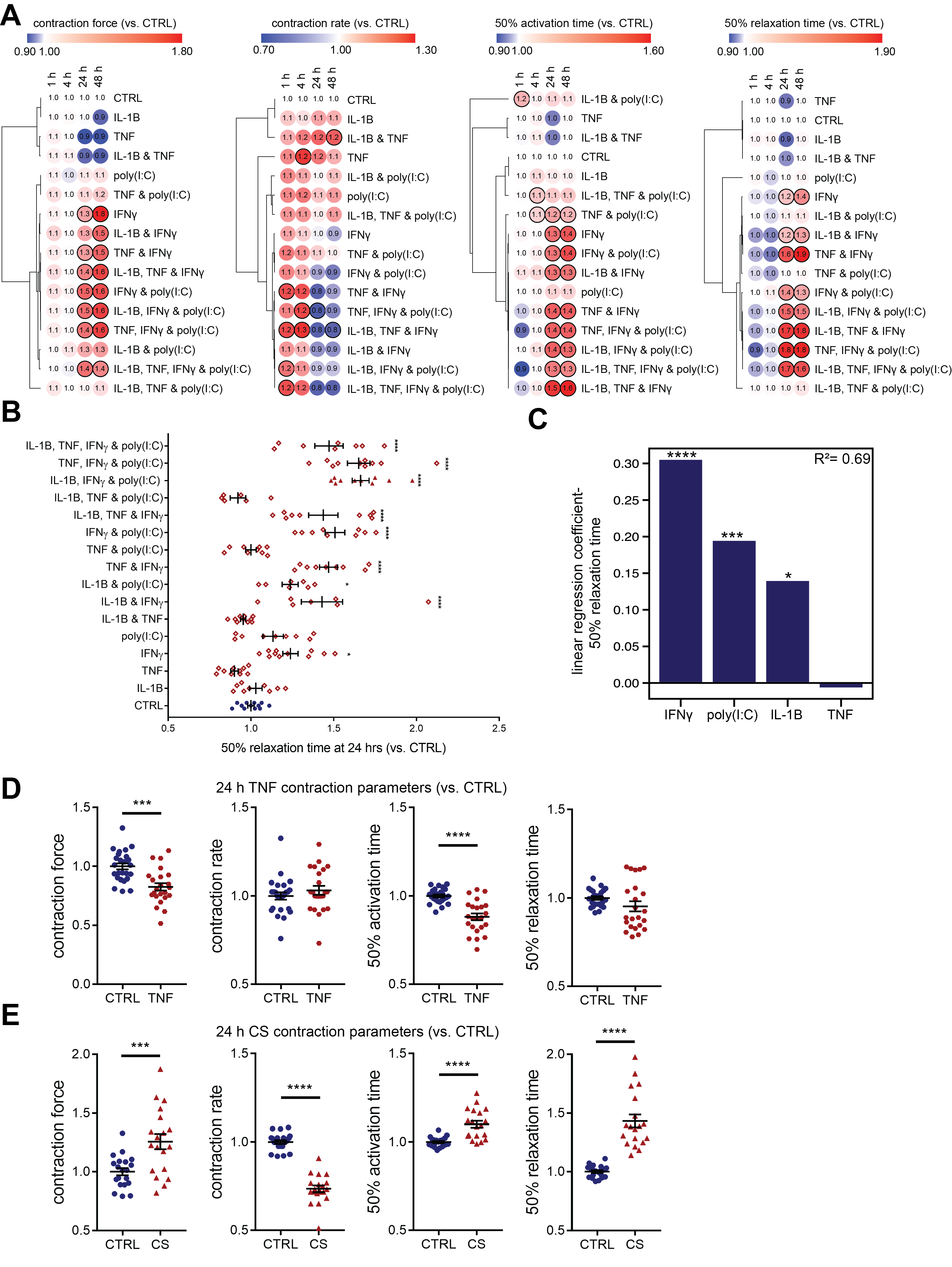

**Supplementary Figure S6: Validation of pro-inflammatory factor screen in an additional cell line and regression analysis.**

A) Validation of functional inflammatory modulator screening parameters in an additional cell line. Bold outline indicates p <0.05 using one-way ANOVA with Dunnett's multiple comparisons test comparing each condition to CTRL at its’ time point. n= 2-6 hCOs per cytokine treatment and, n= 7 hCOs for CTRL from 1 experiment. hPSC cardiac cells - HES3, Endothelial cells - CC.
B) Overall impact of inflammatory modulators on time to 50% relaxation (diastolic function) at 24 h for both hPSC lines tested. n= 6-12 hCOs per condition from 2 experiments. hPSC cardiac cells- AA and HES3, Endothelial cells- RM3.5. * p <0.05, **** p<0.0001, using one-way ANOVA with Dunnett's multiple comparisons test comparing each condition to CTRL.
C) Coefficients of linear regression performed (order = 2) using binary predictors (cytokine presence/absence) with relaxation time as the outcome variable. Coefficients represent the mean change in the response given one unit change in the predictor. Sign of the coefficient represents the direction of the change between predictor and response. The presence of IFN-γ, poly(I:C) and IL-1β lead to increased time to 50% relaxation. * p <0.05, *** p<0.001, **** p<0.0001 using regression modelling.
D) Validation of TNF systolic dysfunction in an additional cell line. n = 25 and 23 hCOs for CTRL and TNF conditions, respectively from 3 experiments. hPSCs cardiac cells- AA, Endothelial cells- RM3.5 and CC. *** p<0.001, **** p<0.0001, using Student’s t-test.
E) Validation of CS induced diastolic dysfunction in an additional cell line. n = 20 and 19 hCOs for CTRL and CS conditions, respectively from 3 experiments. hPSCs cardiac cells- AA, Endothelial cells- RM3.5 and CC. *** p<0.001, **** p<0.0001, using Student’s t-test.

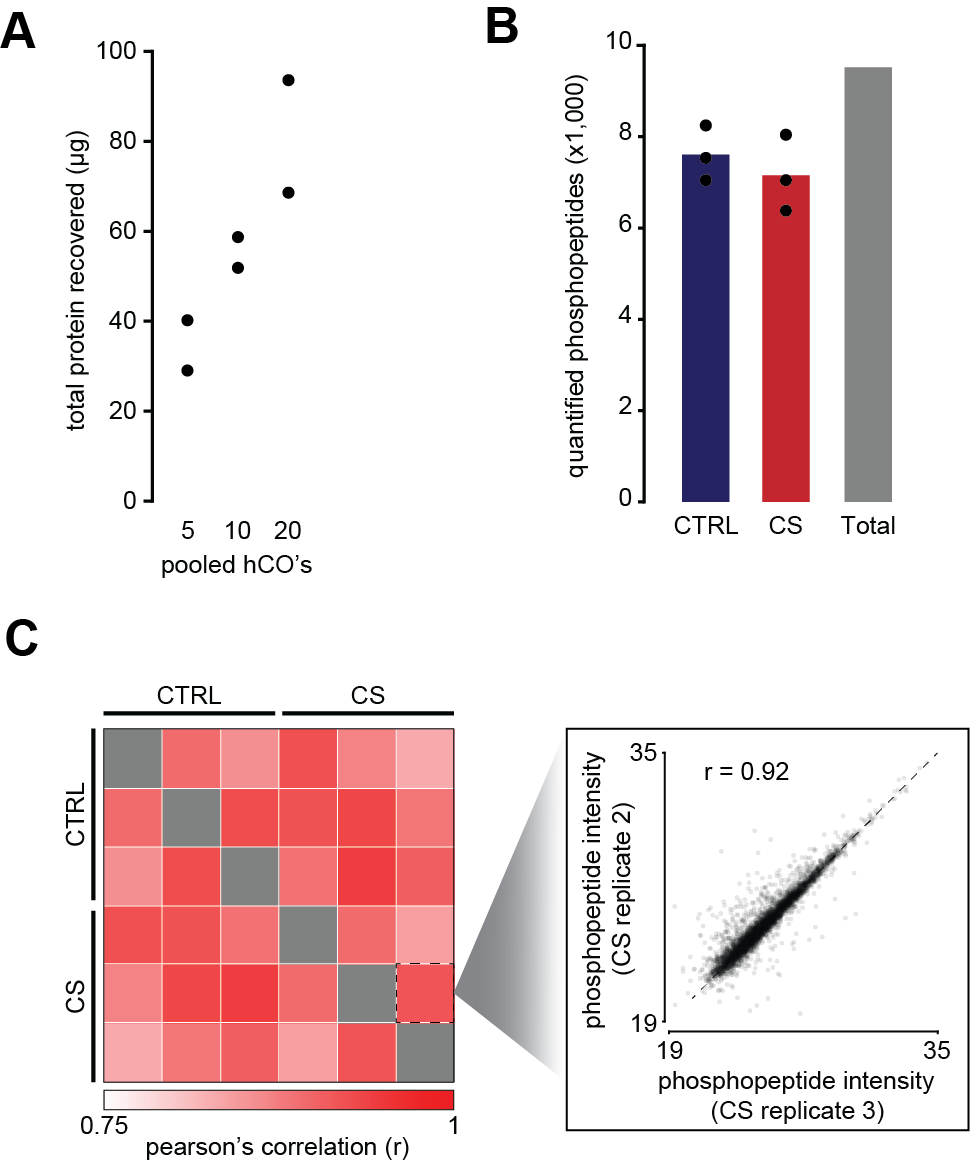

**Supplementary Figure S7: Phosphoproteomic sample quality control.**

A) Recovered protein from different numbers of pooled hCO.
B) Quantified phosphopeptides.
C) Correlation of replicate samples.

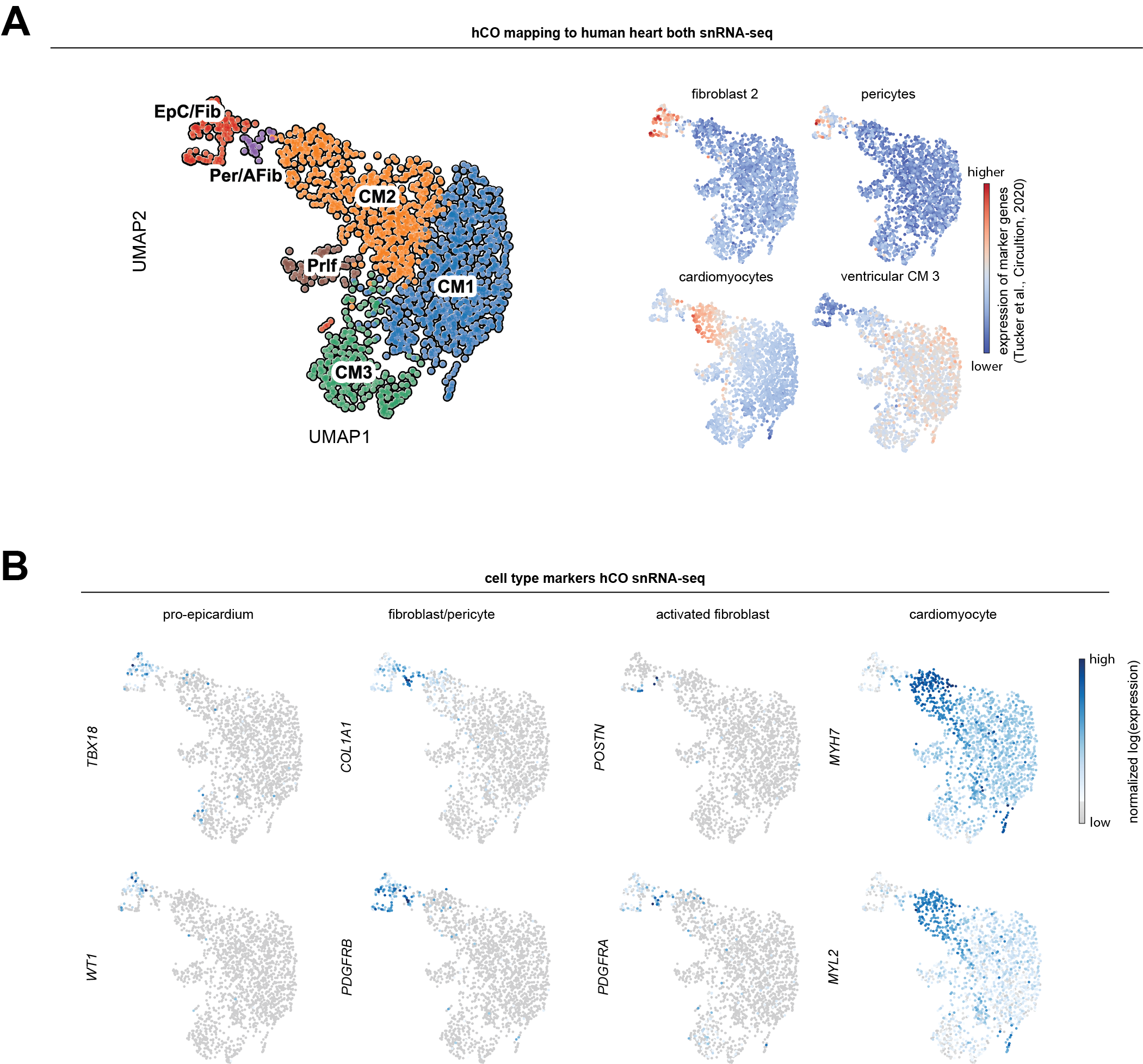

**Supplementary Figure S8: snRNA-seq populations in hCO treated with cardiac cytokine storm.**

A) Normalized expression of genes in hCO that mark the human heart sub-populations defined in a recent publication (*3*).
B) UMAP clustering of snRNA-seq of CS treated hCO. Location of key markers for different cell populations are also highlighted.

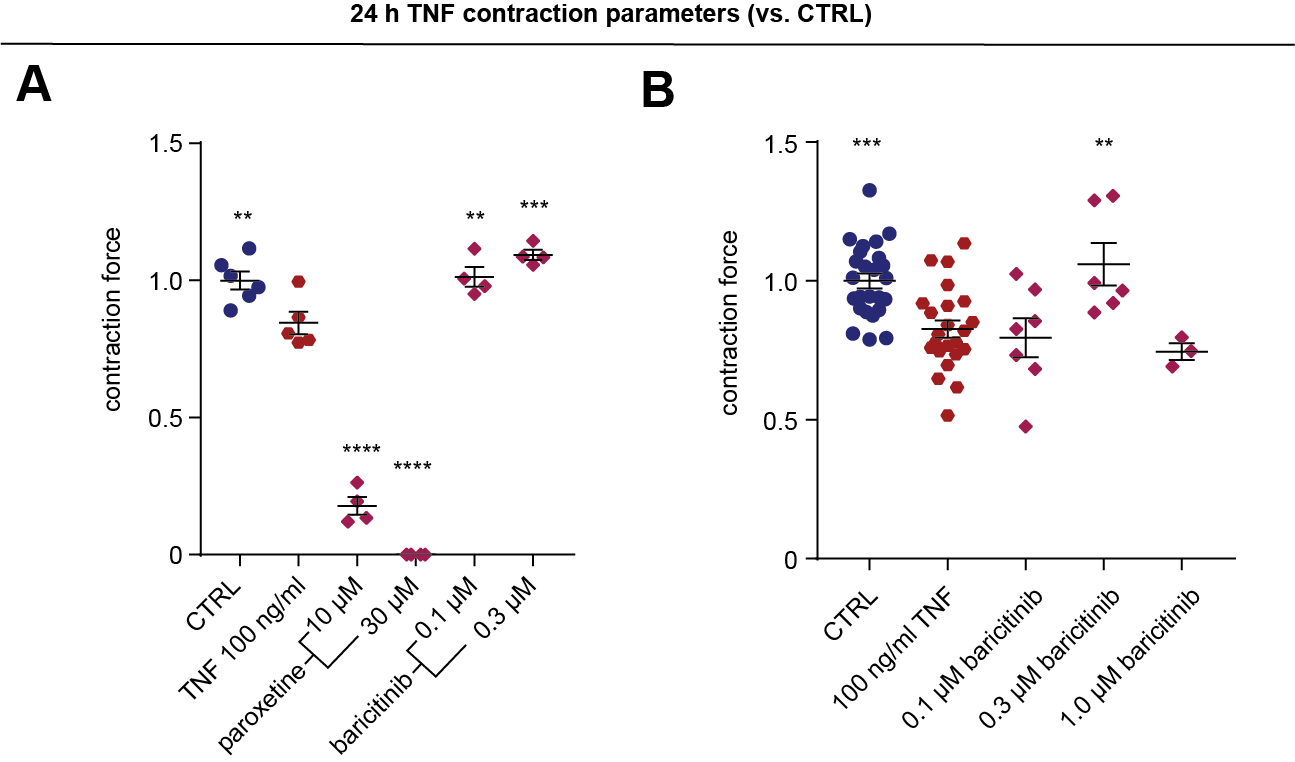

**Supplementary Figure S9: hCO protection from TNF-induced systolic dysfunction.**

A) hCOs were concurrently treated with 100 ng/ml TNF and inhibitors and then functionally assessed at 24 h. n = 4-6 hCOs per condition from 1 experiment, hPSC cardiac cells- HES3, Endothelial cells- RM3.5.
B) Validation in an additional cell line. hCOs were concurrently treated with 100 ng/ml TNF and inhibitors and then functionally assessed at 24 h. n = 3-23 hCOs per condition from 1-2 experiments, hPSCs-cardiac cells - AA, Endothelial cells - CC.

All compounds are in the presence of TNF. ** p<0.01, *** p<0.001, **** p<0.0001, using a one-way ANOVA with Dunnett's multiple comparisons test compared to TNF.

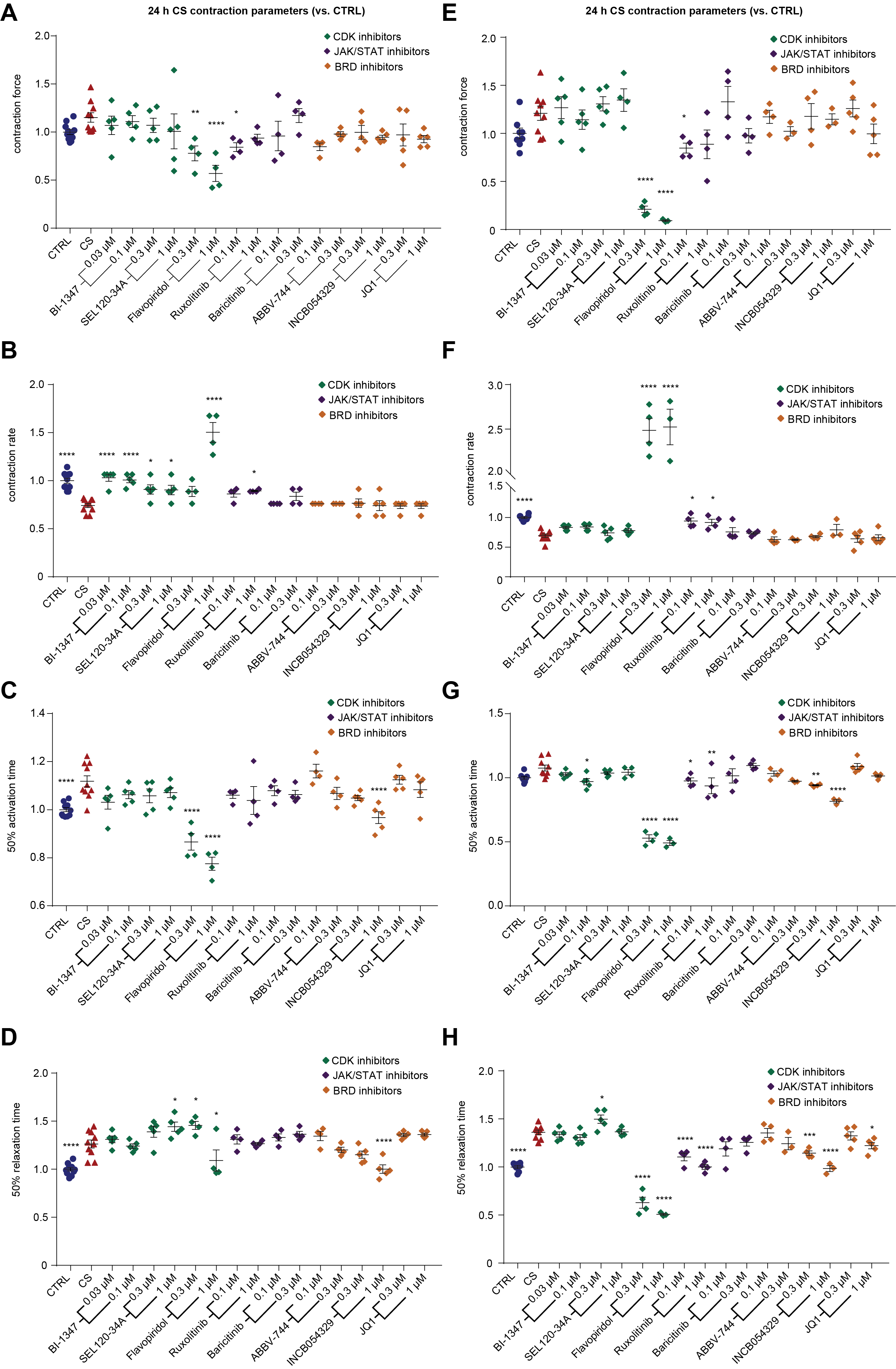

**Supplementary Figure S10: Screening for compounds that prevent diastolic dysfunction in hCO treated with cardiac cytokine storm.**

A) Contraction force in hCO derived from hPSCs cardiac cells - HES3, Endothelial cells - RM3.5.
B) Contraction rate in hCO derived from hPSCs cardiac cells - HES3, Endothelial cells - RM3.5.
C) Time to 50% activation in hCO derived from hPSCs cardiac cells - HES3, Endothelial cells - RM3.5.
D) Time to 50% relaxation in hCO derived from hPSCs cardiac cells - HES3, Endothelial cells - RM3.5.
E) Contraction force in hCO derived from hPSCs cardiac cells - AA, Endothelial cells – CC.
F) Contraction rate in hCO derived from hPSCs cardiac cells - AA, Endothelial cells – CC.
G) Time to 50% activation in hCO derived from hPSCs cardiac cells - AA, Endothelial cells – CC.
H) Time to 50% relaxation in hCO derived from hPSCs cardiac cells - AA, Endothelial cells – CC.

hCOs were concurrently treated with the CS and compounds (all compounds in the presence of CS conditions), and then functionally assessed at 24 h. (a-d) n = 4-11 and (e-f) n = 3-9 hCOs per condition from 1 experiment. *p<0.05, ** p<0.01, *** p<0.001, **** p<0.0001, using one-way ANOVA with Dunnett's multiple comparisons test compared to the CS.

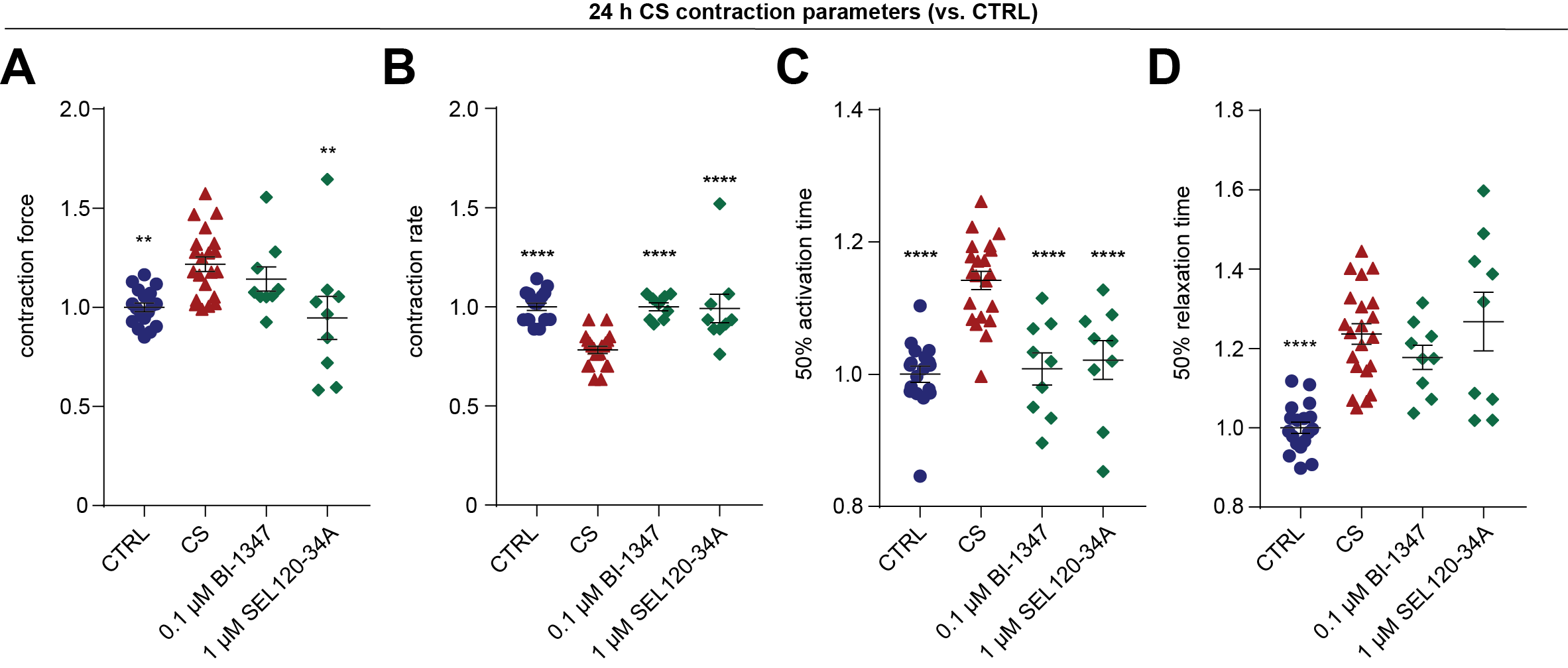

**Supplementary Figure S11: CDK8 inhibitors specifically restore hCO rate and activation time effects of the cardiac cytokine storm, but not diastolic dysfunction.**

A) Contraction force.
B) Contraction rate.
C) Time from 50% activation to peak.
D) Time to 50% relaxation.

hCOs were concurrently treated with the CS and CDK8-STAT1 S727 inhibitors (all compounds in presence of CS), and then functionally assessed at 24 h. n = 9-21 hCOs per condition from 2 experiments, hPSCs cardiac cells - HES3, Endothelial cells- RM3.5. ** p<0.01, **** p<0.0001, using one-way ANOVA with Dunnett's multiple comparisons test compared to CS.

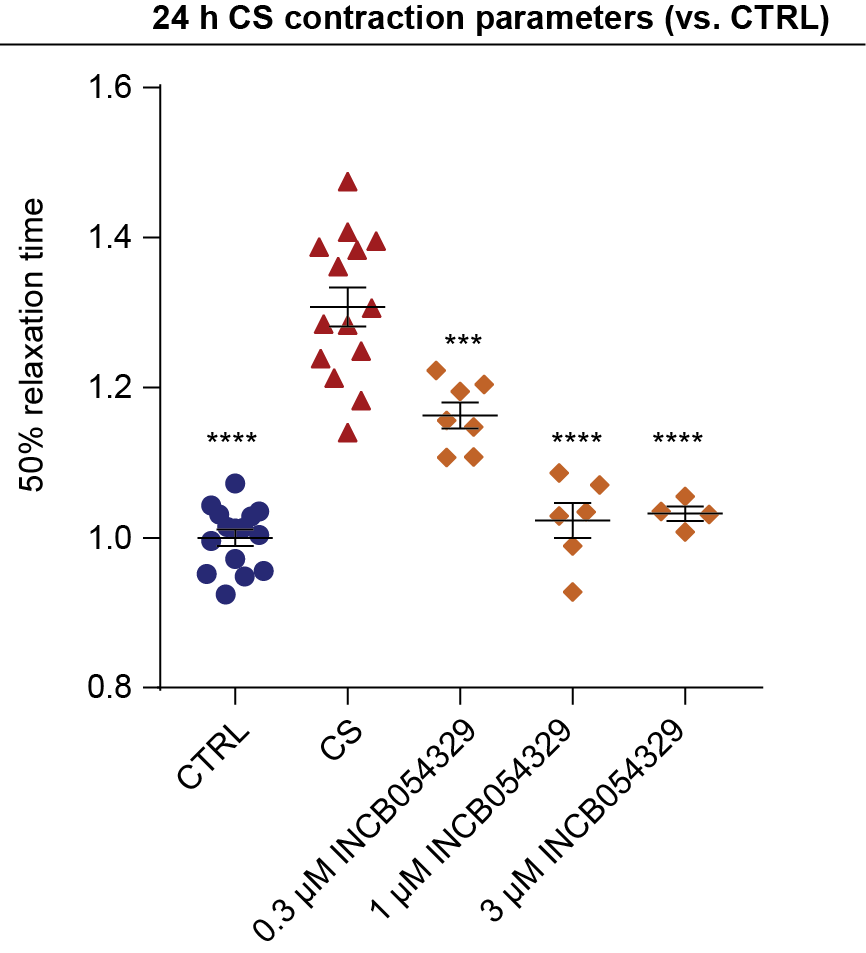

**Supplementary Figure S12: Validation of INCB054329 protection against diastolic dysfunction induced by the cardiac cytokine storm in another hPSC line.**

Time to 50% relaxation. hCOs were concurrently treated with the CS and INCB054329 (all compounds in presence of CS), and then functionally assessed at 24 h. n = 4-12 hCOs per condition from 1-2 experiments, hPSCs cardiac cells - AA, Endothelial cells- CC. *** p<0.001, **** p<0.0001, using one-way ANOVA with Dunnett's multiple comparisons test compared to CS.

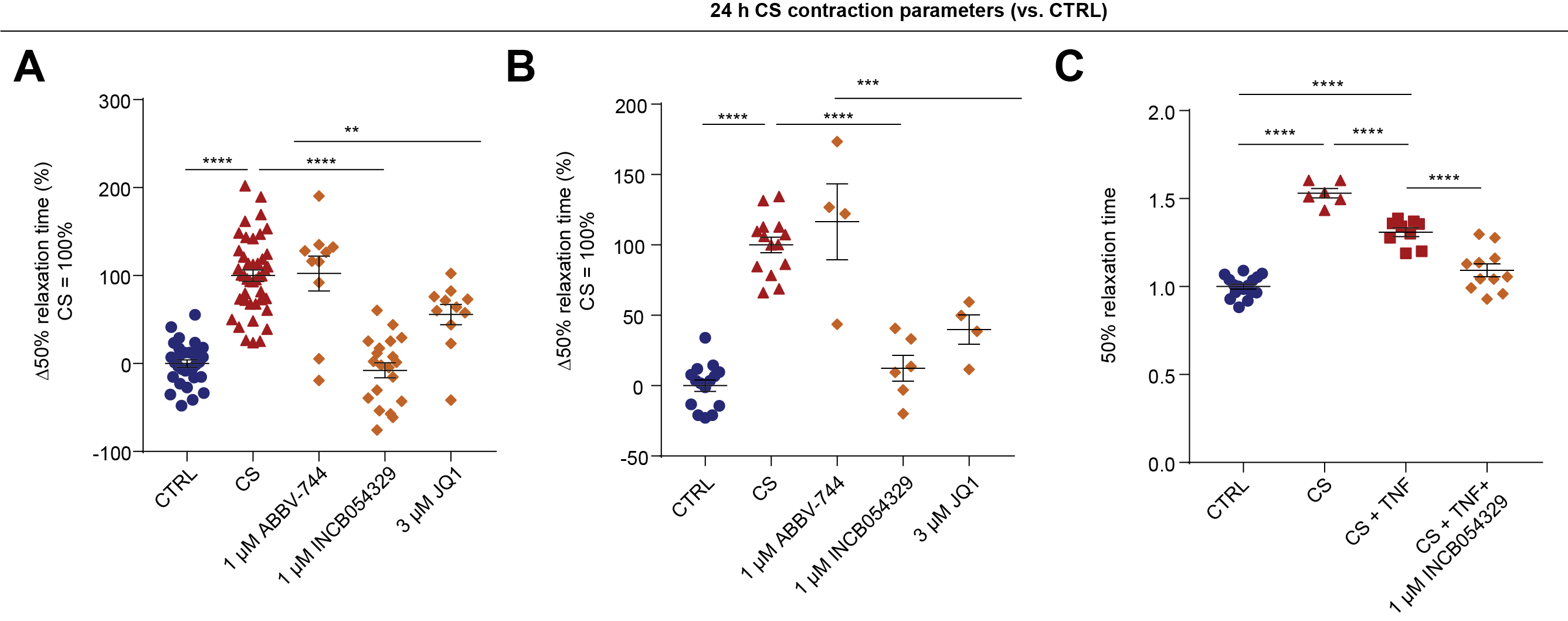

**Supplementary Figure S13: Prevention of diastolic dysfunction in hCO by BRD4 inhibitors**

A) BRD inhibitors prevent CS induced diastolic dysfunction presented as change relative to increased relaxation time. n = 8-43 hCOs per condition from 2-4 experiment, hPSCs cardiac cells- HES3, Endothelial cells- RM3.5. **p<0.01, **** p<0.0001, using one-way ANOVA with Tukey’s multiple comparisons test compared to CS.
B) Validation of results in an additional hPSC line. BRD inhibitors prevent CS induced diastolic dysfunction presented as change relative to increased relaxation time. n = 14-15 for CTRL and CS conditions and 4-6 hCOs per BRD inhibitor from 1-2 experiments, hPSC cardiac cells- AA, Endothelial cells- CC. *** p<0.001, **** p<0.0001, using one-way ANOVA with Tukey’s multiple comparisons test compared to cytokine storm - CS.
C) Assessment of INCB054329 efficacy in conditions with CS and TNF. n = 6-16 hCOs per condition from 1-2 experiments. hPSCs cardiac cells- HES3, Endothelial cells- RM3.5. ** p<0.01, **** p<0.0001, using a one-way ANOVA with Tukey’s multiple comparisons test.

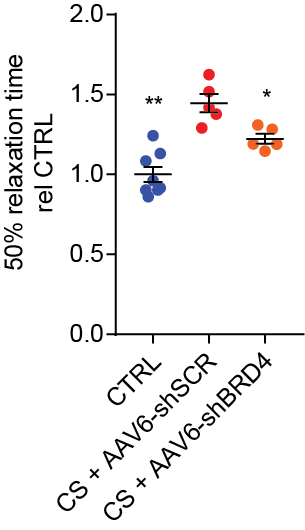

**Supplementary Figure S14: Knockdown of BRD4 prevents diastolic dysfunction.**

BRD4 knockdown prevents CS induced diastolic dysfunction presented as normalized relaxation time. n = 5-8 hCOs per condition from 1 experiment, hPSCs cardiac cells- HES3. *p<0.05, ** p<0.01, using one-way ANOVA with Dunnett’s multiple comparisons test relative to CS + AAV6-shSCR (scramble control).

**Supplementary Figure S15: Ingenuity Pathway Analysis^TM^ networks in hCO and *in vivo*.**

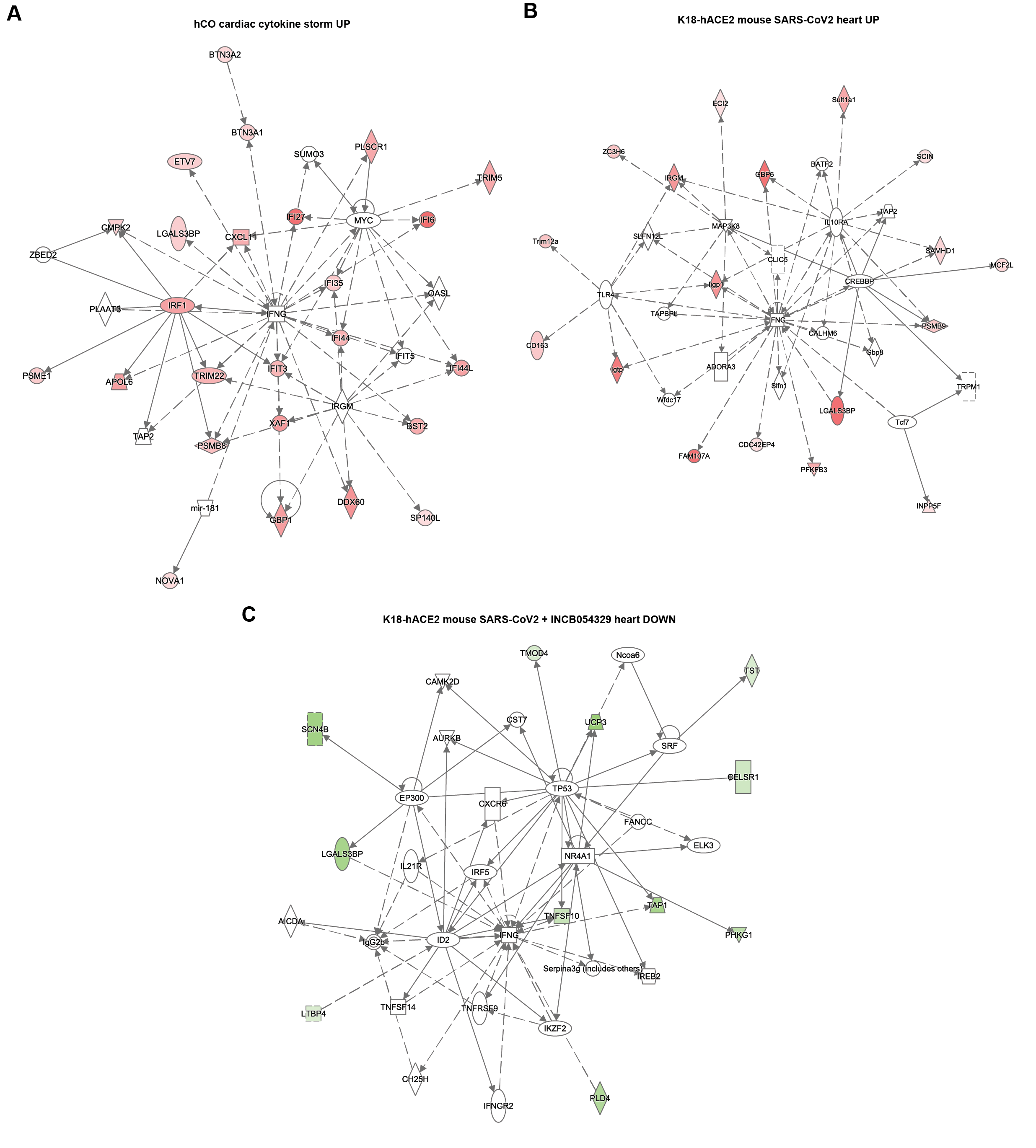

A) Induced genes (logFC > 0.5, FDR < 0.05) in a network centred around IFN-γ induced by CS in hCO.
B) Induced genes (logFC > 0.5, FDR < 0.05) in a network centred around IFN-γ induced by SARS-CoV-2 infection in K18-hACE2 mice.
C) Genes decreased (logFC < -0.5, FDR < 0.05) in a network centred around IFN-γ following treatment with INCB054329 in SARS-CoV-2 infected K18-hACE2 mice.

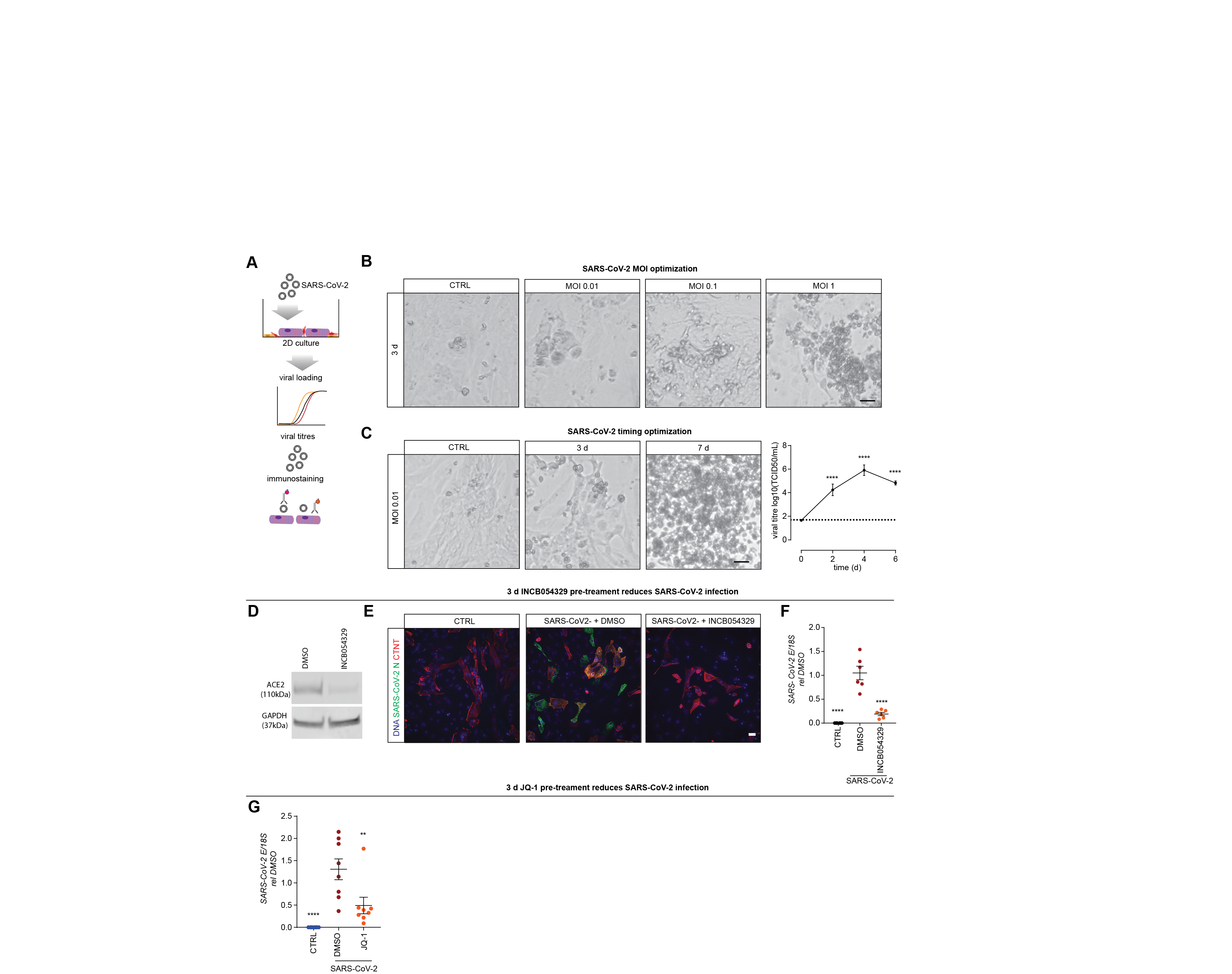

**Supplementary Figure S16: Pre-treatment with INCB054329 prevents SARS-CoV-2 infection of hPSC-cardiac cells.**

A) Schematic of the experiments.
B) Optimization of loading with increasing cell death of 2D hPSC-cardiac cells with increasing SARS-CoV-2 infection.
C) Infection at low MOI (0.01) results in viral replication and eventually death following SARS-CoV-2 infection of 2D hPSC-cardiac cells. n = 6 from 3 experiments (hPSC cardiac cells –pooled from 2 hPSC lines).
D) ACE2 expression in 2D cultured hPSC-cardiac cells pre-treated with 1 µM INCB054329 for 3 days. hPSC cardiac cells – HES3.
E) Immunostaining of cardiomyocytes (CTNT) and SARS-CoV-2 reveals that 1 µM INCB054329 reduces viral loading.
F) Pre-treatment with 1 µM INCB054329 for 3 days reduces SARS-CoV-2 infection. E-gene expression in 2D cultured hPSC-cardiac cells 3 days after infection. n = 6 from 2 experiments hPSC cardiac cells – AA).
G) Pre-treatment with 3 µM JQ-1 for 3 days reduces SARS-CoV-2 infection. E-gene expression in 2D cultured hPSC-cardiac cells 3 days after infection. n = 8 from 2 experiments hPSC cardiac cells – HES3 and AA).

All scale bars = 20 µm. TCID50 - Fifty-percent tissue culture infective dose. Data presented as mean ± standard error of the mean. **** p<0.0001, using one-way ANOVA with Dunnett's multiple comparisons test (C - compared day 0 and F,G – compared to DMSO).

**Supplementary Figure S17: Supporting data for the repression of ACE2 and SARS-CoV-2 infection by compounds targeting the BD2 domain.**

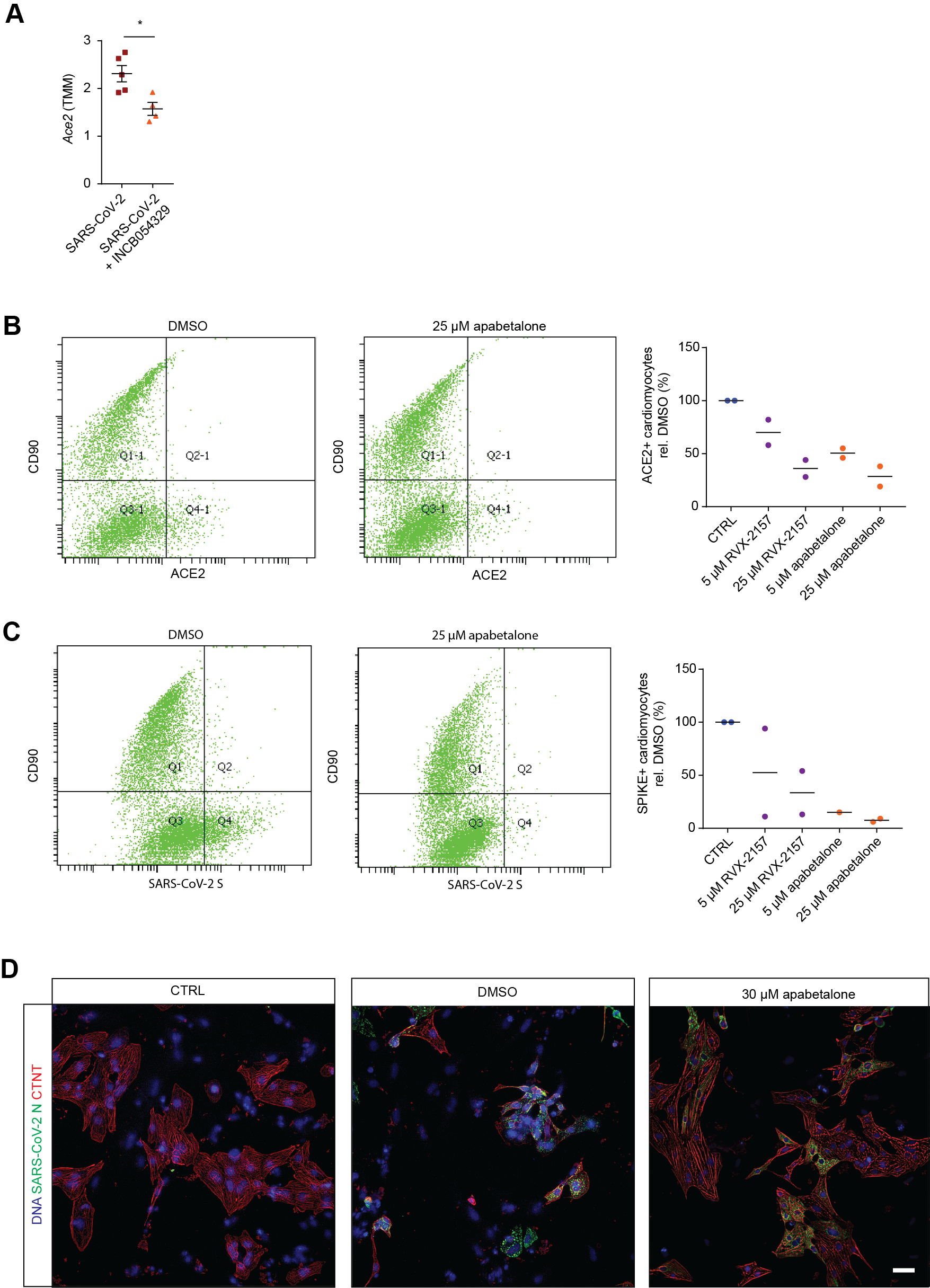

A) INCB054329 decreases endogenous mAce2 expression in hearts in vivo. n = 4-5 mice. * p < 0.05 using Mann-Whitney.
B) Flow cytometry analysis of ACE2 on cardiomyocytes (CD90 negative) and CD90 positive stromal cells. Analysis was performed in 2 different cells lines (HES3 and AA) in separate experiments.
C) Flow cytometry analysis of spike protein binding on cardiomyocytes (CD90 negative) and CD90 positive stromal cells. Analysis was performed in 2 different cells lines (HES3 and AA) in separate experiments.
D) Immunostaining of cardiomyocytes (CTNT) and SARS-CoV-2 reveals that BET bromodomain inhibition with 30 µM apabetalone preserves cardiomyocyte structures and reduces viral loading. Scale bar = 20 µm.

**Supplementary Figure S18: Induction of arrhythmias under cardiac cytokine storm conditions and protection against arrhythmias with INCB054329.**

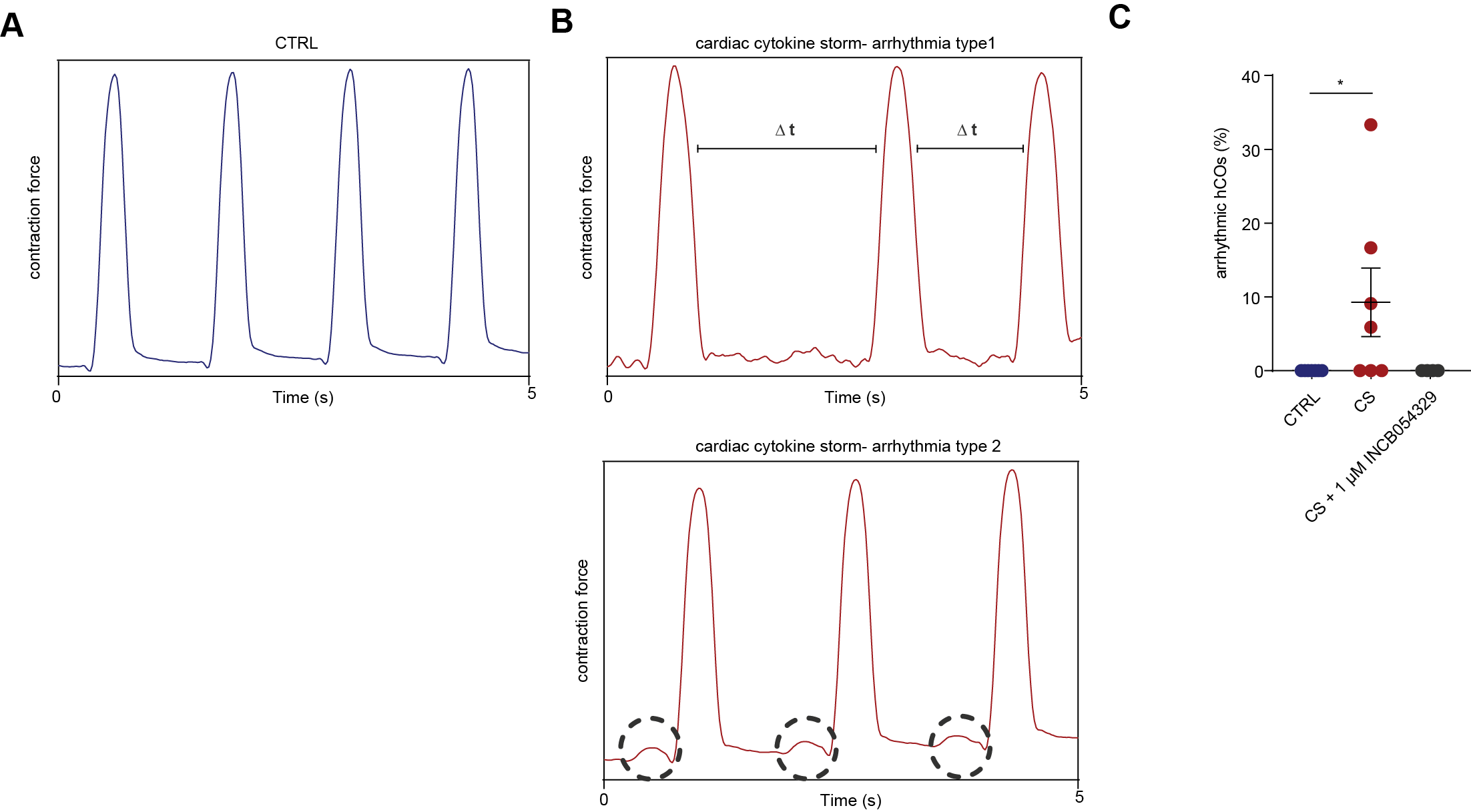

A) Representative force trace of a CTRL hCO.
B) Representative force trace of different types of arrhythmias in hCO under CS conditions.
C) Arrhythmic events in hCO per experiment.

hIPSCs cardiac cells- HES3, Endothelial cells- RM3.5. n = 4-7 experiments. Corrected p-value calculated using one-way ANOVA with Kruskal-Wallis comparisons test to CTRL.

**Supplementary Table S1: Top 10 KEGG pathways for RNA-seq analyses.**

| **Top 10 KEGG Pathways** | **Gene Number** | **Adjusted P-value** | **Genes** |
| --- | --- | --- | --- |
| ***UP in hCO cardiomyocytes*** |  |  |  |
| Epstein-Barr virus infection | 17 | 1.72E-13 | DDX58;STAT1;STAT2;STAT3;HLA-B;HLA-C;TAP1;EIF2AK2;ISG15;HLA-A;HLA-F;HLA-E;TAPBP;CXCL10;OAS2;OAS3;B2M |
| Antigen processing and presentation | 10 | 9.55E-10 | CD74;HLA-B;PSME1;TAP1;HLA-C;HLA-A;HLA-F;B2M;TAPBP;HLA-E |
| Herpes simplex virus 1 infection | 19 | 1.46E-09 | CD74;SP100;DDX58;STAT1;STAT2;HLA-B;HLA-C;TAP1;EIF2AK2;HLA-A;HLA-F;HLA-E;TAPBP;BST2;IFIH1;OAS2;OAS3;CCL2;B2M |
| Influenza A | 12 | 6.65E-09 | IFIH1;CXCL10;RSAD2;DDX58;OAS2;STAT1;OAS3;STAT2;MX1;EIF2AK2;CCL2;ADAR |
| Hepatitis C | 11 | 2.72E-08 | CXCL10;RSAD2;DDX58;OAS2;STAT1;OAS3;STAT2;MX1;STAT3;EIF2AK2;IFIT1 |
| Measles | 10 | 1.07E-07 | IFIH1;DDX58;OAS2;STAT1;OAS3;STAT2;MX1;STAT3;EIF2AK2;ADAR |
| Human immunodeficiency virus 1 infection | 11 | 5.13E-07 | BST2;TRIM5;HLA-B;HLA-C;TAP1;HLA-A;HLA-F;SAMHD1;B2M;HLA-E;TAPBP |
| Viral myocarditis | 7 | 5.36E-07 | CAV1;HLA-B;HLA-C;HLA-A;HLA-F;HLA-E;MYH7 |
| Human cytomegalovirus infection | 10 | 7.17E-06 | STAT3;HLA-B;HLA-C;TAP1;CCL2;HLA-A;HLA-F;B2M;HLA-E;TAPBP |
| Kaposi sarcoma-associated herpesvirus infection | 9 | 1.19E-05 | STAT1;STAT2;STAT3;HLA-B;HLA-C;EIF2AK2;HLA-A;HLA-F;HLA-E |
| ***UP in hCO fibroblasts*** |  |  |  |
| Epstein-Barr virus infection | 27 | 8.39E-17 | TNFAIP3;ICAM1;RELB;CCND3;B2M;LYN;STAT1;DDX58;STAT2;HLA-B;HLA-C;EIF2AK2;TAP1;ISG15;HLA-A;HLA-F;NFKB1;HLA-E;TAPBP;NFKBIA;CXCL10;OAS1;TRAF3;OAS2;OAS3;HLA-DRA;HLA-DRB1 |
| NOD-like receptor signaling pathway | 25 | 3.02E-16 | CXCL8;TNFAIP3;CXCL1;ANTXR2;IFI16;CCL5;NAMPT;CASP4;CCL2;GBP2;GBP1;GBP4;GBP3;GBP5;RIPK2;STAT1;STAT2;NFKB1;NFKBIA;OAS1;TRAF3;OAS2;OAS3;RBCK1;BIRC3 |
| Influenza A | 24 | 9.88E-16 | CXCL8;RSAD2;DDX58;STAT1;STAT2;MX1;EIF2AK2;ADAR;NFKB1;PML;ICAM1;NFKBIA;IFIH1;CXCL10;OAS1;OAS2;OAS3;CCL5;TNFSF10;TRIM25;HLA-DRA;CCL2;JAK2;HLA-DRB1 |
| TNF signaling pathway | 19 | 3.58E-14 | VCAM1;CSF1;MLKL;IL15;TNFAIP3;VEGFC;CXCL1;CFLAR;NFKB1;ICAM1;NFKBIA;CXCL10;CASP10;TRAF3;IRF1;CCL5;CCL2;MAP3K8;BIRC3 |
| Herpes simplex virus 1 infection | 30 | 3.86E-10 | SP100;C3;IFIH1;CCL5;CCL2;JAK2;B2M;CD74;STAT1;DDX58;STAT2;HLA-B;HLA-C;EIF2AK2;TAP1;HLA-A;HLA-F;NFKB1;PML;HLA-E;TAPBP;BST2;NFKBIA;OAS1;TRAF3;OAS2;OAS3;HLA-DRA;HLA-DRB1;BIRC3 |
| Kaposi sarcoma-associated herpesvirus infection | 19 | 4.07E-10 | LYN;CXCL8;STAT1;STAT2;HLA-B;HLA-C;EIF2AK2;CXCL1;HLA-A;HLA-F;HIF1A;NFKB1;ICAM1;HLA-E;C3;NFKBIA;TRAF3;IL6ST;JAK2 |
| NF-kappa B signaling pathway | 14 | 1.31E-09 | LYN;CXCL8;VCAM1;DDX58;TNFAIP3;CFLAR;NFKB1;TNFSF13B;ICAM1;RELB;NFKBIA;TRAF3;TRIM25;BIRC3 |
| Measles | 16 | 1.94E-09 | DDX58;STAT1;STAT2;MX1;EIF2AK2;TNFAIP3;CBLB;ADAR;NFKB1;NFKBIA;IFIH1;CCND3;OAS1;TRAF3;OAS2;OAS3 |
| Antigen processing and presentation | 12 | 1.29E-08 | CD74;HLA-B;TAP1;HLA-C;HLA-DRA;HLA-A;HLA-F;B2M;CTSS;HLA-DRB1;TAPBP;HLA-E |
| RIG-I-like receptor signaling pathway | 11 | 5.58E-08 | IFIH1;NFKBIA;CXCL10;CYLD;CXCL8;CASP10;DDX58;TRAF3;TRIM25;ISG15;NFKB1 |
| ***UP in K18-hACE2 SARS-CoV-2 infected mouse lungs*** |  |  |  |
| Epstein-Barr virus infection | 31 | 3.02E-14 | H2-T23;CDKN1A;H2-T22;PSMD13;H2-Q6;H2-K1;H2-Q7;H2-Q4;TNFAIP3;RELB;PSMD4;MYC;H2-DMB1;BAK1;IKBKE;IFNAR2;STAT1;DDX58;STAT2;ADRM1;TAP1;ISG15;RUNX3;CXCL10;IL6;CCNE1;MDM2;IRF7;H2-D1;IRF9;TLR2 |
| Influenza A | 19 | 2.82E-07 | IFNAR2;CCL12;RSAD2;DDX58;STAT1;STAT2;TNFRSF10B;ADAR;SOCS3;CXCL10;IL6;H2-DMB1;CCL5;IRF7;CCL2;IKBKE;IRF9;HSPA1B;HSPA1A |
| Cytosolic DNA-sensing pathway | 12 | 3.00E-07 | ZBP1;CXCL10;IL6;RIPK3;DDX58;CCL5;TREX1;CCL4;IRF7;CGAS;ADAR;IKBKE |
| Kaposi sarcoma-associated herpesvirus infection | 21 | 3.23E-07 | IFNAR2;H2-T23;CDKN1A;H2-T22;STAT1;STAT2;H2-Q6;H2-K1;H2-Q7;H2-Q4;C3;RCAN1;IL6;MYC;MAPKAPK2;IRF7;BAK1;CCR5;IKBKE;H2-D1;IRF9 |
| Phagosome | 19 | 3.58E-07 | H2-T23;H2-T22;C1RA;H2-Q6;H2-K1;NCF4;H2-Q7;H2-Q4;TAP1;THBS1;FCGR1;C3;TUBA1C;TUBB6;FCGR4;H2-DMB1;ITGA5;H2-D1;TLR2 |
| DNA replication | 9 | 1.10E-06 | RFC5;FEN1;MCM7;LIG1;MCM3;MCM5;MCM6;POLE;MCM2 |
| Complement and coagulation cascades | 13 | 1.10E-06 | C1QB;C1QA;C1RA;SERPINE1;F3;C2;C3;PLAU;C3AR1;BDKRB1;SERPING1;CFB;C1QC |
| Antigen processing and presentation | 13 | 1.27E-06 | H2-T23;H2-T22;H2-Q6;H2-K1;H2-Q7;H2-Q4;TAP1;H2-DMB1;PSME1;H2-D1;HSPA1B;HSPA1A;LGMN |
| Measles | 16 | 1.46E-06 | IFNAR2;DDX58;STAT1;STAT2;TNFAIP3;ADAR;IL6;CCNE1;IL2RB;IRF7;BAK1;IKBKE;IRF9;HSPA1B;TLR2;HSPA1A |
| TNF signaling pathway | 14 | 1.60E-06 | CCL12;CSF1;MLKL;RIPK3;LIF;TNFAIP3;SOCS3;CXCL10;IL6;MMP14;CCL5;BCL3;CCL2;GM5431 |
| ***UP in K18-hACE2 SARS-CoV-2 infected mouse hearts*** |  |  |  |
| Antigen processing and presentation | 11 | 3.37E-06 | H2-T23;H2-T22;PSME2B;H2-Q6;H2-K1;H2-Q7;H2-Q4;TAP1;PSME2;B2M;H2-D1 |
| Epstein-Barr virus infection | 16 | 3.37E-06 | H2-T23;H2-T22;GADD45B;STAT1;DDX58;H2-Q6;H2-K1;H2-Q7;H2-Q4;EIF2AK2;TAP1;PIK3R1;IRF7;B2M;H2-D1;IRF9 |
| Human immunodeficiency virus 1 infection | 16 | 3.85E-06 | TRIM12C;H2-T23;APOBEC3;H2-T22;H2-Q6;H2-K1;H2-Q7;H2-Q4;TAP1;PIK3R1;SAMHD1;BST2;GNB3;B2M;H2-D1;BCL2L1 |
| Herpes simplex virus 1 infection | 21 | 6.82E-06 | H2-T23;H2-T22;STAT1;DDX58;H2-Q6;H2-K1;H2-Q7;H2-Q4;EIF2AK2;TAP1;PIK3R1;C3;BST2;IFIH1;SOCS3;EIF4EBP1;IRF7;B2M;H2-D1;IRF9;BCL2L1 |
| Kaposi sarcoma-associated herpesvirus infection | 14 | 2.12E-05 | H2-T23;H2-T22;STAT1;H2-Q6;H2-K1;H2-Q7;H2-Q4;EIF2AK2;PIK3R1;C3;IRF7;GNB3;H2-D1;IRF9 |
| Cellular senescence | 13 | 2.12E-05 | H2-T23;H2-T22;GADD45B;H2-Q6;SERPINE1;H2-K1;H2-Q7;H2-Q4;PIK3R1;FOXO3;EIF4EBP1;SQSTM1;H2-D1 |
| Viral carcinogenesis | 13 | 1.95E-04 | H2-T23;H2-T22;H2-Q6;H2-K1;H2-Q7;H2-Q4;EIF2AK2;PIK3R1;C3;SCIN;IRF7;H2-D1;IRF9 |
| Viral myocarditis | 8 | 2.81E-04 | H2-T23;H2-T22;H2-Q6;H2-K1;H2-Q7;H2-Q4;H2-D1;MYH7 |
| Drug metabolism | 9 | 2.81E-04 | ADH1;DPYD;MGST1;GSTT2;AOX1;FMO1;FMO2;XDH;FMO5 |
| Allograft rejection | 7 | 2.81E-04 | H2-T23;H2-T22;H2-Q6;H2-K1;H2-Q7;H2-Q4;H2-D1 |
| ***DOWN in INCB054329 treated K18-hACE2 SARS-CoV-2 infected mouse hearts*** |  |  |  |
| Epstein-Barr virus infection | 14 | 3.00E-10 | H2-T22;H2-EB1;STAT1;H2-K1;TAP1;H2-AA;ICAM1;RELB;CDK6;IRF7;PLCG2;B2M;CD44;H2-AB1 |
| Antigen processing and presentation | 10 | 6.66E-10 | CD74;H2-T22;H2-EB1;H2-K1;PSME1;TAP1;PSME2;B2M;H2-AA;H2-AB1 |
| Toxoplasmosis | 7 | 2.80E-05 | H2-EB1;LAMB3;STAT1;IGTP;IRGM1;H2-AA;H2-AB1 |
| African trypanosomiasis | 5 | 3.05E-05 | HBA-A2;HBB-BT;HBA-A1;HBB-BS;ICAM1 |
| Viral myocarditis | 6 | 6.49E-05 | H2-T22;H2-EB1;H2-K1;H2-AA;ICAM1;H2-AB1 |
| Malaria | 5 | 6.49E-05 | HBA-A2;HBB-BT;HBA-A1;HBB-BS;ICAM1 |
| Herpes simplex virus 1 infection | 11 | 8.07E-05 | CD74;H2-T22;H2-EB1;STAT1;H2-K1;IRF7;TAP1;CFP;B2M;H2-AA;H2-AB1 |
| Staphylococcus aureus infection | 6 | 8.07E-05 | C4B;H2-EB1;PTAFR;H2-AA;ICAM1;H2-AB1 |
| Kaposi sarcoma-associated herpesvirus infection | 8 | 9.75E-05 | H2-T22;CDK6;STAT1;H2-K1;IRF7;PLCG2;GNB3;ICAM1 |
| Allograft rejection | 5 | 1.33E-04 | H2-T22;H2-EB1;H2-K1;H2-AA;H2-AB1 |

**Supplementary Table S2: Top predicted ENCODE transcriptional regulators in RNA-seq analyses.**

| **Transcriptional regulator** | **Gene Number** | **Adjusted**  **P-value** |
| --- | --- | --- |
| ***UP in hCO cardiomyocytes*** |  |  |
| STAT2 K562 hg19 | 57 | 4.14E-74 |
| STAT1 K562 hg19 | 57 | 6.77E-68 |
| STAT1 HeLa-S3 hg19 | 35 | 4.41E-17 |
| IRF1 K562 hg19 | 66 | 1.15E-13 |
| IKZF1 GM12878 hg19 | 36 | 2.84E-08 |
| STAT3 HeLa-S3 hg19 | 35 | 7.00E-08 |
| EP300 CH12.LX mm9 | 35 | 8.54E-08 |
| PRDM1 HeLa-S3 hg19 | 31 | 1.21E-07 |
| ***UP in hCO fibroblasts*** |  |  |
| STAT2 K562 hg19 | 76 | 2.26E-76 |
| STAT1 K562 hg19 | 79 | 3.27E-72 |
| STAT1 HeLa-S3 hg19 | 62 | 7.09E-23 |
| IKZF1 GM12878 hg19 | 89 | 2.97E-22 |
| EP300 CH12.LX mm9 | 69 | 7.03E-11 |
| RELA GM12878 hg19 | 40 | 1.21E-10 |
| PRDM1 HeLa-S3 hg19 | 59 | 4.91E-10 |
| STAT3 HeLa-S3 hg19 | 66 | 6.80E-10 |
| ***UP in K18-hACE2 SARS-CoV-2 infected mouse lungs*** |  |  |
| EP300 CH12.LX mm9 | 111 | 1.71E-19 |
| STAT2 K562 hg19 | 31 | 6.73E-12 |
| STAT1 K562 hg19 | 32 | 5.63E-10 |
| ETS1 CH12.LX mm9 | 81 | 9.03E-07 |
| JUND CH12.LX mm9 | 63 | 4.56E-06 |
| JUN CH12.LX mm9 | 64 | 1.84E-04 |
| ETS1 MEL cell line mm9 | 67 | 0.001847007 |
| STAT1 HeLa-S3 hg19 | 40 | 0.002081127 |
| ***UP in K18-hACE2 SARS-CoV-2 infected mouse hearts*** |  |  |
| STAT2 K562 hg19 | 29 | 5.54E-16 |
| STAT1 K562 hg19 | 28 | 1.56E-12 |
| ETS1 MEL cell line mm9 | 58 | 2.38E-08 |
| ETS1 CH12.LX mm9 | 49 | 5.79E-04 |
| STAT1 HeLa-S3 hg19 | 29 | 0.001169147 |
| TCF12 myocyte mm9 | 32 | 0.007490599 |
| POLR2A heart mm9 | 45 | 0.007490599 |
| EP300 CH12.LX mm9 | 43 | 0.026728037 |
| ***DOWN in INCB054329 treated K18-hACE2 SARS-CoV-2 infected mouse hearts*** |  |  |
| EP300 CH12.LX mm9 | 24 | 3.25E-04 |
| STAT2 K562 hg19 | 8 | 0.004464122 |
| STAT1 K562 hg19 | 7 | 0.109330724 |

**Supplementary Table S3: IPA top predicted upstream regulators in RNA-seq analyses.**

| **IPA top 10 predicted upstream regulators** | **Activation Z-score** | **P-value of overlap** |
| --- | --- | --- |
| ***All FDR < 0.05 in K18-hACE2 SARS-CoV-2 infected mouse lungs*** |  |  |
| lipopolysaccharide | 9.449 | 9.94E-44 |
| IFNG | 8.651 | 2.18E-49 |
| poly rI:rC-RNA | 7.831 | 2.35E-46 |
| TNF | 7.257 | 8.64E-30 |
| tetradecanoylphorbol acetate | 7.008 | 9.34E-21 |
| Interferon alpha | 6.852 | 2.47E-39 |
| STAT1 | 6.616 | 6.73E-43 |
| IL1B | 6.521 | 1.09E-30 |
| IRF7 | 6.221 | 2.57E-34 |
| PDGF BB | 6.018 | 1.08E-23 |
| ***All FDR < 0.05 in K18-hACE2 SARS-CoV-2 infected mouse hearts*** |  |  |
| IRF7 | 5.429 | 4.55E-14 |
| poly rI:rC-RNA | 5.35 | 6.39E-07 |
| STAT1 | 5.164 | 2.97E-13 |
| FOXO3 | 5.008 | 5.93E-08 |
| Ifnar | 4.876 | 1.48E-13 |
| IRF3 | 4.848 | 1.96E-07 |
| IFNG | 4.791 | 7.45E-14 |
| Interferon alpha | 4.56 | 3.87E-11 |
| pirinixic acid | 4.157 | 1.01E-13 |
| IFNA2 | 4.036 | 2.04E-11 |
| ***All FDR < 0.05 in INCB054329 treated K18-hACE2 SARS-CoV-2 infected mouse hearts*** |  |  |
| lipopolysaccharide | -4.378 | 1.40E-09 |
| IFNG | -4.329 | 8.10E-10 |
| poly rI:rC-RNA | -4.116 | 6.03E-09 |
| TNF | -3.39 | 4.83E-06 |
| Interferon alpha | -3.382 | 1.74E-05 |
| STAT1 | -3.263 | 7.84E-10 |
| NFkB (complex) | -3.222 | 2.00E-03 |
| IL27 | -3.086 | 2.49E-07 |
| IRF3 | -3.076 | 7.58E-07 |
| tretinoin | -3.075 | 9.99E-03 |

**VIDEO LEGENDS**

**Supplementary Video 1:** hCO cultured under CTRL conditions. The video was taken over a period of 10 seconds and is displayed in real time (50 frames/s).

**Supplementary Video 2:** hCO cultured under CS conditions. The video was taken over a period of 10 seconds and is displayed in real time (50 frames/s).

**Supplementary Video 3:** hCO cultured under CTRL conditions paced at 1 Hz. The video was taken over a period of 5 seconds and is displayed in real time (50 frames/s).

**Supplementary Video 4:** hCO cultured under CS paced at 1 Hz. The video was taken over a period of 5 seconds and is displayed in real time (50 frames/s).

**Supplementary Video 5:** hCO cultured under CS conditions treated with 1µM INCB054329. The video was taken over a period of 10 seconds and is displayed in real time (50 frames/s).

**SUPPLEMENTARY REFERENCES**

1.	R. J. Mills *et al.*, Functional screening in human cardiac organoids reveals a metabolic mechanism for cardiomyocyte cell cycle arrest. *Proceedings of the National Academy of Sciences* **114**, E8372-E8381 (2017).

2.	G. A. Quaife-Ryan *et al.*, Multicellular Transcriptional Analysis of Mammalian Heart Regeneration. *Circulation* **136**, 1123-1139 (2017).

3.	N. R. Tucker *et al.*, Transcriptional and Cellular Diversity of the Human Heart. *Circulation* **0**, (2020).

4.	R. Gilsbach *et al.*, Distinct epigenetic programs regulate cardiac myocyte development and disease in the human heart in vivo. *Nature communications* **9**, 391 (2018).
